# How the TREX-2 complex associates with the nuclear pore

**DOI:** 10.64898/2026.02.20.707084

**Authors:** Agnieszka Obarska-Kosinska, Ying Zhu, Katharina Geißler, Remus R E Rosenkranz, Nanako Yokoyama, Jan Philipp Kreysing, Huaipeng Xing, Desislava Glushkova, Marta Anna Kubańska, Stefanie Böhm, Hans-Georg Kräusslich, Beata Turoňová, Fan Liu, Martin Beck

## Abstract

Nuclear pore complexes (NPCs) control nucleocytoplasmic transport in eukaryotes, yet their architecture remains incompletely understood. Here we report a substantially extended structure of the human NPC, obtained by combining cryo-electron tomography, crosslinking mass spectrometry, and AI-assisted integrative modeling. We resolve the molecular arrangement of TPR, NUP153, NUP50 and ZC3HC1, and reveal that five additional proteins — TMEM209, SMPD4, GANP, Centrin-2 and ENY2 — are incorporated into the NPC. Unexpectedly, GANP, Centrin-2 and ENY2, core members of the TREX-2 mRNA export complex, are built into the nuclear ring. This finding establishes TREX-2 not as a transiently associated factor, but as an integral NPC module, positioning it opposite of the cytoplasmic NUP214 mRNA export platform. Together, our results redefine the molecular composition of the inner ring, nuclear ring and nuclear basket. They suggest a direct structural basis that couples TREX-2-mediated mRNP remodeling to NPC-facilitated transport.

## Introduction

Nuclear pore complexes (NPCs) facilitate nucleocytoplasmic exchange, and are crucial for the maintenance of genetic stability in eukaryotic organisms.^1,2^ They are large ∼120 MDa complexes made up of multiple copies of about 30 different nucleoporins (NUPs).^3,4^ These nucleoporins can be broadly grouped into scaffold NUPs, which form the cylindrical core architecture, and FG-NUPs, which line the central transport channel and mediate interactions with cargo complexes. Through this organization, NPCs regulate both protein transport and mRNA export, two fundamental processes for eukaryotes^5^.

mRNA export requires the coordinated activity of several nuclear export factors and NPC components.^6^ In humans, the NUP214 subcomplex associates with the cytoplasmic face of the NPC and recruits DDX19, a helicase that is thought to remodel mRNPs upon exit from the central channel^3,4^. Furthermore, the nucleoplasmic mRNA export complex TREX-2, comprised of GANP, PCID2, ENY2, DSS1 and Centrin-2/3 (CETN2/3)^7–9^, localizes to the nuclear envelope (NE) and associates with the NPC. This depends on the nuclear basket NUP translocated promoter region (TPR).^10,11^ GANP acts as a scaffold for TREX-2^12^, and depletion of either GANP or ENY2 leads to mRNA retention phenotypes similar to those observed upon loss of nuclear basket NUPs, suggesting that TREX-2 anchoring to the NPC is required for efficient mRNA export.^10^ Despite this functional link, the structural basis of TREX-2 recruitment and its precise positioning at the NPC remain unclear.

The overall architecture of the NPC scaffold has been delineated by decades of biochemical and integrative structural studies^3,13–17^, revealing a tripartite arrangement comprising the cytoplasmic ring (CR), inner ring (IR) and nuclear ring (NR). The CR and NR are scaffolded by so-called Y-complexes and inner ring core modules build the IR ^18–22^. While these analyses have provided composite models of the scaffold at increasing molecular precision, several regions of density in cryo-EM maps remain unassigned. The structural arrangement of multifunctional scaffold NUPs that engage in multiple different, often mutually exclusive interactions, and form compositionally distinct variants of NUP subcomplexes, is incompletely understood. Further, it is not yet clear how exactly the scaffold interfaces with many of the peripheral structures. In particular, the nuclear basket—an eight-filament assembly protruding from the nuclear face of the NPC— is poorly understood. The basket has been implicated in diverse functions ranging from NPC assembly^23,24^ to mRNA export^25–27^, yet its molecular organization, its anchoring to the NR, and its interactions with transport complexes remain incompletely defined.

TPR is the major scaffolding component of the nuclear basket and essential for its assembly.^28–32^ It forms elongated coiled-coil homodimers that constitute the basket filaments. Other components, such as NUP153, NUP50, and the more recently identified ZC3HC1, contribute to basket assembly and transport regulation^29,30,33–38^, but structural information about their organization and interactions is limited. As a result, fundamental questions remain unanswered, namely: how are basket filaments anchored to the NPC scaffold, how do mRNA export factors such as the TREX-2 complex engage with the basket, and how do these interactions promote the passage of mRNPs into the central channel?

Here, we addressed these questions using in-situ crosslinking mass spectrometry (XL-MS), in-cell cryo-electron tomography (cryo-ET), AI-based structure modeling using AlphaFold 3 (AF3)^39^, and integrative modeling methods. This multimodal approach enabled us to build a substantially extended model of the human NPC including its nuclear basket. We identified five proteins previously not annotated as nucleoporins, defined the anchoring and structure of the basket filaments, and unexpectedly uncovered TREX-2 components as integral scaffolding components of the nuclear ring. Our model thus establishes a molecular framework to understand how mRNA export complexes engage with the NPC to promote transport through its central channel.

## Results

### An improved framework for integrative modeling of the human nuclear pore

Despite countless biochemical, structural and modeling analyses over decades, the structural model of the human NPC covers ∼70 MDa of the expected 120 MDa^18^ and the molecular basis for some of its functions, such as mRNA export, is not fully understood. We set out to fill in some of the remaining gaps, specifically focusing on multifunctional scaffold NUPs, including those at the nuclear basket. To obtain an overall spatial framework for modeling, we relied on previously published cryo-EM maps^18,40^ and *in situ* structural analysis of previously published large cryo-ET datasets of human macrophages^41^ and HEK293 cells (Xing et al. *unpublished*). We extracted 321 and 250 NPC particles from macrophages and HEK293 cells, respectively, and applied subtomogram averaging to their asymmetric units with a dedicated mask targeting the nuclear basket (Fig. S1a, see methods, ^41^). The nuclear basket filaments were readily apparent in the primary data and the averages (Fig. 1a,b). In both macrophages and HEK293 cells, they extended from the NR towards the nucleoplasm, almost parallel to the NPC’s central transport axis. The density of the basket filament appears to be divided into two thinner, individual filaments resulting in an overall two-pronged cross-section, that branches when it approaches the NR (Fig. 1a), pointing to two individual substructures that are further discussed below.

**Fig. 1.**
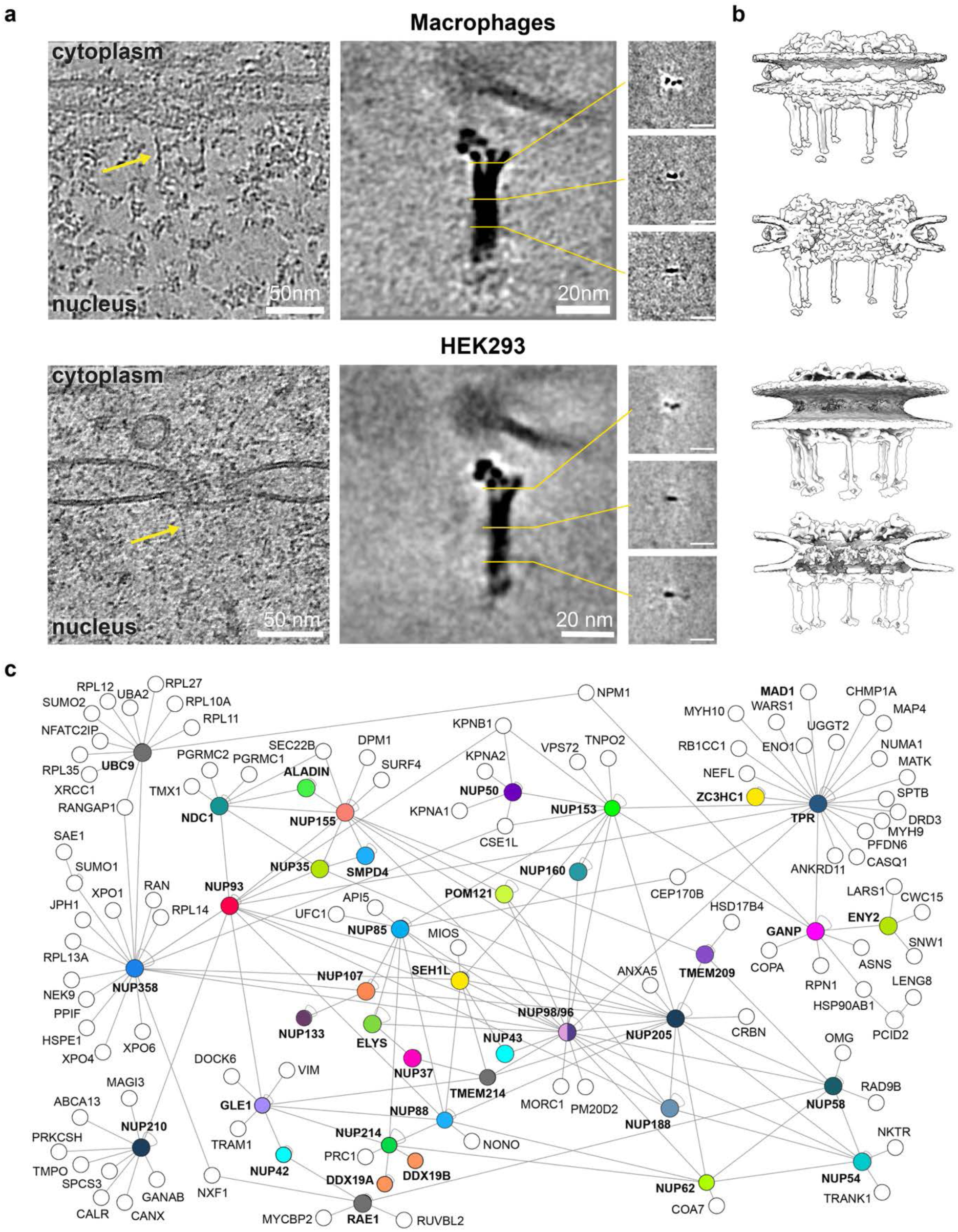
An improved framework for integrative modeling of the human nuclear pore. **(a)** Representative slices through tomograms of nuclear envelopes from macrophages and HEK293 cells (*left*) visualizing the nuclear basket filaments of the NPC (yellow arrows); central slices through subtomogram averages of the individual nuclear basket filaments (*right*). Insets show the three cross-sections as indicated. **(b)** Surface-rendered representations of composite cryo-ET maps of the entire NPC shown in side view (*top*) and sliced through the nucleocytoplasmic axis (*bottom*). **(c)** Crosslink map of human NUPs and associated proteins. Lines indicate at least one crosslink between the proteins. NUPs and stationary components of the NPC are shown color-coded (as throughout the manuscript).

To obtain complementary data on residue–residue proximities within the endogenous human NPC, we performed *in situ* XL-MS using an optimized approach (^42^, see methods). We identified 1600 unique residue-to-residue connections from three XL-MS datasets of HEK293 and Jurkat cells, within or across known NUPs, or those newly identified in this study (see below). The resulting proximity map considerably exceeds previous attempts made for the human NPC ^43^, and human nuclei ^44^, broadly covering NUPs, but also including multiple crosslinks involving the basket components TPR, ZC3HC1, NUP153, and NUP50, (Fig. 1c; Fig. S1b). Our dataset represents the deepest crosslink map of the human NPCs to date, suggesting multiple novel interactors and providing a large number of spatial restraints to guide and validate integrative modeling.

Although previous models of the human NPC architecture were thought to contain about 90% of the scaffold^18^, the respective tomographic maps still contained unassigned densities in distinct regions, suggesting the presence of additional, yet undiscovered components. Therefore, we searched our extensive XL-MS dataset for potential new interactions involving NUPs. To investigate whether any of these proteins could be part of the NPC, we assessed whether the crosslinks with known NUPs support high scoring AF3 models that, if placed into the map of the NPC, explain the currently unassigned density.

### SMPD4 and TMEM209 are inner ring scaffold nucleoporins

The IR contains several multifunctional scaffold NUPs, including NUP155, NUP205 and NUP93. It is subdivided into 32 core modules, organized into 16 inner and 16 outer copies arranged in C2 symmetry across the nuclear envelope. How exactly the inner and outer copies of these core modules are structurally specified remains unknown. Furthermore, the IR is embraced by the luminal ring (LR), composed of 64 copies of NUP210 (GP210). How the LR and IR are connected through the membrane such that they rotationally register, remains unknown.

Our XL-MS data identified two proteins proximate to well established IR NUPs. TMEM209, which is annotated as a nuclear envelope transmembrane protein, exhibited eleven crosslinks to NUP58, NUP205 and NUP155. SMPD4, a protein annotated as a sphingomyelin phosphodiesterase, underwent two crosslinks to known NUPs, one to NUP35 and one to NUP155 (Fig. 1c; Fig. S1b). A literature search revealed that both proteins were not only previously shown to localize to the NE^45–48^, but also had been independently found to interact with IR NUPs. Taken together, multiple lines of evidence strongly suggest that both SMPD4 and TMEM209 are *bona fide* IR NUPs.

We therefore set out to incorporate these two proteins into the model of the IR of the human NPC. AF3 identified high-confidence interactions with IR NUPs within three domains of TMEM209 that are separated by linker regions (Fig. 2a; Fig. S2a). Its NTD (aa 1–120) interacted with the NUP155 NTD and the transmembrane domain of NUP210, its middle region (aa 215–300) with NUP205 and its CTD (aa 325–561) with two NUP62 subcomplexes (Fig. 2a). The CTD of TMEM209 is therefore configured to link the inner and outer copies of the NUP62 subcomplex.

**Fig. 2.**
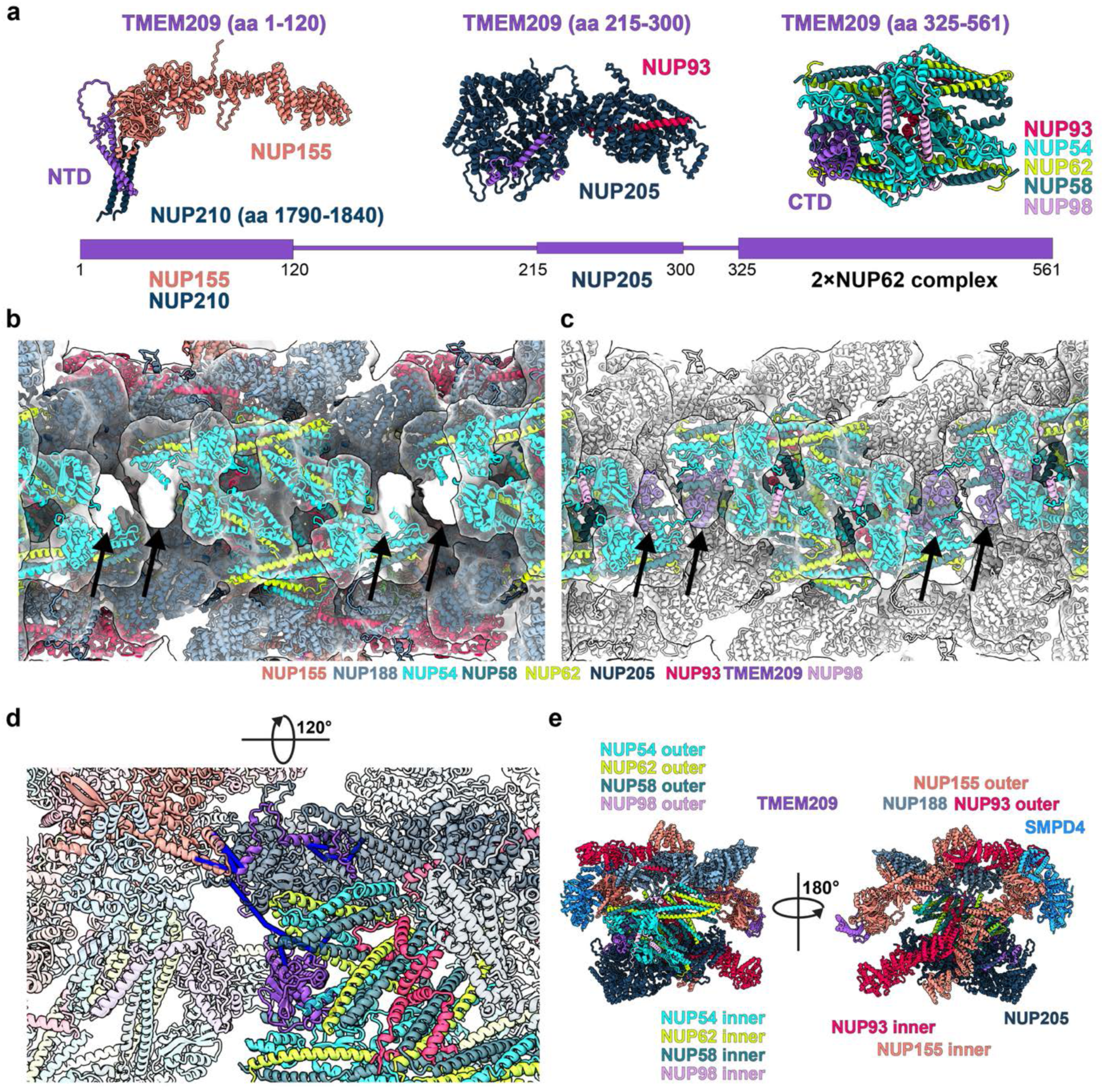
TMEM209 is a *bona fide* scaffold NUP of the IR. **(a)** AF3 models of the three subcomplexes involving TMEM209 (top; Fig. S2a) and their respective interaction sites within the domain architecture of TMEM209 (bottom). **(b)** Overlay of the previous structural model of the human NPC with the cryo-ET map (PDB-7R5K; EMD-14322, ^18^) as seen from the inside of the central channel; unassigned density indicated by black arrows; proteins color-coded according to Fig. 1c. **(c)** Same as **(b)** but additionally overlaid with the TMEM209 CTD**–**2**×**(NUP98**–**93**–**62**–**58**–**54) subcomplex positioned by systematic fitting (Fig. S3a) shown color-coded as in Fig. 1c; previously assigned proteins shown in white. **(d)** Crosslinks outgoing from TMEM209 supporting the assignment of the middle and C-terminal domain. Crosslinks are shown mapped onto the model as pseudo-bonds between Cα atoms of the crosslinked lysine residues (All detected crosslinks shown, satisfied restraints in blue). **(e)** Organization of the C2 symmetry plane of the IR. The same subcomplex is shown in context of the adjacent IR core modules, including NUP205, NUP188, NUP155, and SMPD4 (see Fig. 3).

Systematic fitting is a computational method for model placement into a cryo-EM map, which exhaustively interrogates potential configurations. For each fit, it scores the agreement of a given structural signature, here, an AF3 model, with the map. Subsequently, the statistical confidence is calculated from the overall score distribution for the top-scoring positions.^43^ Systematic fitting of the subcomplex consisting of the TMEM209 CTD–2×(NUPs 98,93,62,58,54) into the cryo-ET map of the human NPC (EMD-14322, ^18^) identified its position (Fig. S3a). This was consistent with the known position of the two respective NUP62 subcomplexes, whereby the TMEM209 CTD explained density that was previously unaccounted for (Fig. 2b, c). The XL-MS data further confirmed this, as we identified eleven crosslinks between TMEM209 and the surrounding NUPs (Fig. 2d). We therefore conclude that TMEM209 is an IR component with scaffold, linker and transmembrane NUP properties (Fig. 2e).

SMPD4 was crosslinked to NUP155 and NUP35 (Fig. 1c; Fig. S1b), the latter two were known to interact with each other.^18,49,50^ AF3 modeling identified a high-confidence interaction of SMPD4 with NUP155, but not with NUP35. In the model, the SMPD4 NTD engages with the NUP155 NTD, while the SMPD4 CTD contacts another region of the NUP155 NTD (Fig. 3a, Fig. S2a). Systematic fitting of the NUP155–SMPD4 model into the cryo-ET map of the human NPC (EMD-14322, ^18^) was consistent with the previously determined position of the inner copy of NUP155 (Fig. 3b; Fig. S3b). In addition, it explains the experimentally observed crosslink to NUP155 (Fig. S3c) and places the amphipathic and transmembrane α-helices of SMPD4 close to the membrane (Fig. 3c).

**Fig. 3.**
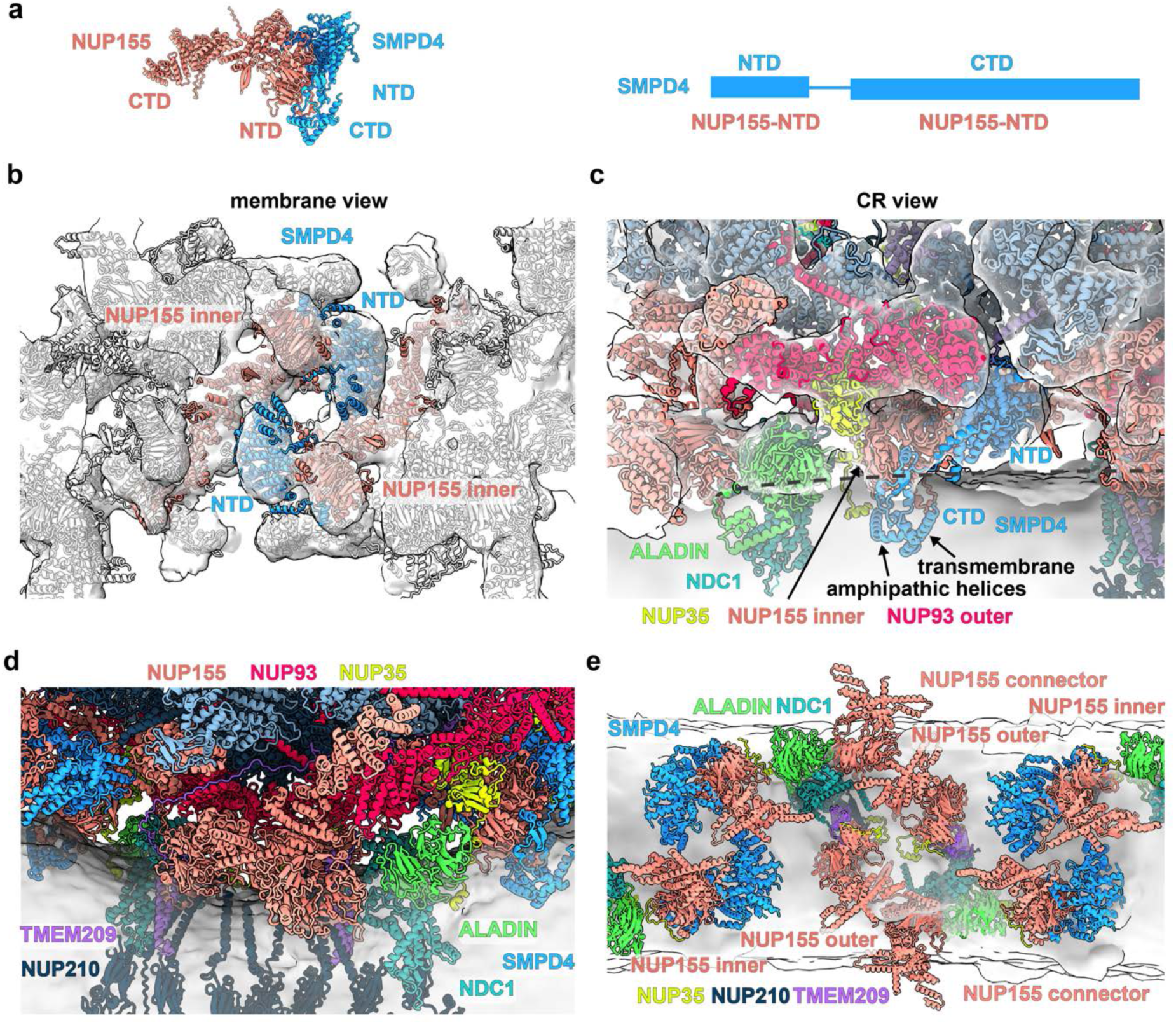
SMPD4 is a *bona fide* scaffold NUP of the IR. **(a)** AF3 model of NUP155**–**SMPD4 subcomplex (*left*; Fig. S2a) and domain architecture of SMPD4 (*right*). **(b)** Overlay of the previous structural model of the human NPC with the cryo-ET map (PDB-7R5K; EMD-14322, ^18^) as seen from the membrane (white) with the NUP155**–**SMPD4 subcomplex positioned by systematic fitting (Fig. S3b); proteins color-coded as in Fig. 1c. **(c)** Same as (**b**) but seen from the CR; proteins color-coded according to Fig. 1c. SMPD4 amphipathic and transmembrane α-helices are indicated. AF3 model of NUP93**–**NUP35 subcomplex positioned by systematic fitting (Fig. S3d). **(d)** Structural model of the IR and LR (this study) shown overlaid with the isosurface-rendered membrane (transparent grey) as seen from the CR; highlighting the interaction between IR and LR. **(e)** Overview of the NPC membrane-interacting NUPs of the IR as seen from the central channel. Membrane shown as an isosurface-rendered in transparent grey.

To further support our assignment of SMPD4 to the NPC, we performed immunofluorescence (IF) staining of SMPD4 in HEK293 cells, and found that it indeed localizes to the ER and nuclear envelope (Fig. S4a). We previously used mass spectrometry (MS) coupled to BioID labeling to identify NUPs proximate to ALADIN, another IR NUP that localizes to this region.^18^ Here, we reanalyzed the respective data and found both TMEM209 and SMPD4 prominently enriched (Fig. S4b). Therefore, the assignment of SMPD4 as an IR scaffold NUP is supported by several lines of evidence, namely XL-MS (Fig. 1c; Fig. S1b), proximity labeling MS (Fig. S4b), AF3 modeling (Fig. S2a), systematic fitting (Fig. S3b), the positioning of its transmembrane domain (Fig. 3c), IF (Fig. S4a) and previous literature.^45,46,51,52^

NUP35 is a dimeric IR nucleoporin positioned adjacent to the nuclear envelope membrane. It is known to interact with NUP155 and NUP93 via short linear motifs embedded in intrinsically disordered regions at both termini^53,54^, which previously led to the view that NUP35 is flexibly tethered to the IR scaffold.^18^ In contrast, AF3 identified a direct interface between NUP35 and NUP93 (Fig. S2a), reminiscent of the organization observed in the yeast NPC.^16^ This assignment is compatible with the known interaction network of NUP35. It is further supported by systematic fitting that identified both outer copies of NUP93 in the IR (Fig. 3c; Fig. S3d). The placement of NUP35 in this region is further supported by multiple crosslinks, however, no single conformation or copy of the NUP35 dimer can simultaneously satisfy all observed restraints (Fig. S3c), suggesting the presence of alternative conformations, or alternatively, additional copies of NUP35. Taken together, these observations indicate that the precise positioning and conformation of NUP35 and SMPD4 are likely dynamic and warrant further investigation in future studies (see limitations).

Overall, we conclude that both TMEM209 and SMPD4 are transmembrane scaffold NUPs of the IR, whereby TMEM209 also has linker NUP properties. Interestingly, AF3 identified an interaction of the TMEM209 NTD with the transmembrane domain of NUP210 (Fig. 2a; Fig. S2a) that in turn interacts with NUP93 (Fig. S2a; Fig. S3e), consistent with the XL-MS data (Fig. S3e). These data suggest that TMEM209 links the LR and IR (Fig. 3d), in line with a recent study that functionally characterized TMEM209.^55^ Our analysis further clarifies that both TMEM209 and SMPD4 contribute to specifying the inner and outer core modules within the IR. Thereby, three dedicated copies of the multifunctional scaffold NUP155 form three mutually exclusive interfaces with specific binding partners (Fig. 3e), namely ALADIN/NDC1^18^, TMEM209/NUP210 and SMPD4 (Fig. S3f). Furthermore, the CTD of TMEM209 is configured to link the inner and outer copies of the NUP62 subcomplex (Fig. 2e).

### The TREX-2 component GANP binds the basket attachment region of the nuclear ring

Compartment specific peripheral structures attach to the CR and NR of the human NPC. The cytoplasmic filament complexes that emanate from the CR into the immediate nuclear pore proximity in the cytosol are understood in considerable detail.^19,56^ However, how their nuclear counterparts are attached to the scaffold of the NR remains less well understood. Therefore, we next investigated unassigned densities in the NR. We fitted the previous model of the human NPC scaffold^18^ into the cryo-ET map of the human NPC obtained from macrophages^41^, and observed unassigned densities in between the inner and the outer copies of NUP85 bridging towards the outer copies of NUP96, NUP107 and NUP133 (Fig. 4a). Inspection of higher-resolved EM maps of the *Xenopus laevis* NPC (EMD-31065, EMD-31941, EMD-32394; ^40^) and the *Saccharomyces cerevisiae* (yeast) NPC (EMD-24231; ^16^) revealed similar unassigned densities (Fig. S5a-d) with visible secondary-structure elements in the corresponding region (Fig. S5c). This region and the respective unassigned density overlap with the proximal part of the nuclear basket filaments when subjected to focused refinement (Fig. 1b), suggesting that the density represents their attachment sites.

**Fig. 4.**
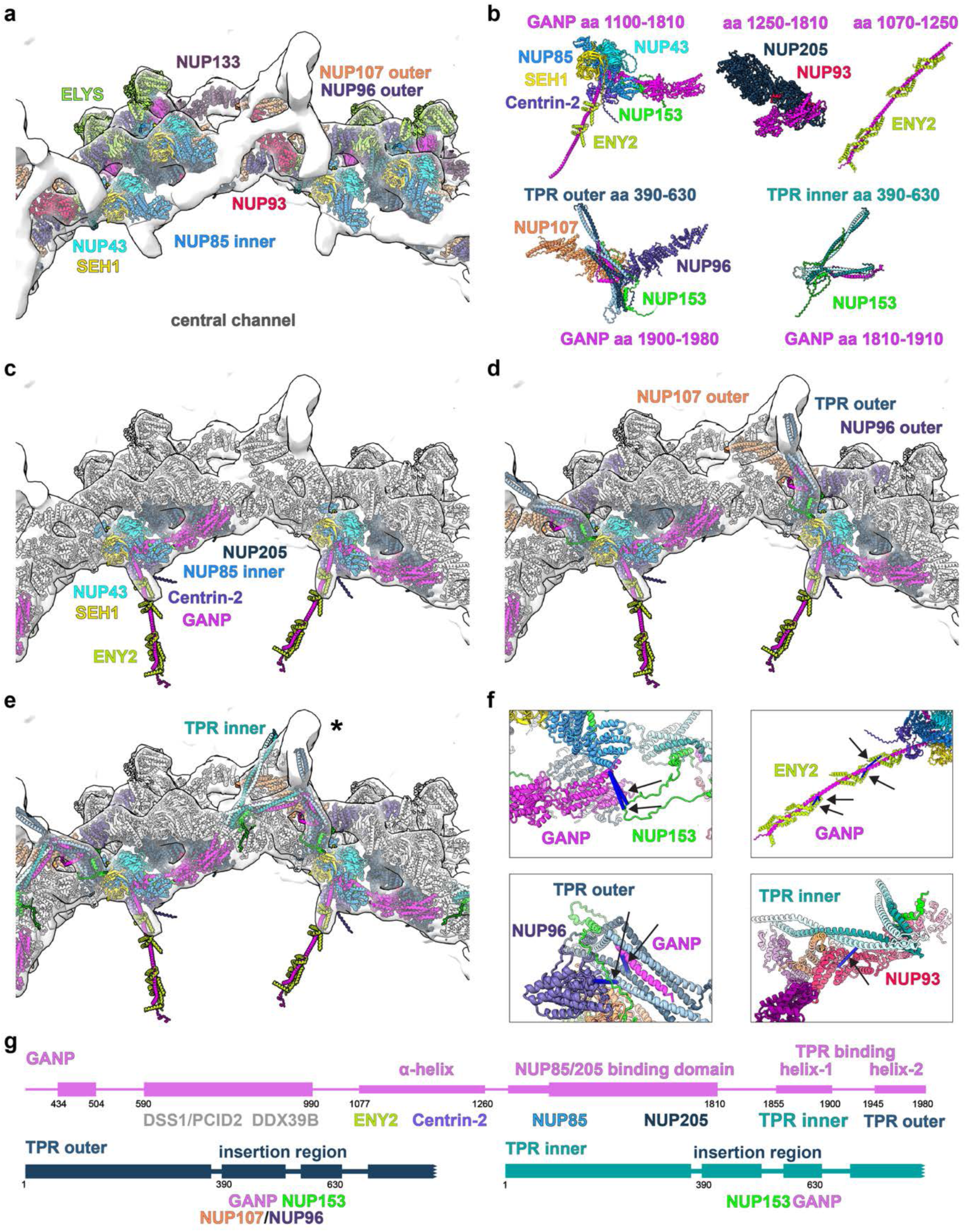
The attachment of the nuclear basket filaments to the scaffold of the NR involves GANP. **(a)** Previous structural model of the human (PDB-7R5K; ^18^) fitted into the NR region of the cryo-ET map of the human macrophage NPC as seen from the nucleoplasm; unassigned density in white; proteins color-coded according to Fig. 1c. **(b)** AF3 models of the subcomplexes including GANP and the TPR insertion region (bottom). See also Fig. S2. (**c)** Same as (a) but overlaid with the scaffold-interacting part of the TREX-2 subcomplex positioned by systematic fitting (Fig. S6) shown color-coded as in Fig. 1c; previously assigned proteins shown in white. **(d)** Same as (c) but showing the subcomplexes involving the outer TPR copy fitted according to Fig. S7 in addition. **(e)** Same as (d) but showing the subcomplexes involving the inner TPR copy fitted according to Fig. S7. The unassigned density marked by * corresponds to the basket filament, as shown in Fig. 5. **(f)** Crosslinks validating the assignments shown in (c)-(e), depicted as pseudo-bond between Cα atoms of the crosslinked lysine residues (All detected crosslinks shown, satisfied restraints in blue). **(g)** Domain architectures of GANP and TPR, the latter shown separately for outer and inner copies and truncated after the insertion region.

We found that the TREX-2 component GANP was crosslinked to TPR, NUP153 and ENY2 (Fig. 1c; Fig. S1b). Thus, we probed interactions of NUPs with TREX-2 components using AF3 and obtained the following high-confidence models: NUP85–SEH1–NUP43–GANP–NUP153, NUP85–SEH1–NUP43–GANP–NUP153–Centrin-2–ENY2×2, GANP–ENY2×4, NUP205–NUP93–GANP (Fig. S2b), and, with the requirement that TPR is an obligate dimer^30^, NUP96–NUP107–TPR×2–NUP153–GANP (Fig. S2c), and TPR×2–NUP153–GANP (Fig. 4b; Fig. S2d). Systematic fitting assigned the NUP85–SEH1–NUP43–GANP–NUP153 subcomplex to a position which overlapped with the known location of the inner copy of NUP85 subcomplex (Fig. 4b,c; Fig. S6a). It further positioned an independent AF3 model of GANP in complex with NUP205–NUP93 such that it overlapped with the known position of NUP205 in the NR (Fig. S6a). Thus, systematic fitting of these two models independently yielded the same assignment of the GANP NUP85/205 binding domains (Fig. 4b,c; Fig. S6a). This assignment is also in agreement with the XL-MS data, which contained multiple crosslinks supporting this configuration (Fig. 4f). We conclude that the GANP region comprising residues 1260–1810 contains binding domains for NUP85 and NUP205, and is positioned into previously unassigned density of the NR proximate to the Y-complex. We furthermore modeled the adjacent TREX-2 components Centrin-2 and ENY2 in complex with the NUP85 subcomplex and NUP153, and found that they bind to the GANP region (aa 1077–1260), likely to facilitate its extension away from the scaffold and towards the central channel (Fig. 4b, c).

To explore the conservation of the GANP interaction with the Y complex, we first modeled the similarly structured *X. laevis* subcomplex (Fig. S2j) and systematically fitted it into the respective high-resolved EM map (Fig. S6b), resulting in the same assignment. In this case, the respective α-helical fold of the NUP85/205 binding domain of GANP is even explained well by the experimentally observed secondary structure elements, which are distinguishable due to the high resolution of the EM map (Fig. S6b). Second, we assessed this interaction in yeast and noted that the yeast homolog Sac3 has a distinct domain architecture and lacks the yNUP192 (hNUP205)-binding domain, consistent with the absence of yNUP192 in the yeast NR. Accordingly, no density corresponding to the human GANP NUP85/205-binding domain was observed in the yeast NR. Nevertheless, AF3 modeling of the yeast Sac3 with Nup85 subcomplex (Fig. S2k) suggested that Sac3 binds both copies of Nup85 in the NR, positioning the yeast TREX-2 components in a manner analogous to the corresponding region of the human NR, consistent with the distinctive experimental density observed in yeast (Fig. S6c). Together, these observations further support the assignment of the corresponding density in the human NPC to the human GANP NUP85/205-binding domains.

TPR is the major component of the nuclear basket filaments. Biochemical studies revealed that TPR consists of a ∼1500 aa long structured N-terminal domain (NTD), which is composed of α-helices that are arranged in a double-stranded coiled-coil^30^, and a largely unstructured flexible C-terminal domain (CTD). The part of the NTD is known to interact with the scaffold of the NR^30^ (aa 390–660) is from here on referred to as the insertion region. To determine if GANP may contribute to the attachment of the nuclear basket filaments to the NR, we systematically fitted NUP96–NUP107–TPR×2–NUP153–GANP subcomplex into the cryo-ET map of the NR (Fig. S7a). This analysis positions the TPR×2–NUP153–GANP subcomplex into the previously unassigned density within the nuclear basket attachment region (Fig. 4d; Fig. S7a) and correctly overlaps NUP96 and NUP107 with the outer Y-complex. We will further refer to this TPR as the outer copy.

For further validation, we generated a homologous *X. laevis* model (Nup96–Nup107–Tpr×2–Nup153-Ganp) and systematically fitted it into the cryo-EM map of the NR of the *X. laevis* NPC (EMD-31065), confirming this assignment (Fig. S7b). The coiled-coils formed by the TPR insertion region and the C-terminal α-helix of Ganp were consistent with α-helical secondary structures (Fig. S7b). This interaction is furthermore supported by the XL-MS data (Fig. 4f), and confirms a motif in TPR that was previously identified to mediate its binding to the NPC using mutagenesis studies.^30^ We therefore conclude that the C-terminal region of GANP contributes to the NR attachment of the TPR insertion region (Fig. 4g).

Overall, our findings show that TREX-2 is a NUP subcomplex whereby GANP contributes to both, scaffolding of the NR and attachment of the nuclear basket filaments. Thus, the TREX-2 complex is ideally positioned to associate the additional components, PCID2, DSS1 and the DDX39B helicase, which are important for mRNP processing^57^, with the entry of the central channel of the NPC.

### Two TPR insertion regions are attached to distinct interfaces at the nuclear ring

The analyses described above did not yet explain the entire unassigned density observed in the nuclear basket attachment region (Fig. 4a,d). Since the basket filaments appear to branch where they approach the NR (Fig. 1a) and because of the previously determined TPR stoichiometry^58^, we hypothesized that two TPR dimers might be attached to individual sites at the NR. We therefore set out to identify a second attachment site for the TPR insertion region.

Indeed, an AF3 model of the TPR insertion region in complex with NUP153 and GANP matched the remaining unassigned density in the basket attachment region (Fig. 4b,e; Fig. S7a right panel), supporting the presence of a second TPR binding site (inner copy of TPR) within the NR. Furthermore, the XL-MS data independently support the assignment of each of the two TPR copies (Fig. 4f). In this model, the two C-terminal α-helices of GANP bind separately to the inner and outer TPR dimers, such that a single copy of GANP engages two TPR dimers simultaneously (Fig. 4b-e; Fig. S2c,d; Fig. S7). We therefore hereafter refer to these helices within GANP as TPR-interacting helix-1 (aa 1855-1900) and helix-2 (aa 1945-1980), which bind to the inner and outer TPR dimer, respectively.

Again, we next asked if this interaction was conserved in the yeast NPC. We generated several AF3 models containing the insertion region of the two yeast homologs of TPR, Mlp1 and Mlp2, together with other NUPs (Fig. S2l). We systematically fitted the resulting models into the cryo-EM map of the NR of the yeast NPC (EMD-24231 ^16^) (Fig. S7c), indeed revealing two Mlp1/2 attachment sites: Mlp2 binds the outer Nup84–Nup145C subcomplex (Fig. S7c), while Mlp1 bridges the outer and inner Nup84 copies (Fig. S7c). The arrangement of two homodimers per filament explains the observed electron density better than the previously proposed single homodimer.^58^ We therefore conclude that TPR is a multifunctional scaffold NUP that associates with the NR at two different sites.

### A dimer of TPR dimers constitutes the core of the nuclear basket filaments

We next aimed to structurally model the basket filaments. Rod-shaped homodimers of TPR^30^ constitute the basket filaments^28,29^, whereby one part of the TPR homodimer folds back onto another part of itself, such that N- and C-termini extend into the nucleoplasm. We first focused on a single TPR homodimer. Modeling attempts of the entire subcomplex using AF3 did not yield high-scoring models. However, TPR is extensively covered by the XL-MS data and numerous intra-molecular crosslinks between domains distant in sequence were found (Fig. 5a), indicating that these regions must be spatially proximal in the tertiary structure. We therefore used AF3x, a protocol that considers crosslinks as spatial restraints during AF3 modeling^59^, to assemble the dimer spanning residues 1–1460 (see methods). We selected five structural arrangements that best satisfied the XL-MS data and had the highest pLDDT scores (Fig. S8a). Although the models differed slightly in the relative arrangement of the TPR coiled-coil fragments (aa 1-390 and 660–1460) (Fig. S8a), our results converged on a consistent architectural principle: All models are based on an antiparallel bundle of four stacked TPR coiled-coils spanning residues 1–390 and 660–1460. Those established extensive lateral interactions that are likely responsible for the rigidity and stability of the basket filaments. The insertion region (∼aa 390–660) remained outside of the coiled-coil stack and did not form specific interactions with the coiled-coil bundle (Fig. S8a).

**Fig. 5.**
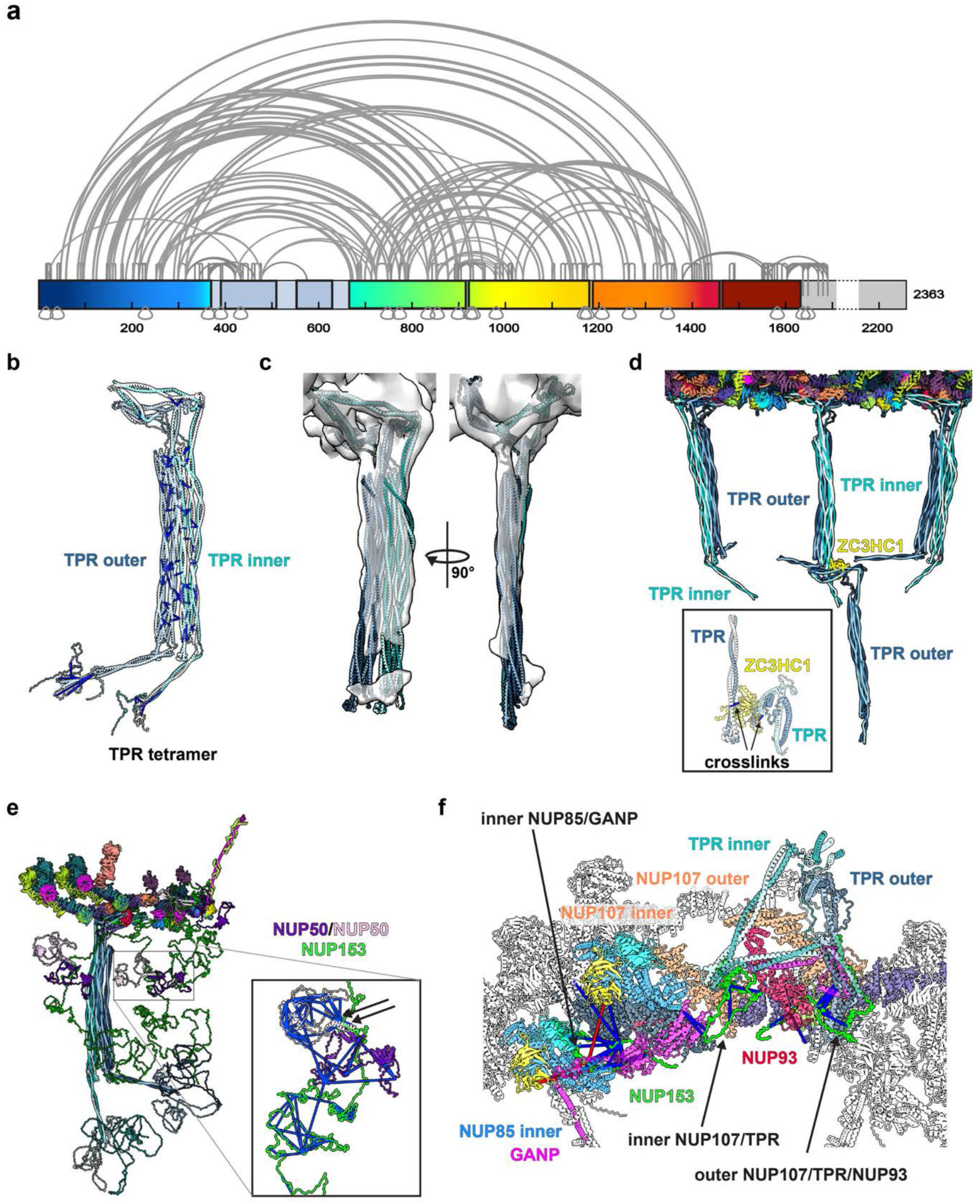
Structural organization of the nuclear basket. **(a)** Intra-protein crosslinks mapped onto the primary structure of TPR; Rectangles represent individual coiled-coil fragments derived from structural models of the TPR dimer (rainbow from N to C). **(b)** Crosslinks mapped onto the structural model of TPR tetramer (aa 1-1670, Fig. S9a). Only crosslinks that are satisfied (≤40Å) and between residues separated by at least 20 aa are shown for clarity (blue); all crosslinks are shown in Fig. S9b and c. **(c)** Structural model of the TPR tetramer fitted into the cryo-ET map of the human macrophage NPC. **(d)** Structural model of three adjacent rotational segments of the NR shown together with the respective TPR tetramers and ZC3HC1 assembled by superposition of the respective AF3 models shown in Fig. S2. Inset shows crosslinks of ZC3HC1 with either end of the TPR filament. **(e)** Structural model of one NR and nuclear basket subunit assembled by superposition of the models shown in Fig. 4e and 5b, and the AF3 models of NUP153 and NUP50 shown in Fig. S2e. Intrinsically disordered domains and C-terminal coiled-coils of TPR are placed in arbitrary positions. Inset shows all crosslinks for NUP153 (aa 400**–**1200) and NUP50 with arrows indicating two crosslinks validating the interaction between NUP50 (aa 150**–**200) and NUP153. **(f)** Crosslinks validating the proximity of NUP153 (aa 240**–**400) to the NR, shown in the context of the structural model of a subunit of the NR as seen from the nucleoplasm. Satisfied crosslinks in blue and those **>**40 Å in red (the latter all map into disordered regions and can be satisfied due by flexibility). Crosslinks are depicted as pseudo-bonds between Cα atoms of lysine residues.

We next assembled the TPR tetramer using the above-discussed TPR homodimers as templates in ColabFold.^60^ To reduce computational complexity, we focused on residues 1–320 and 660–1460, excluding the insertion region (Fig. S8b). The five top scoring structural arrangements had comparable scores and satisfied the XL-MS data to a similar extent (Fig. S8b). We integrated the full-length TPR tetramer into a model of an NPC asymmetric unit by aligning the filament structure and insertion regions with the cryo-ET map from macrophages^41^, while positioning the C-terminal coiled-coils and disordered regions of TPR arbitrarily (Fig. S9a). This model agrees with domain mapping experiments of TPR^28^, satisfies additional crosslinks (Fig. 5b), explains the observed electron optical density comprehensively (Fig. 5c), aligns with the two-pronged cross-section of the subtomogram averages (Fig. 1a; Fig. 5c), and, is consistent with the identification of two attachment sites for the TPR insertion region at the NR (Fig. 4d,e).

### How ZC3HC1 extends the nuclear filaments

ZC3HC1 was previously identified as a *bona fide* nuclear basket component.^36,37^ ZC3HC1’s nuclear basket interaction domain (aa 72–290 and 398–467) binds to TPR and is necessary and sufficient for association with the basket.^38^ Inspired by a recent study suggesting a role for the yeast ZC3HC1 homolog Pml39 in extending basket filaments^61^, along with our observation of extended filaments in tomograms (Fig. S10a), we explored the role of ZC3HC1 in basket filament organization. AF3 modeling suggested that ZC3HC1 interacts with both the N-terminal region (aa 40-60) of TPR as well as the TPR insertion region (Fig. 5d; Fig. S2f), which are positioned on opposite ends of the TPR filament. Nevertheless, both interactions are supported by the XL-MS data (Fig. 5d, inset). Therefore, in agreement with prior studies on ZC3HC1’s interaction with the nuclear basket^36–38^ and the recently proposed model for the yeast nuclear basket^61^, our data suggest that ZC3HC1 elongates basket filaments end-to-end, by linking consecutive TPR homodimers end to end (Fig. 5d; Fig. S10a,b).

### NUP153 tethers NUP50 to the nuclear basket

NUP153 contributes to anchoring TPR to the NPC ^29,30^ at the Y-complex ^33^, and interacts with NUP50, that plays a role in nuclear export mediated by the exportin CRM1^34^ and Importin-α mediated import^35^. To incorporate these two components into our model, we generated the following high-scoring AF3 models of subcomplexes involving basket NUPs: NUP96–NUP107–TPR×2–NUP153–GANP, TPR×2–NUP153–GANP, NUP85–SEH1–NUP43–GANP–NUP153, NUP107–NUP153, NUP93–NUP153 and NUP50×2–NUP153 (Fig. S2c-e). This analysis suggests that NUP153 contains multiple interaction regions: residues 280–320 interact with NUP85, 250–275 with NUP107, 305–320 with TPR, 360–370 with NUP93, and 520–560 with NUP50. At least three different copies at five binding sites for NUP153 at the inner copy of NUP85, as well as inner and outer copies of NUP107/TPR – all bound to GANP – are supported by density in the cryo-EM map of *X. laevis* (Fig. S10c; Table S6) and the XL-MS data (Fig. 5e,f).

NUP50 attaches to the flexible, disordered region of NUP153 via its dimerization domains (aa 150–200) (Fig. 5e). The interaction involves NUP153 residues 520–560, which is consistent with previous findings showing that the NUP153 aa 401–609 region is necessary and sufficient for its interaction with NUP50 ^62^. The disordered regions of NUP153 were modeled in arbitrary conformations (Fig. 5e) (see limitations of the study). The resulting partial model of NUP153–NUP50 was then added for each NUP153 copy. In the resulting model of the nuclear basket, NUP153 tethers NUP50 in a flexible position (Fig. 5e), consistent with the XL-MS data (Fig. 5e inset). Hereby NUP153 and NUP50 are positioned at the nuclear entrance of the central channel within the exclusion zone formed by the basket filaments.

Finally, we generated a composite model of the entire NPC by combining information of the human NPC scaffold from Mosalaganti et al.^18^ and this study, the cytoplasmic filaments^56^ and the nuclear basket (this study) (Fig. 6; Fig. S10a,b). We used Assembline^63^ to position and refine all scaffold components in the cryo-ET map from macrophages^41^. Disordered regions were placed in arbitrary positions. Previous studies estimated the molecular weight of the human nuclear pore to about 120 MDa.^18^ Taking the compositional refinement of this study into account, the sum of the molecular weight of all full-length proteins contained in the human NPC is about 126 MDa (Table S5), >102 MDa of which are contained in our model (Fig. 6a). Here, the nuclear basket accounts for 18.8 MDa, excluding the GANP interactors DSS1, PCID, DDX39B that would contribute another 0.8 MDa (Fig. 6; Fig. S2g). This estimate also does not include nuclear transport receptors that are prominently bound to the central channel, lipids or post-translational modifications of NUPs.

**Fig. 6.**
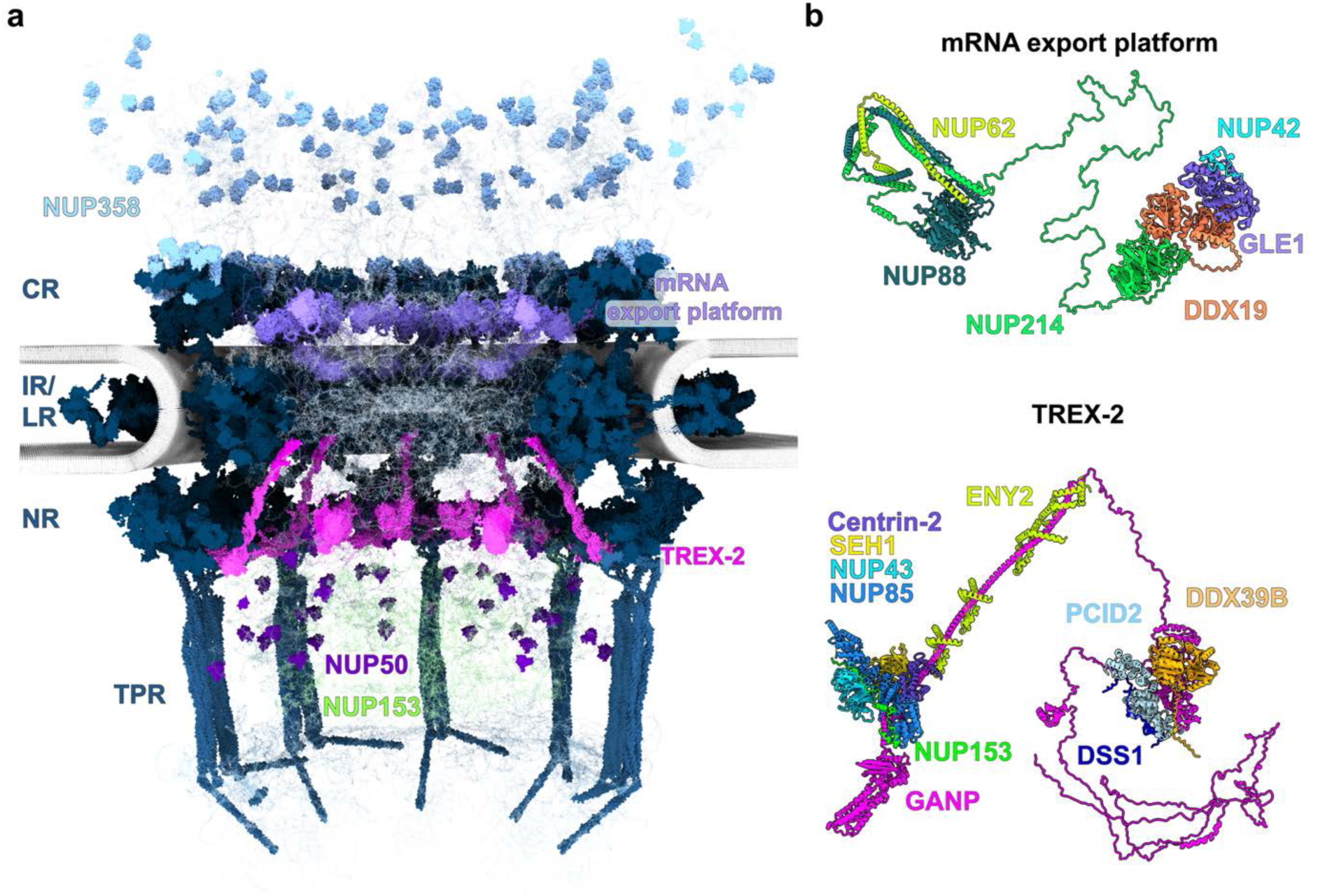
How mRNP-interacting helicases are associated with the NPC scaffold. **(a)** Structural Model of the entire human NPC including the nuclear basket and TREX-2 (Fig. S2g), as well as the cytosolic mRNA export platform (Fig. S2h). The disordered regions and folded domains connected to those were positioned arbitrarily. **(b)** AF3 models of the NUP214-subcomplex (mRNA export platform)^64^ and the TREX-2 complex^65^ (Fig. S2g, h) highlighting the flexible association with the DDX19 and DDX39B helicases, respectively.

Our model illustrates how the organization of the high copy number NUPs, NUP358 and NUP153, results in a highly crowded region at each of the two entrances to the central channel, which may be important for the initial docking of import cargos, and to locally concentrate RanGAP. It furthermore places the two modules important for mRNP export, the NUP214 and TREX-2 subcomplex, at either entrance of the central channel (Fig. 6; Fig. S2g,h). Our model thereby provides a molecular basis for more accurate mechanistic studies and computer simulations of nuclear transport, in particular mRNA export.

## Discussion

Our integrative analysis that combines advanced XL-MS with AF3-based modeling and systematic fitting into cryo-EM density identified five previously unrecognized *bona fide* nucleoporins: SMPD4, TMEM209, and the TREX-2 components GANP, Centrin-2, and ENY2. Previous studies largely assumed that TREX-2 binds peripherally to the nuclear basket.^8,10^ This notion likely arose because TREX-2 components were not detected by proteomics-based analyses of the NPC^8,11,66,67^, with the notable exception of Cdc31, the yeast homolog of Centrin-2.^68^ Our data however show that the TREX-2 component GANP contains an NPC binding domain that is an integral part of the NR, and not distally associated with the basket. In yeast, NPCs with and without nuclear baskets exist^58^, and TREX-2 association was shown to preferentially occur at NPCs with a nuclear basket.^58,69,70^ At this stage, we cannot definitively answer the question if TREX-2 is always bound to all human NPCs, but our data indicates that a basket and TREX-2-less NPC would likely constitute a minor subset.

This placement has direct implications for mRNA export. TREX-2, located at the nucleoplasmic entrance of the central channel, provides a structural platform for the remodeling of export-competent mRNPs that may occur as follows: The PCID2-interacting domain of GANP^71^ is flexibly connected to the NR via a long linker (∼70 aa). Therefore, the associated complex of the DEAD-box ATPase DDX39B (UAP56), DSS1 and PCID2 can explore a relatively large space within the nuclear basket region and clamp onto the mRNP that is to be exported.^8^ The multiple copies of the TREX-2 complex may do this simultaneously to tether the mRNP to the entrance of the central channel. Recent reports proposed that a trigger loop in GANP promotes unclamping of DDX39B and its release from the mRNP^65,71,72^, which could result gradual melting of the mRNP into the FG-meshwork. Many previous studies assumed that the respective steps occur in the nucleoplasm. Our model however places GANP- and DDX39B-mediated remodeling of the mRNP directly at the nuclear entry of the central channel. Once inside the central channel, the FG-repeat binding protein NXF1 mediates interactions between the mRNP and FG-NUPs.^67^ Upon exit from the central channel, NXF1 binds to the FG-repeats of the cytoplasmic nucleoporin NUP214. This leads to further mRNP remodeling involving the DEAD-box ATPase DDX19, likely removing NXF1^73,74^, and consequently release of the mRNP into the cytosol. As proposed, this mechanism would prevent re-binding of the mRNP to the NPC and ensure unidirectionality of transport. It also argues that mRNA export is coordinated by at least two spatially distinct platforms—TREX-2 and NUP214—upon entry into and exit from the central channel.

The structural organization of the TREX-2 subcomplex is intriguing. Similar to the NUP214 subcomplex, the helicase-containing module consisting of DDX39B–PCID2–DDS1 is flexibly connected to the scaffold and thus can explore a large conformational space within the central channel (Fig. 6). In case of the TREX-2 subcomplex, however, at least four copies of ENY2 appear to stabilize the helix of GANP (aa 1077–1260) that links the NR scaffold to the helicase containing module (Fig. 4; Fig. 6b). It is interesting to speculate that this intriguing motif may have a regulatory function and could be used to position the mRNP-interacting helicase towards the central channel on demand. Such a concept is not unprecedented, and has previously been observed for the cytosolic mRNA export platform in yeast, where the DID/Dyn2 motif extends an otherwise disordered stretch in Nup159, although its regulatory function remains underexplored.^75^ We note that the directionality of this helix of GANP with respect to the NR scaffold in our model is set by the orientation of the corresponding AF3 model, but it may well be bent into another direction, possibly by additional factors that are not modeled here.

Transmembrane proteins are notoriously difficult to identify by mass spectrometry.^76^ The present notion is that human NPCs contains three transmembrane NUPs, NUP210, POM121 and NDC1, although only NDC1 appears to be essential for membrane anchoring of the NPC.^77,78^ Therefore, amphipathic motifs in soluble NUPs were thought to critically contribute to NPC membrane anchoring.^79^ Here, we identify and assign structural roles to two additional transmembrane NUPs - SMPD4 and TMEM209. Both were previously reported to interact with nucleoporins^46,47,80^, but to the best of our knowledge not yet discussed as structural NPC components. SMPD4, although annotated as sphingomyelin phosphodiesterase 4, exhibits a mostly α-helical fold that markedly differs from the conserved fold of such enzymes.^81,82^ No enzymatic activity of purified SMPD4 has been reported, but inferred based on activity detected in cell extracts.^83^ We propose that both SMPD4 and TMEM209 are scaffold NUPs in the IR that contribute to anchoring the NPC IR in the membrane. TMEM209 also acts as a linker NUP bridging the inner and outer copies of the NUP62 subcomplexes and contacting NUP210. Together with ALADIN–NDC1, the mutually exclusive interaction partners TMEM209 and SMPD4 specify the different configurations of the multifunctional scaffold NUP155 (Fig. 3e). The interaction network of TMEM209 explains how the IR and the LR associate with each other through the fused membranes and rotationally register. POM121 was crosslinked to NUP188 and NUP155 (Fig. 1c) and these interactions are supported by AF3 models with excellent and moderate scores, respectively (Fig. S2a). One possible interpretation is that POM121 is anchored to the IR where it interacts with NUP188 and projects along the connector copy of NUP155 towards the Y-complex, consistent with a previously proposed role of POM121 in linking the IR and Y-complexes.^84^ Overall, our findings emphasize that the structural importance of transmembrane NUPs for NPC membrane anchoring may have been underestimated. Interestingly, the XL-MS data suggest yet another candidate transmembrane NUP, namely TMEM214 (Fig. 1c), that remains to be further validated in the future.

We present a molecular model of the nuclear basket and its attachment to the scaffold (Fig. 6). A striking difference between the nuclear and the cytoplasmic filaments is the incorporation of an elongated rigid element into the basket filaments. A need for this is possibly explained by the polymer-like properties and high local concentration of chromatin at the nuclear side of the NPC. The eight rigid filaments could constitute a physical barrier for heterochromatin that is laterally attached to the inner nuclear membrane, thus maintaining an exclusion zone underneath the basket.^31,85^ This protected space likely contains the disordered domains of NUP153, NUP50 and TPR and can be further extended by ZC3HC1. This model could explain how large cargos such as the HIV capsid, pre-ribosomes or mRNPs can pass the nuclear basket region without tangling.^86^

Previously, a distal ring was proposed as a structural constituent of the nuclear basket^87,88^, which we do not directly observe in our data. One could speculate that the two C-terminal coiled-coil domains (aa 1460–1628) emanating from proximate TPR protomers could fold onto each other in an antiparallel fashion and form a distal ring. However, neither the cryo-ET data nor AF3 modeling support this idea. We cannot formally exclude that such or alternative arrangements may form, for example in the absence of heterochromatin in germline cells, as a nuclear basket was initially reported in *X. laevis* oocytes using heavy metal shadowing EM.^87,88^ Such arrangements may furthermore involve yet unidentified components.

Taken together, our work underscores the power of combining XL-MS, cryo-ET, AI-based structural and integrative modeling, as it allowed us to uncover novel components and organizational principles within the human nuclear pore, a complex that has been extensively characterized. Our updated framework structurally defines the NPC as a multifunctional assembly that not only mediates bidirectional protein transport and coordinates mRNA remodeling, but also establishes a chromatin exclusion zone and a reaction space for cargo complexes beneath the central channel.

### Limitations of the study

Although we can confidently conclude that TMEM209, SMPD4 and the TREX-2 complex are NPC components based on multiple lines of evidence we secured, our data may less strongly support some of the individual domain interactions. This is exemplified by the interface between NUP210 and TMEM209/NUP155 identified by AF3, but not secured by XL-MS data. It is however consistent with the local interaction networks and the electron optical density. An overview of how individual experimental data and AF3 models support the respective assignments is provided in Table S6. Furthermore, our models should be interpreted primarily at the architectural, rather than at the atomic level.

While AF3 models for various NUP subcomplexes exhibit very high confidence scores, the absolute values were lower for TPR homodimers. This likely reflects the fact that elongated coiled-coils are underrepresented in the AF3 training data. Thus, the scores for TPR models are not reliable indicators of correctness and these were not judged based on this criterium. Importantly however, the TPR models we report are exhaustively covered and supported by XL-MS data, cryo-ET densities and various prior biochemical experiments.^28–30^ Although alternative models are possible for the exact geometry of the coiled-coil domains of TPR, they all adhere to the same basic architectural principles (Fig. S8a-c).

The structural model of the NPC we put forward is a single state model that has limitations in conceptualizing the properties of dynamic NUPs, as exemplified in the following. Our false discovery rate of our XL-MS data is calibrated by stringent statistical methods (see methods; ^89^). It is satisfied to 87% by the previous scaffold model of the human NPC (PDB-7R5K^18^), to 92% by the single state model from this study and to 94% if alternative conformations are considered. The issue of capturing the dynamic nature is exemplified by TPR that displays crosslinks that cannot occur simultaneously, e.g. end-to-end arrangements of TPR homodimers that extend the basket filaments, as well as folding of the insertion region or the C-terminal coiled-coil segment onto the basket filament (Fig. S9b). Our XL-MS data further suggests that MAD1 can align with one TPR dimer, but does not reveal if this interaction occurs at the NPC (Fig. S9e). Another example of this limitation are the disordered regions or flexibly connected parts of NUP153, NUP50, NUP358, POM121, and NUP98, as well as the NUP214, NUP62 and TREX-2 subcomplexes, and the C-terminal disordered domain of TPR, which are represented at arbitrary positions in our model. Furthermore, GANP helix (aa 1077–1260) that points towards the central channel is oriented as within the respective AF3 model, but its exact orientation and extension is likely dependent on ENY2, and possibly other factors, and thus is likely dynamic. At last, multiple lines of evidence indicate that NUP153 occurs in multiple copies per spoke, but not all of those may be always occupied. Appropriate conceptualization of such dynamics would require a multistate ensemble model, which will be the focus of future efforts.

## RESOURCE AVAILABILITY

The original tilt series will be deposited in the Electron Microscopy Public Image Archive (EMPIAR). EM maps will be deposited in the Electron Microscopy Data Bank (EMDB). The structural models will be deposited in PDB-Dev. The XL-MS-data will be made available upon publication of the manuscript.

## Acknowledgements

We would like to thank Jan Kosinski from EMBL Hamburg for advice on modeling and critical reading of the manuscript. We thank the Max Planck Computing and Data Facility for support with scientific computing. We thank the Central Electron Microscopy Facility at Max Planck Institute of Biophysics for support with data acquisition. We thank EMBL Heidelberg for cluster computing. We thank Prof. Si-min He and Pengzhi Mao for providing a demo version of pLink3 (v3.0.16). This work was funded by the following sources: Max Planck Society and in part by the German Research Foundation (CRC 1507 – Membrane-associated Protein Assemblies, Machineries and Supercomplexes (project no. 450648163); Project 17 to M.B., CRC 1129 – Integrative Analysis of Pathogen Replication and Spread (project no. 240245660); Projects 5 to H.-G.K. and 20 to M.B.), by Hessian Ministry of Higher Education, Research, Science and the Arts (HMWK) as part of the cluster project EnABLE, and by the Excellence Initiative of the German Federal and State Governments EXC 3094 – 533751785 (Subcellular Architecture of LifE, SCALE, to B.T. and M.B.). The XL-MS work was funded by SAW (P70/2018) (F.L.) and DFG (LI 3260/6-1/) (F.L.)

## Autor contributions

AOK (conceptualization, manuscript writing, molecular modeling), YZ, KG, RRER, NY (crosslinking MS), HX, JPK, MAK, and BT (cryo-electron tomography, image processing), DG (immunofluorescence), SB, HGK (conceptualization, manuscript writing), FL (crosslinking MS, supervision, manuscript writing), MB (supervision, conceptualization, manuscript writing).

## Declaration of generative AI and AI-assisted technologies in the writing process

During the preparation of this work, the author(s) used ChatGPT, a generative AI language model, to review and improve phrasing, grammar, and readability. After using this tool, the author(s) reviewed and edited the content as needed and take full responsibility for the accuracy and integrity of the publication.

## STAR METHODS

### EXPERIMENTAL MODEL AND STUDY PARTICIPANT DETAILS

#### Cell Culture

HEK293 sfGFP cells were cultured in DMEM (Sigma) with 10% fetal bovine serum and 5 μg/ml tetracycline hydrochloride at 37°C with 5% CO_2_. HEK293-F cells were cultured in Freestyle medium (Thermo Fisher Scientific) at 37°C and 8% CO2 on an orbital shaker platform at 120×rpm. Expi293F cells were cultured in Expi293™ Expression Medium at 37°C and 8% CO_2_ on an orbital shaker platform at 100×rpm. Jurkat cells were cultured in RPMI Medium 1640 +D-glucose, Glutamine, sodium bicarbonate, sodium pyruvate (Catalog# A10491-01) supplemented with 10% FBS for dataset 3, both at 37 °C and 8% CO_2_ on an orbital shaker platform at 100×rpm. The HEK293 sfGFP cells were generated as described in *Xing et al.* (unpublished). In brief, pcDNA5-sfGFP was introduced using the HEK Flp-In T-Rex 293 system according to the manufacturer’s protocol (Invitrogen).

### METHOD DETAILS

#### Sample preparation for cryo-ET of HEK293 cells

Grids were prepared as described previously.^90^ The HEK293 cells used in this study were Flp-In T-Rex 293 cells stably expressing sfGFP (*Xing et al. unpublished*). Quantifoil R2/2 gold grids with 200 mesh were glow discharged and placed in 3.5 cm cell culture dishes. 2 ml cell suspensions (200,000 cells/ml) were added to the dishes and cultured for 5 hours before plunge freezing.

The grids were blotted on the back side for 6 seconds using a Leica EM GP2 plunger, set in an environment of 70% humidity and at a temperature of 37°C. The grids were quickly plunged into liquid ethane and then preserved in liquid nitrogen. The grids underwent FIB milling with the Aquilos FIB-SEM system (Thermo Scientific). They were coated with a protective layer of organometallic platinum via the gas injection system for 15 seconds. The lamella preparation was executed in stages, utilizing gallium ion-beam currents gradually reduced from 0.5 nA to 30 pA.

#### Data acquisition and tomogram reconstruction for HEK cells

Tilt series were acquired at 300 KV on a Titan Krios G2 (Thermo Scientific) and equipped with a Gatan BioQuantum-K3 imaging filter in counting mode, at ∼6K x 4K pixel dimensions, 2° tilt increment, tilt range of −64° to 52°, at a magnification of ×33,000, pixel size of 2.682 Å, a total dose of 150 e/Å2 per tilt series, and target defocus of −2.5 to −4.0 µm, using SerialEM software.^91^ Tilt series were aligned with patch-tracking in IMOD^92^ and 115 tomograms reconstructed at bin4 with SIRT-like filtering of 10 iterations. Additionally, 3D CTF-corrected tomograms were reconstructed using novaCTF.^93^

#### HEK293 NPC subtomogram averaging

Subtomogram averaging of NPC was performed as described previously (^94^, https://zenodo.org/records/5921012). Briefly, 247 NPC particles were manually picked from the filtered tomograms. With the picked particle coordinates, the refinement of NPC was performed using the 3D CTF-corrected tomograms at bin8 and bin4. The coordinates of the NR subunits were generated from the above refinement based on the C8 symmetry of NPC. Subsequently, the subtomograms of NR were extracted and aligned. The nuclear basket became visible at the NR subtomogram averaging map at this stage. An elliptical mask generated in Dynamo ^95^ covering the potential basket area was used to refine the basket filaments further. The stability of the filaments based on positions from subtomogram averaging was performed within cryoCAT package ^96^.

#### Macrophage NPC subtomogram averaging

To improve map quality of the NPC ring subunits two published human macrophage cryo-ET datasets (EMPIAR-12454 (HIV-infected) and EMPIAR-12457 (control))^41^ were combined. The NPC subtomogram averaging and alignment was performed as described in ^41^. In brief, NPC particles were extracted from the combined macrophage dataset from novaCTF-corrected tomograms ^93^ and subtomogram alignment and averaging was performed using novaSTA.^97^ First, a whole NPC average was obtained with imposed C8 symmetry. Next, subunit coordinates were extracted using C8 symmetry while also splitting the dataset into halfsets. The subunit positions were used to obtain an average structure of the asymmetric unit of the NPC. For the averages of individual NPC rings, particle positions were re-centered to the respective area (CR, IR, NR, LR, nuclear basket) based on their position in the subunit average and then aligned and averaged again. For generating a C8-symmetric composite NPC map, the individual ring maps were fitted into the asymmetric subunit average. The composite map was then created by applying C8 symmetry based on the coordinates used for splitting the initial whole NPC average into asymmetric units. The FSC calculations for each dataset were performed in Relion (v3.1) ^98^ and plotted using Python.

#### Nuclear interactome: Cell culture, nuclei isolation, crosslinking, protein digestion

Three different XL-MS data sets have been acquired for this study and merged for our structural modeling approach. They are referred to as dataset 1, dataset 2 and dataset 3 in the following. For dataset 1, we crosslinked isolated nuclei purified from HEK293-F cells. Cells were cultured and washed with PBS once, and stored at −80°C. For cellular lysis, the cell pellets were thawed in lysis buffer (25 mM HEPES pH 7.3, 150 mM NaCl, 0.1% IGEPAL CA-630, 0.5 mM DTT, 0.5 mM EDTA pH 8.0) supplemented with complete protease inhibitor EDTA-free cocktail, followed by passing through a G24 needle ten times and incubation on ice for 30 minutes. Nuclei were collected by centrifugation at 1,000xg at 4°C for 5 min. The nuclear pellet was washed twice by gentle resuspension in wash buffer (25 mM HEPES pH 7.3, 150 mM NaCl, 0.5 mM EDTA pH 8.0) supplemented with complete protease inhibitor EDTA-free cocktail and subsequent centrifugation. Then, the nuclear pellet was resuspended in wash buffer to an approximate protein concentration of 10 mg/ml. Crosslinking was performed using 4 mM Azide-A-DSBSO for 15 min at room temperature with constant mixing. The reaction was quenched with a final concentration of 20 mM Tris-HCl pH 8.0 for 30 min at room temperature with constant mixing. The crosslinking efficiency was monitored by SDS-PAGE. The nuclei were collected by centrifugation and resuspended in urea buffer (50 mM TEAB pH 8.55, 7.12 M Urea, 0.1% ProteaseMax).

Datasets 2 and 3 were derived from crosslinked homogenates of HEK and Jurkat cells, respectively. For the HEK dataset (dataset 2), HEK Expi293F cells were cultured in Expi293™ Expression Medium. Cells were washed with PBS and resuspended in lysis buffer (250mM sucrose, 10mM HEPES, 2mM EDTA, 2mM magnesium acetate tetrahydrate, pH 7.4) supplemented with complete protease inhibitor EDTA-free cocktail. Cells were then lysed by homogenization using dounce homogenizer at 1000×rpm until lysis efficiency reach 90%. Cell debris were spun down at 200×g for 5 min at 4 °C. The supernatant was collected and protein concentration was determined using the BCA protein assay kit (Thermo Fisher Scientific) according to the manufacturer’s instructions. The lysate was adjusted to a concentration of 10 mg/mL using lysis buffer and crosslinked with 2 mM Azide-A-DSBSO for 15 min at room temperature with constant mixing. The reaction was quenched with a final concentration of 20mM Tris-HCl pH 8.0 for 30 min at room temperature with constant mixing. The crosslinked samples were fractionated by differential centrifugation to reduce the proteome complexity. Pellets obtained at different centrifugation speed were collected separately and stored until further use.

For the Jurkat dataset (dataset 3), crosslinking was performed with 4 mM Azide-A-DSBSO for 30 min at room temperature with constant mixing. All other steps were the same as those described for dataset 2.

#### Protein digestion, cross-link enrichment and off-line HPLC fractionation

For dataset 1, nuclear lysis and DNA shearing were performed by sonication on ice (UP50H Ultrasonic Processor, Hielscher Ultrasonics GmbH) at 0.7 cycles and 60% amplitude for 1 min ten times, with at least 30 seconds in between with constant mixing at room temperature. Nuclear lysis was completed by prolonged incubation at room temperature with constant mixing for one hour total. Nuclear lysis efficiency was evaluated by microscopy. The nuclear extract was cleared by centrifugation at room temperature at 16,000xg for 3 min. The supernatant was collected and protein concentration was determined using the BCA protein assay kit (Thermo Fisher Scientific) according to the manufacturer’s instructions. After protein reduction (5 mM TCEP, 15 min RT) and alkylation (40 mM Chloroacetamide, 1 hour 37°C), proteins were digested in solution using LysC (Mass Spec Grade, Promega) at a ratio of 1:75 for 2.5 hours at RT. Subsequently, the samples were digested in 0.05% ProteaseMax and <2 M Urea/50 mM TEAB using Trypsin (Sequencing Grade Modified Trypsin, Promega) at a ratio of 1:75 for 22 hours at 37°C. After quenching with 1% formic acid (final concentration), peptides were desalted using C8 cartridges (Waters), quantified spectrophotometrically (DeNovix DS-11) and dried in vacuo.

For dataset 2 and dataset 3, each pellet was re-suspected in 8M urea dissolved in 50 mM TEAB for denaturation, and further reduced and alkylated. Proteins were digested with Lys-C at an enzyme-to-protein ratio of 1:75 (w/w) for 4 h at 37 °C. After diluting with 50 mM TEAB to a final concentration of 2 M urea, trypsin was added at an enzyme-to-protein ratio of 1:100 (w/w) for overnight at 37 °C. For dataset 2, the digestion was quenched by adding formic acid to a final concentration of 1%. Peptides were desalted with Sep-Pak C18 cartridges (Waters) according to the manufacturer’s protocol and dried in a vacuum concentrator. For dataset 3, the digested peptide mixture was directly subjected for further DSBSO enrichment.

For all datasets, the digested peptide mixtures were enriched by dibenzocyclooctyne (DBCO)-coupled sepharose beads. Briefly, peptides after digestion were added to the prewashed DBCO beads for incubation overnight at room temperature with constant mixing. After incubation, the beads were washed with water, followed by a washing step with 0.5% SDS at 37 °C for 15 min. Next, the beads were washed thrice with 0.5% SDS, thrice with 8 M urea in 50mM TEAB, thrice with 10% ACN and twice with water using ten bead volumes each. The crosslinked peptides were eluted with two bead volumes using 10 % (v/v) trifluoracetic acid (TFA) for 2 h at room temperature and then dried using a vacuum concentrator. Enriched crosslinks were further fractionated by size exclusion chromatography (SEC) followed by Hi-pH fractionation. SEC was performed using a SuperdexTM 30 Increase 3.2/300 column (GE Healthcare) on an Agilent 1260 Infinity II system and Hi-pH fractionation was performed using Phenomenex Gemini C18 column on an Agilent 1260 Infinity II UPLC system (for dataset 1 and 2) or Vanquish C18+, 1.5 μm, 2.1×50 mm column (Thermo Fisher Scientific) on Vanquish HPLC system (for dataset 3). Hi-pH fractions were subjected to LC-MS analysis.

#### LC-MS and data analysis

For the analysis of DSBSO-crosslinked peptides, collected fractions were analyzed by LC-MS using an UltiMate 3000 RSLC nano LC system coupled on-line to an Orbitrap Fusion Lumos mass spectrometer (Thermo Fisher Scientific) (for dataset 1 and 2) or an Orbitrap Exploris 480 mass spectrometer (Thermo Fisher Scientific) (for dataset 3). Reversed phase separation was performed with an in-house packed C18 analytical column (Poroshell 120 EC-C18, 2.7µm, Agilent Technologies). Fractions were run with 3 h LC gradients. The following MS parameters were applied: MS resolution 120,000; MS2 resolution 60,000; charge state 4-8 enable for MS2; MS2 isolation window, 1.6 m/z; stepped normalized collision energy, 19-25-30%; FAIMS compensation voltages were set to −50, −60, and −75. Data analysis was performed using Scout (https://github.com/theliulab/Scout)^99^ and pLink (https://github.com/pFindStudio/pLink3) with the following parameters: minimum peptide length 6; maximal peptide length 60; missed cleavages 3. Fixed modification was set as cysteine carbamidomethylation, and variable modification was set as methionine oxidation. The DSBSO cross-linker mass was specified as 308.0388 Da (short arm: 54.0106 Da, long arm: 236.0177 Da). For datasets 2 and 3, protein N-terminal acetylation was included as an additional variable modification. Precursor mass tolerance was set to 10 ppm, and fragment mass tolerance to 20 ppm. MS2 spectra were searched against a reduced target-decoy Swiss-Prot human database, which was derived from the proteomic measurements of all fractions.

Cross-links were extracted from result tables generated by the two software programs. For dataset 1, Scout results were filtered at 1% FDR at the peptide-pair level. Plink3 results were filtered at 1% FDR at the peptide-pair level for unique cross-links and for shared cross-links (those containing shared peptides) at 1% FDR at the peptide-pair level. The results from both search engines were combined for the final report. For datasets 2 and 3, pLink3 results were filtered at a 1% FDR at the PPI level for unique cross-links. All other filtering parameters were applied as described for dataset 1.

#### Modeling using AlphaFold 3

Models were generated using AlphaFold 3 server or standalone version of AlphaFold 3. Structural similarity of the five generated models for the given complex was assessed using USalign^100^. The highest-ranked model according to AlphaFold 3 was selected.

#### Modeling the TPR homodimer

To reduce computational complexity while minimizing bias, the crosslinks were grouped into sets based on equivalence, i.e., interactions defining equivalent arrangements between TPR coiled-coil fragments. For each modeling run, we randomly selected five crosslinks from different groups. Models were then generated for all possible combinations of two or three crosslinks from each reduced set using AF3x.^101^ For each combination, 50 independent modeling runs were performed with different random seeds. The resulting models were ranked based on their total pLDDT scores and the number of satisfied crosslinks.

#### Modeling of the tetramer

The TPR tetrameric coiled-coil bundle spanning residues 1–320 and 660–1460 was modeled using ColabFold^60^ with the five top models of the TPR homodimers as custom templates.

#### Quality assessment

The quality of the AF3 and AF3x models was assessed using the scores provided by AlphaFold: the predicted local-distance difference test (pLDDT), a metric for the local accuracy, and the Predicted Aligned Error (PAE), which assesses the relative orientation of the proteins and protein domains. The quality of the models was also assessed by agreement with XL-MS data.

#### Utilization of different cryo-EM maps

We used different maps for model building and validation to capitalize on the highest available resolution. Systematic fitting and construction of the initial IR models were performed using the IR region of the published cryo-ET map of the NPC from HEK293 cells (EMD-14322^18^), representing the constricted state and best possible resolution for the IR. Systematic fitting of NR components and model building of the NR and basket were performed using the NR region of the cryo-ET map of the human macrophage NPC, representing the dilated state and best possible resolution for the NR. In addition, cryo-EM maps from *X. laevis* (EMD-31065, EMD-32394 ^40^) and *S. cerevisiae* (EMD-24231 ^16^) were used for validation, both representing isolated, constricted NPCs. While the *X. laevis* map enables the assessment of secondary structure of the NR, the yeast map adds value for evolutionary comparisons. To generate the model of the entire human NPC, subcomplexes were fitted into the cryo-ET map of the human macrophage NPC, representing the dilated state, and subsequently further refined.

#### Systematic fitting

To assign atomic structures within the cryo-ET density maps, we employed a systematic fitting approach as previously described.^43,63^ Prior to fitting, all high-resolution atomic models were filtered to a resolution between 10 and 30 Å to match the resolution of the ET data. These simulated maps were then globally fitted into the EM maps using UCSF Chimera^102^ via Assembline.^63^ Each fitting run began with 100,000 random placements of the model, with a constraint requiring that at least 30% of the model map overlapped with the cryo-ET density contour at a defined low threshold. For each model, this process typically produced between 1,000 and 20,000 non-redundant fits upon clustering. The cross-correlation about the mean (cam score), equivalent to the Pearson correlation coefficient, calculated in UCSF Chimera^102^, was used as the primary metric to assess the quality of each atomic model fit. To evaluate statistical significance, cam scores were converted to z-scores using Fisher’s transformation, centered, and used to calculate two-sided *p*-values. The standard deviation required for this step was derived from an empirical null distribution generated from all nonredundant fits and modeled using the *fdrtool* package in R.^103^ To account for multiple comparisons, the resulting *p*-values were adjusted using the Benjamini-Hochberg method.^104^

#### Generation of the model of the nuclear basket

We built an initial structural model of the IR based on a previously published model and a cryo-ET map of the constricted state NPC.^18^ Newly generated AF3 models of the previously assigned NUPs and their subcomplexes were superposed onto their counterparts in the published structure. AF3 models of the newly identified NUP subcomplexes, NUP155–SMPD4, TMEM209 CTD–2×(NUP98–93–62–58–54), and NUP93–NUP35×2, were positioned through systematic fitting using Assembline^63^. This model was then fitted as a rigid body into the cryo-ET map of the human macrophage NPC in the dilated conformation.

The NR and nuclear basket were modeled using the cryo-ET map of the human macrophage NPC and the published model^18^. AF3 models of the previously assigned NUPs and their subcomplexes were superposed onto the corresponding components in the published structure fitted into the cryo-ET map of the human macrophage NPC. AF3 models of the newly identified subcomplexes, NUP85–SEH1–NUP43–GANP–NUP153, NUP205–NUP93–GANP and NUP96–NUP107–TPR×2–NUP153-GANP, were positioned through systematic fitting using Assembline.^63^ AF3 models of the NUP85–SEH1–NUP43–GANP–NUP153–Centrin-2–ENY2×2 subcomplex was placed by superposition with the NUP85–SEH1–NUP43–GANP–NUP153 model, and then used as a reference to position AF3 model of the GANP–ENY2×4 subcomplex. Models of the outer TPR insertion region TPR×2–NUP153–GANP and the TPR tetramer were manually placed and locally refined in ChimeraX.^105^

Models of the CR and LR were generated using the same approach as for the NR. The second copy of the NUP214 subcomplex was positioned in the CR through systematic fitting using Assembline.^63^

The NPC model was subsequently refined using Assembline^63^, built on the Integrative Modeling Platform (IMP v2.15)^106^ and the Python Modeling Interface (PMI).^107^ For refinement, the models were divided into smaller segments, each treated as rigid body in a Cα-only representation. Inter-domain and inter-subunit interfaces were restrained using elastic distance networks derived from AF3 models of the NUP subcomplexes. The Assembline^63^ refinement step was applied to optimize map fitting, minimize steric clashes, and ensure connectivity between sequentially adjacent domains. The scoring function combined: (i) EM fit restraints, (ii) clash penalties (SoftSpherePairScore in IMP), (iii) connectivity restraints between sequentially adjacent domains, and (iv) elastic network restraints from AF3-modeled NUP subcomplexes. Final atomic-resolution models were obtained by back-mapping the refined Cα-only representation to the original AF3 models.

AF3 models of the TPR C-terminal coiled-coil, NUP50×2–NUP153, the NUP50 folded domain, GANP–PCID2–DSS1–DDX39B, the zinc-finger domains of NUP153, NUP214 mRNA export platform, and the Ran-binding and zinc-finger domains of NUP358, as well as all disordered regions, were placed arbitrarily. Fragments within individual protein chains were then connected using ISOLDE.^108^

#### Visualization

Figures were prepared using ChimeraX.^105^ Crosslink diagrams were prepared using xiNET ^109^. Crosslinks mappings onto the models were done using XMAS plugin to ChimeraX.^110^

#### Immunofluorescence

For immunofluorescence staining, adherent human embryonic kidney (HEK293) cells were cultured on laminin-coated ibidi 18-well dishes (Roche, 11243217001). Cells were fixed with 4% paraformaldehyde in PBS for 20 min at room temperature (RT), permeabilized with 0.1% Triton X-100 in PBS for 5 min at RT, and blocked for 1 h at RT in blocking buffer (2% BSA and 0.1% Tween-20 in PBS). Cells were incubated with a rabbit polyclonal antibody against human SMPD4 diluted 1:75 in 1 % BSA in PBS for 1 h at RT, followed by incubation with a goat anti-rabbit IgG secondary antibody conjugated to Alexa Fluor 680 diluted 1:1000 in 1 % BSA in PBS for 1 h at RT. Nuclei were stained with DAPI (0.25 µg/mL in PBS) for 5 min at RT. The samples were sealed with Prolonged Antifade Glass mounting medium (Invitrogen). Images were acquired on an inverted Stellaris 5 confocal microscope (Leica) using a 63×/1.4 NA oil-immersion objective and processed in ImageJ2.^111^

### QUANTIFICATION AND STATISTICAL ANALYSIS

The details of quantification and all statistical analyses have been described in the relevant sections of the METHOD DETAILS.

