## Supplementary Data for "How the TREX-2 complex associates with the nuclear pore"

### Supplementary Figures

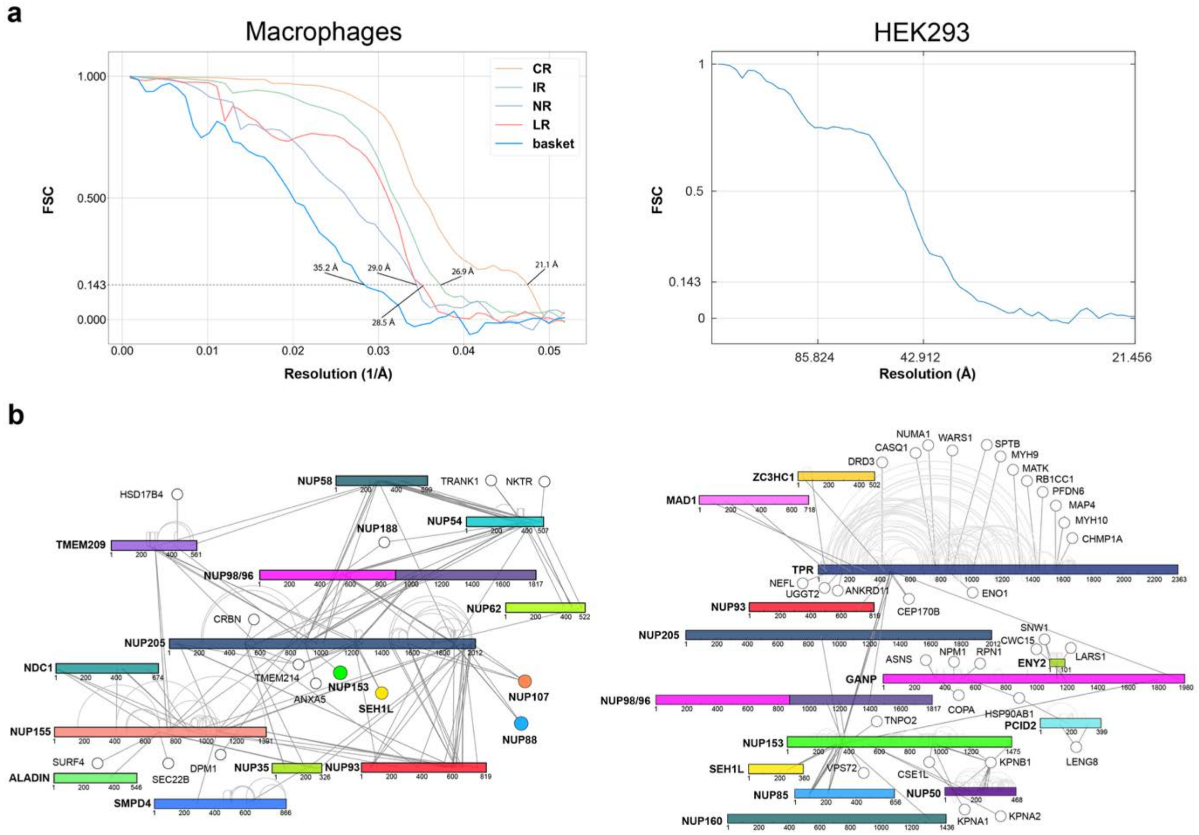

**Fig. S1. Validation of cryo-ET and XL-MS data. (a)** FSC curves of the cryo-ET maps of the different NPC substructures from macrophages[41], and the basket filaments for cryo-ET map from HEK293 cells (this study). **(b)** Crosslink map of the IR (*left*) and the TREX-2 subcomplex and nuclear basket (*right*). Key proteins show as primary structure plots.

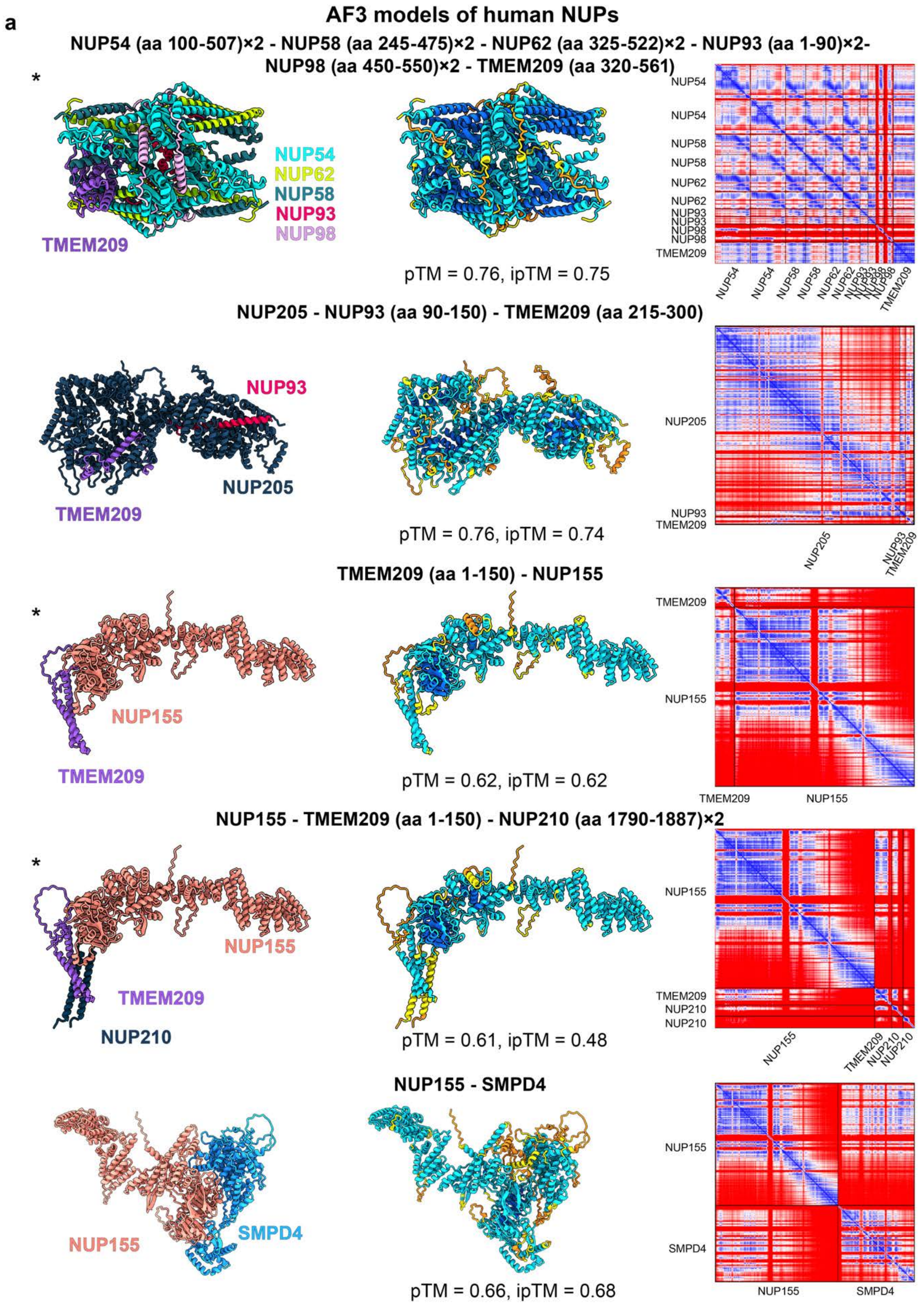
a
AF3 models of human NUPs
NUP54 (aa 100-507)×2 - NUP58 (aa 245-475)×2 - NUP62 (aa 325-522)×2 - NUP93 (aa 1-90)×2-
NUP98 (aa 450-550)×2 - TMEM209 (aa 320-561)
*
NUP54
NUP62
NUP58
NUP93
NUP98
TMEM209
pTM = 0.76, ipTM = 0.75
NUP205 - NUP93 (aa 90-150) - TMEM209 (aa 215-300)
NUP93
NUP205
TMEM209
pTM = 0.76, ipTM = 0.74
TMEM209 (aa 1-150) - NUP155
*
NUP155
TMEM209
pTM = 0.62, ipTM = 0.62
NUP155 - TMEM209 (aa 1-150) - NUP210 (aa 1790-1887)×2
*
NUP155
TMEM209
NUP210
pTM = 0.61, ipTM = 0.48
NUP155 - SMPD4
NUP155
SMPD4
pTM = 0.66, ipTM = 0.68

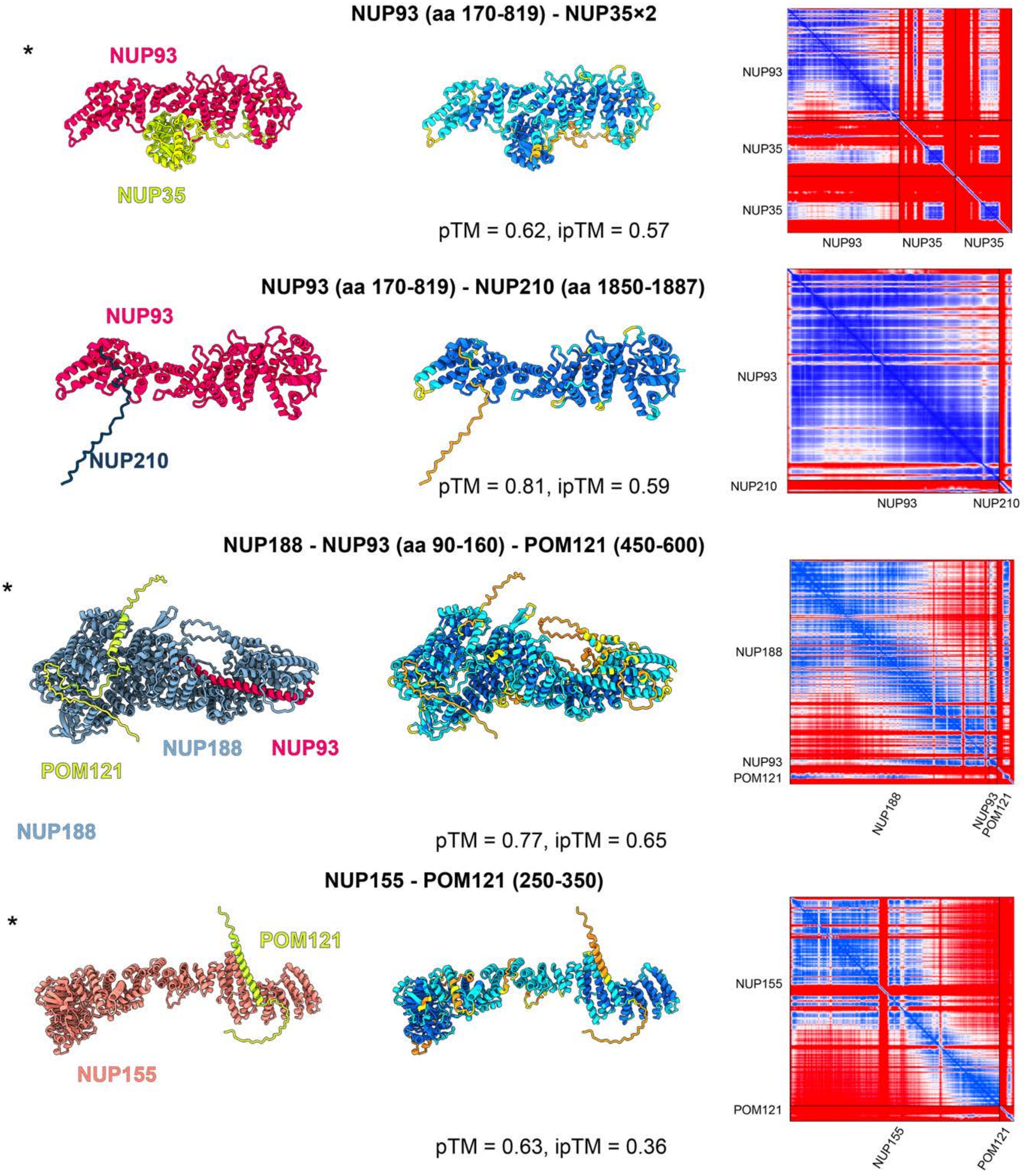
NUP93 (aa 170-819) - NUP35×2
*
NUP93
NUP35
pTM = 0.62, ipTM = 0.57
NUP93
NUP35
NUP35
NUP93 NUP35 NUP35
NUP93 (aa 170-819) - NUP210 (aa 1850-1887)
NUP93
NUP210
pTM = 0.81, ipTM = 0.59
NUP93
NUP210
NUP93 NUP210
NUP188 - NUP93 (aa 90-160) - POM121 (450-600)
*
POM121
NUP188
NUP93
NUP188
pTM = 0.77, ipTM = 0.65
NUP188
NUP93
POM121
NUP188
NUP93
POM121
NUP155 - POM121 (250-350)
*
POM121
NUP155
pTM = 0.63, ipTM = 0.36
NUP155
POM121
NUP155
POM121

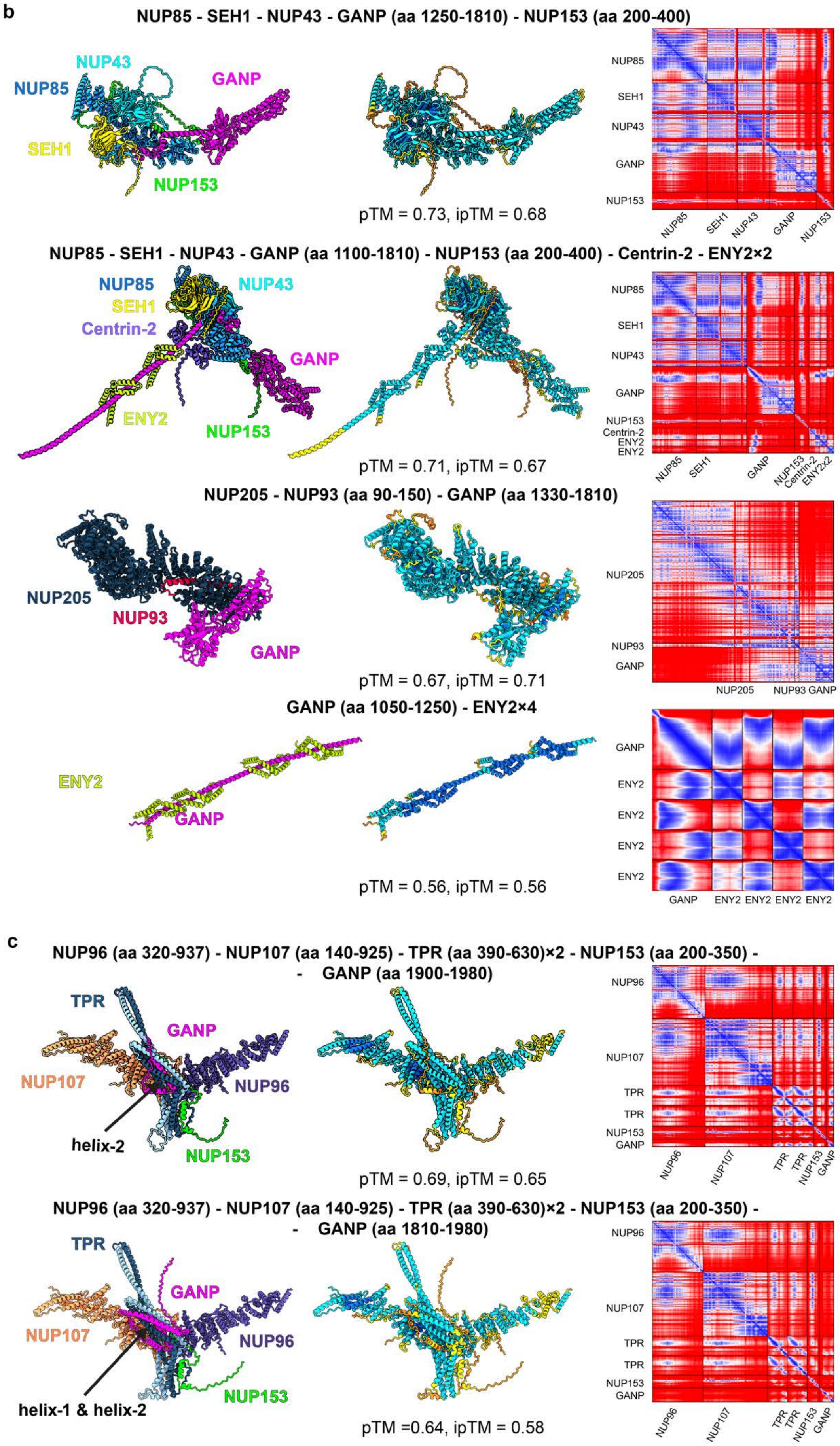
b
NUP85 - SEH1 - NUP43 - GANP (aa 1250-1810) - NUP153 (aa 200-400)
NUP43
NUP85
GANP
SEH1
NUP153
pTM = 0.73, ipTM = 0.68
NUP85
SEH1
NUP43
GANP
NUP153
NUP85 - SEH1 - NUP43 - GANP (aa 1100-1810) - NUP153 (aa 200-400) - Centrin-2 - ENY2×2
NUP85
NUP43
SEH1
Centrin-2
GANP
ENY2
NUP153
pTM = 0.71, ipTM = 0.67
Centrin-2
ENY2
ENY2x2
NUP205 - NUP93 (aa 90-150) - GANP (aa 1330-1810)
NUP205
NUP93
GANP
pTM = 0.67, ipTM = 0.71
GANP (aa 1050-1250) - ENY2×4
ENY2
GANP
pTM = 0.56, ipTM = 0.56
c
NUP96 (aa 320-937) - NUP107 (aa 140-925) - TPR (aa 390-630)×2 - NUP153 (aa 200-350) -
- GANP (aa 1900-1980)
TPR
GANP
NUP107
NUP96
helix-2
NUP153
pTM = 0.69, ipTM = 0.65
NUP96 (aa 320-937) - NUP107 (aa 140-925) - TPR (aa 390-630)×2 - NUP153 (aa 200-350) -
- GANP (aa 1810-1980)
TPR
GANP
NUP107
NUP96
helix-1 & helix-2
NUP153
pTM =0.64, ipTM = 0.58

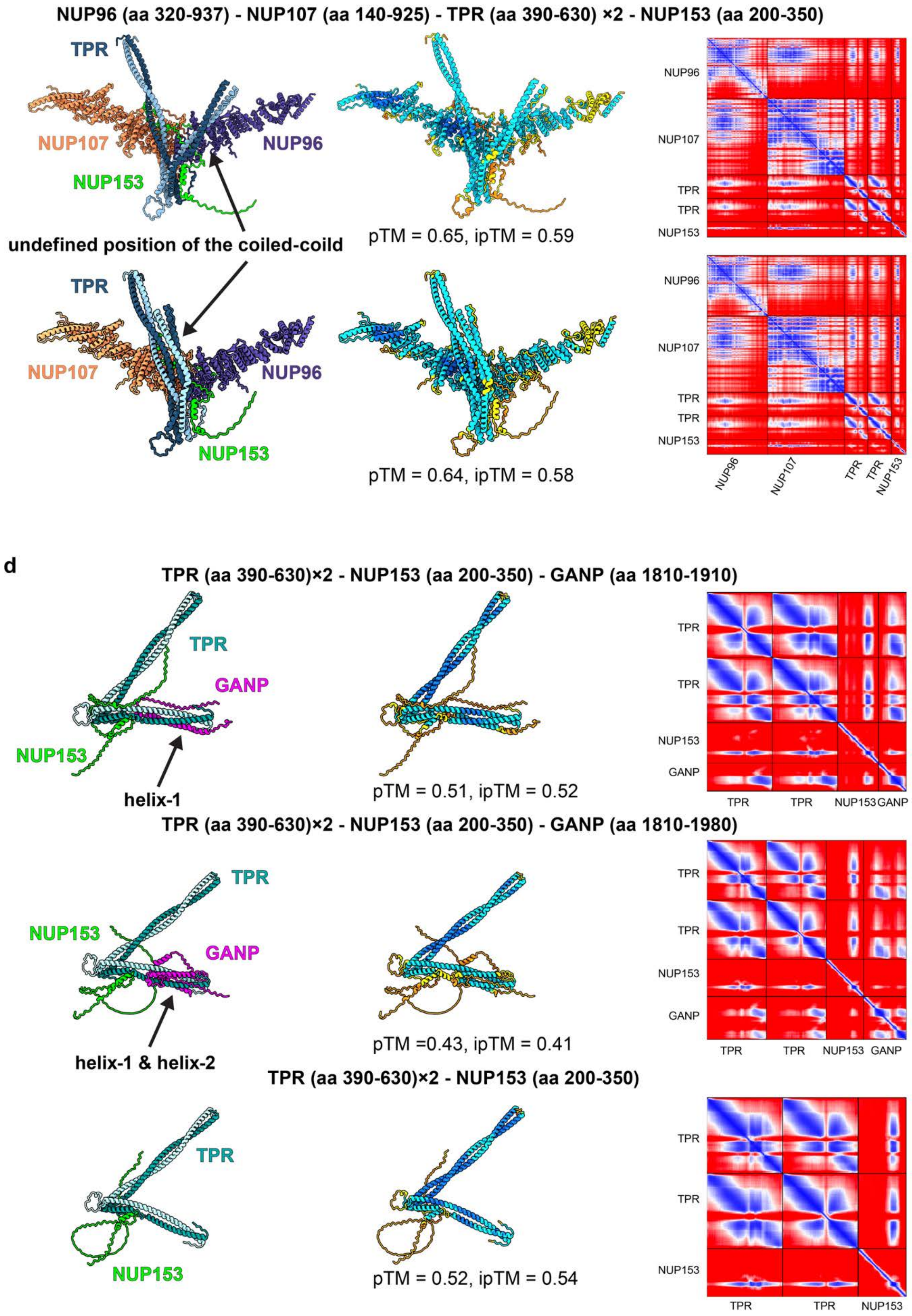
NUP96 (aa 320-937) - NUP107 (aa 140-925) - TPR (aa 390-630) ×2 - NUP153 (aa 200-350)
TPR
NUP107
NUP96
NUP153
undefined position of the coiled-coild
pTM = 0.65, ipTM = 0.59
TPR
NUP107
NUP96
NUP153
pTM = 0.64, ipTM = 0.58
NUP96
NUP107
TPR
TPR
NUP153
d
TPR (aa 390-630)×2 - NUP153 (aa 200-350) - GANP (aa 1810-1910)
TPR
GANP
NUP153
helix-1
pTM = 0.51, ipTM = 0.52
TPR
TPR
NUP153
GANP
TPR (aa 390-630)×2 - NUP153 (aa 200-350) - GANP (aa 1810-1980)
TPR
NUP153
GANP
helix-1 & helix-2
pTM =0.43, ipTM = 0.41
TPR (aa 390-630)×2 - NUP153 (aa 200-350)
TPR
NUP153
pTM = 0.52, ipTM = 0.54

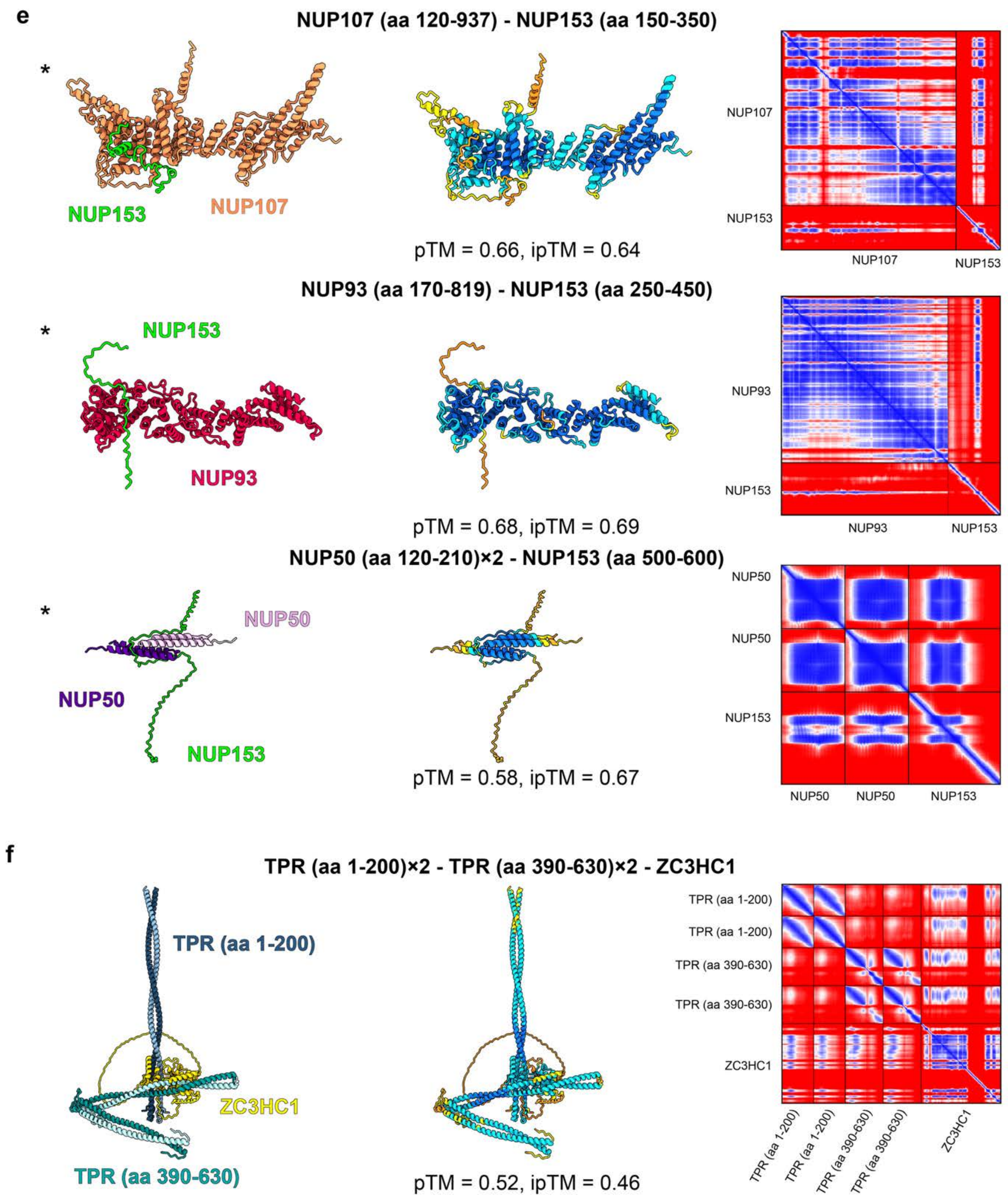
e
NUP107 (aa 120-937) - NUP153 (aa 150-350)
*
NUP153
NUP107
pTM = 0.66, ipTM = 0.64
NUP107
NUP153
NUP107
NUP153
NUP93 (aa 170-819) - NUP153 (aa 250-450)
*
NUP153
NUP93
pTM = 0.68, ipTM = 0.69
NUP93
NUP153
NUP93
NUP153
NUP50 (aa 120-210)×2 - NUP153 (aa 500-600)
*
NUP50
NUP50
NUP153
pTM = 0.58, ipTM = 0.67
NUP50
NUP50
NUP153
NUP50
NUP50
NUP153
f
TPR (aa 1-200)×2 - TPR (aa 390-630)×2 - ZC3HC1
TPR (aa 1-200)
ZC3HC1
TPR (aa 390-630)
pTM = 0.52, ipTM = 0.46
TPR (aa 1-200)
TPR (aa 1-200)
TPR (aa 390-630)
TPR (aa 390-630)
ZC3HC1
TPR (aa 1-200)
TPR (aa 1-200)
TPR (aa 390-630)
TPR (aa 390-630)
ZC3HC1

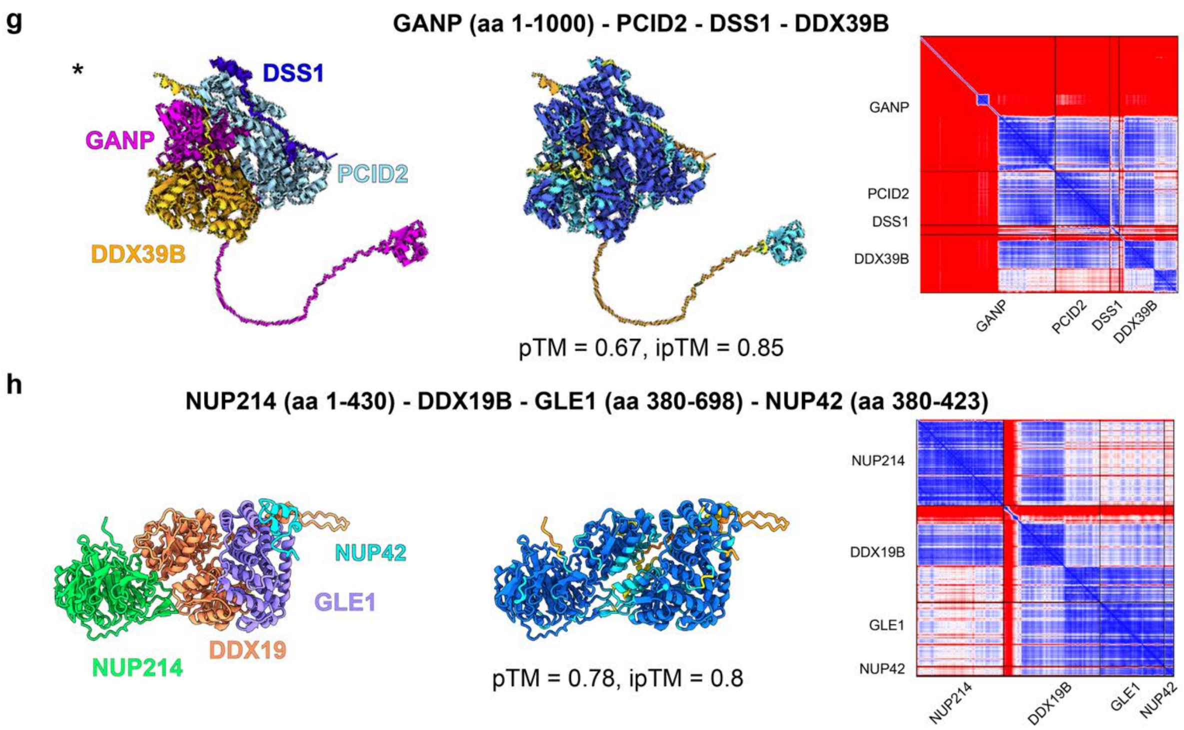
g
GANP (aa 1-1000) - PCID2 - DSS1 - DDX39B
*
DSS1
GANP
PCID2
DDX39B
pTM = 0.67, ipTM = 0.85
GANP
PCID2
DSS1
DDX39B
GANP
PCID2
DSS1
DDX39B
h
NUP214 (aa 1-430) - DDX19B - GLE1 (aa 380-698) - NUP42 (aa 380-423)
NUP42
GLE1
DDX19
NUP214
pTM = 0.78, ipTM = 0.8
NUP214
DDX19B
GLE1
NUP42
NUP214
DDX19B
GLE1
NUP42

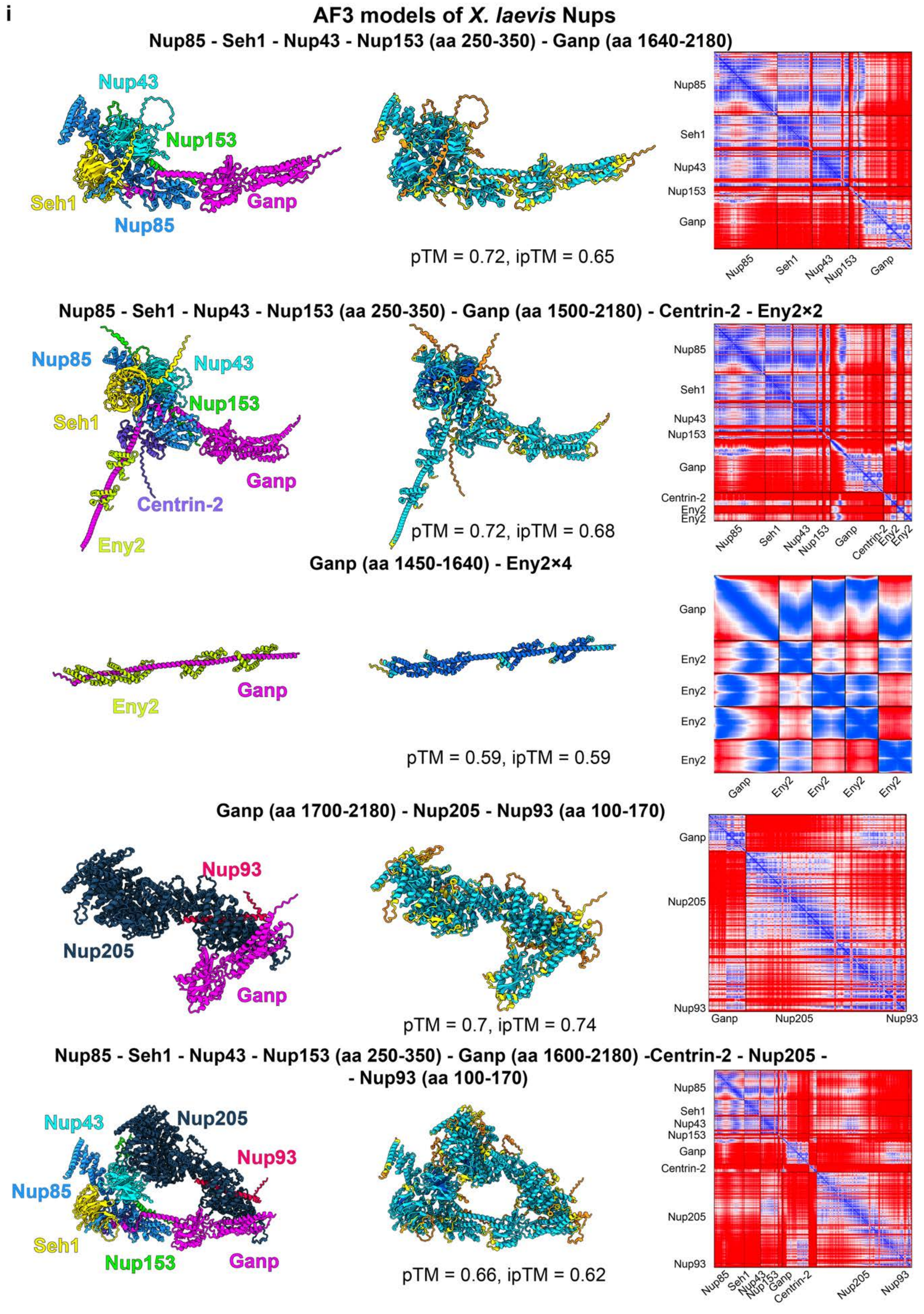
i
AF3 models of X. laevis Nups
Nup85 - Seh1 - Nup43 - Nup153 (aa 250-350) - Ganp (aa 1640-2180)
Nup43
Nup153
Seh1
Ganp
Nup85
pTM = 0.72, ipTM = 0.65
Nup85
Seh1
Nup43
Nup153
Ganp
Nup85 - Seh1 - Nup43 - Nup153 (aa 250-350) - Ganp (aa 1500-2180) - Centrin-2 - Eny2×2
Nup85
Nup43
Nup153
Seh1
Ganp
Centrin-2
Eny2
pTM = 0.72, ipTM = 0.68
Nup85
Seh1
Nup43
Nup153
Ganp
Centrin-2
Eny2
Eny2
Ganp (aa 1450-1640) - Eny2×4
Eny2
Ganp
pTM = 0.59, ipTM = 0.59
Ganp
Eny2
Eny2
Eny2
Eny2
Ganp (aa 1700-2180) - Nup205 - Nup93 (aa 100-170)
Nup93
Nup205
Ganp
pTM = 0.7, ipTM = 0.74
Ganp
Nup205
Nup93
Nup85 - Seh1 - Nup43 - Nup153 (aa 250-350) - Ganp (aa 1600-2180) -Centrin-2 - Nup205 -
- Nup93 (aa 100-170)
Nup43
Nup205
Nup93
Nup85
Seh1
Nup153
Ganp
pTM = 0.66, ipTM = 0.62
Nup85
Seh1
Nup43
Nup153
Ganp
Centrin-2
Nup205
Nup93

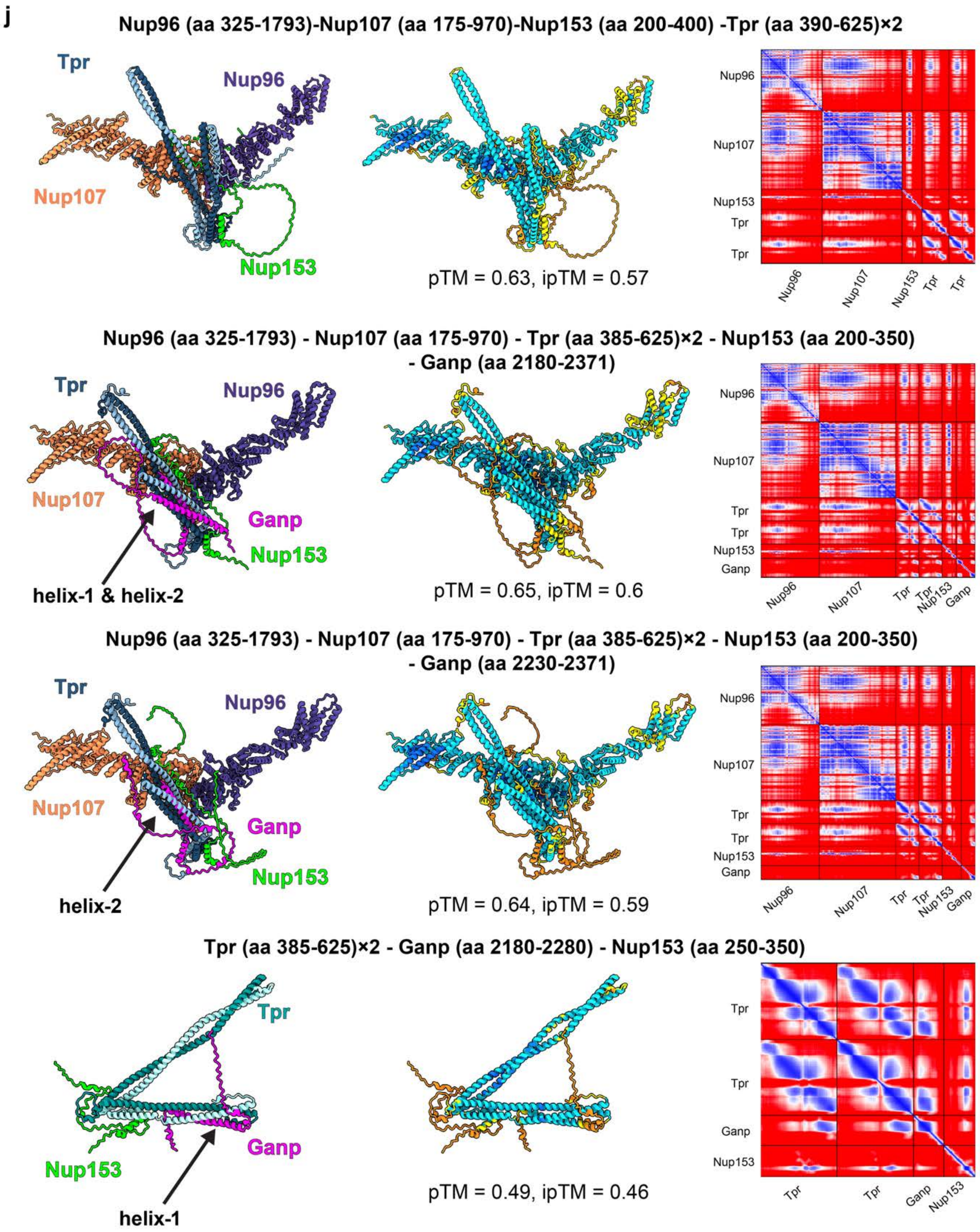
j
Nup96 (aa 325-1793)-Nup107 (aa 175-970)-Nup153 (aa 200-400) -Tpr (aa 390-625)×2
Tpr
Nup96
Nup107
Nup153
pTM = 0.63, ipTM = 0.57
Nup96 (aa 325-1793) - Nup107 (aa 175-970) - Tpr (aa 385-625)×2 - Nup153 (aa 200-350) - Ganp (aa 2180-2371)
Tpr
Nup96
Nup107
Ganp
Nup153
helix-1 & helix-2
pTM = 0.65, ipTM = 0.6
Nup96 (aa 325-1793) - Nup107 (aa 175-970) - Tpr (aa 385-625)×2 - Nup153 (aa 200-350) - Ganp (aa 2230-2371)
Tpr
Nup96
Nup107
Ganp
Nup153
helix-2
pTM = 0.64, ipTM = 0.59
Tpr (aa 385-625)×2 - Ganp (aa 2180-2280) - Nup153 (aa 250-350)
Tpr
Ganp
Nup153
helix-1
pTM = 0.49, ipTM = 0.46

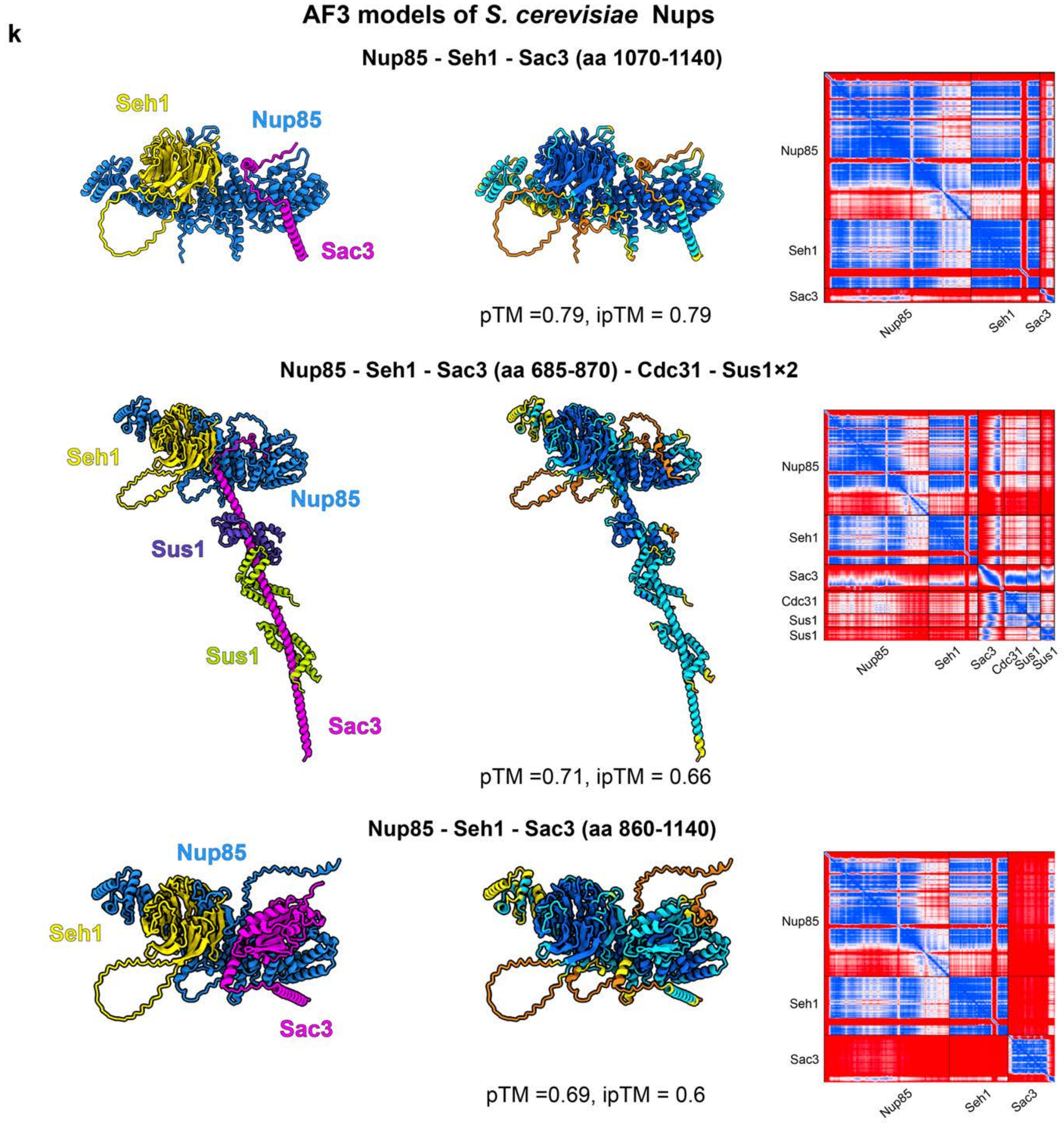
k
AF3 models of S. cerevisiae Nups
Nup85 - Seh1 - Sac3 (aa 1070-1140)
Seh1
Nup85
Sac3
pTM =0.79, ipTM = 0.79
Nup85
Seh1
Sac3
Nup85 - Seh1 - Sac3 (aa 685-870) - Cdc31 - Sus1×2
Seh1
Nup85
Sus1
Sus1
Sac3
pTM =0.71, ipTM = 0.66
Nup85
Seh1
Sac3
Cdc31
Sus1
Sus1
Nup85 - Seh1 - Sac3 (aa 860-1140)
Nup85
Seh1
Sac3
pTM =0.69, ipTM = 0.6
Nup85
Seh1
Sac3

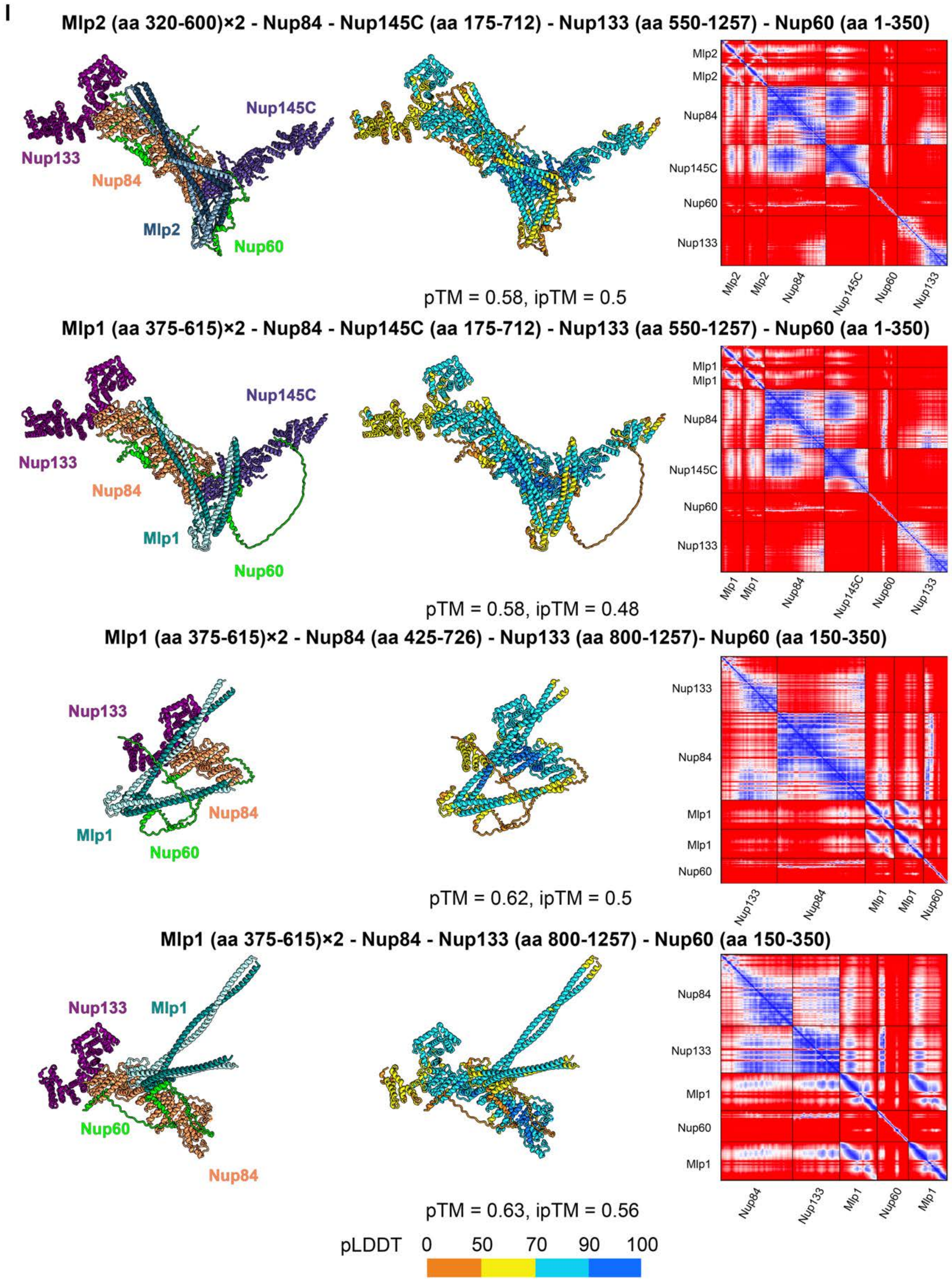

**Fig. S2. AF3 models of NUP subcomplexes used throughout the study.** Panels are organized into human **(a-h)**, *X. laevis* **(i-j)** and yeast **(k-l)** subcomplexes. In all panels, the model is shown colored by NUP composition (*left*) and colored by local confidence (pLDDT score, *middle*). Color bar at the bottom of the figure. The Predicted Aligned Error (PAE) plot is shown on the right, displaying the expected positional error between residue pairs. In the PAE plots, both axes represent the sequence from N- to C-terminus, with colors indicating confidence in relative positions (blue = high confidence, red = low confidence). In the images of the models marked with an asterisk, longer disordered fragments are omitted for clarity. **(a)** IR NUPs. **(b)** GANP. TPR insertion region of the outer **(c)** and inner copy **(d).** **(e)** NUP153. **(f)** ZC3HC1. Helicases of TREX-2 **(g)** and NUP214-subcomplexes **(h).** **(i)** Ganp. **(j)** Tpr insertion region. **(k)** Sac3. **(l)** Mlp1/2 insertion region. Structural comparison of the AF3 models generated for the given subcomplex from a single run revealed that they were highly similar (Table S2).

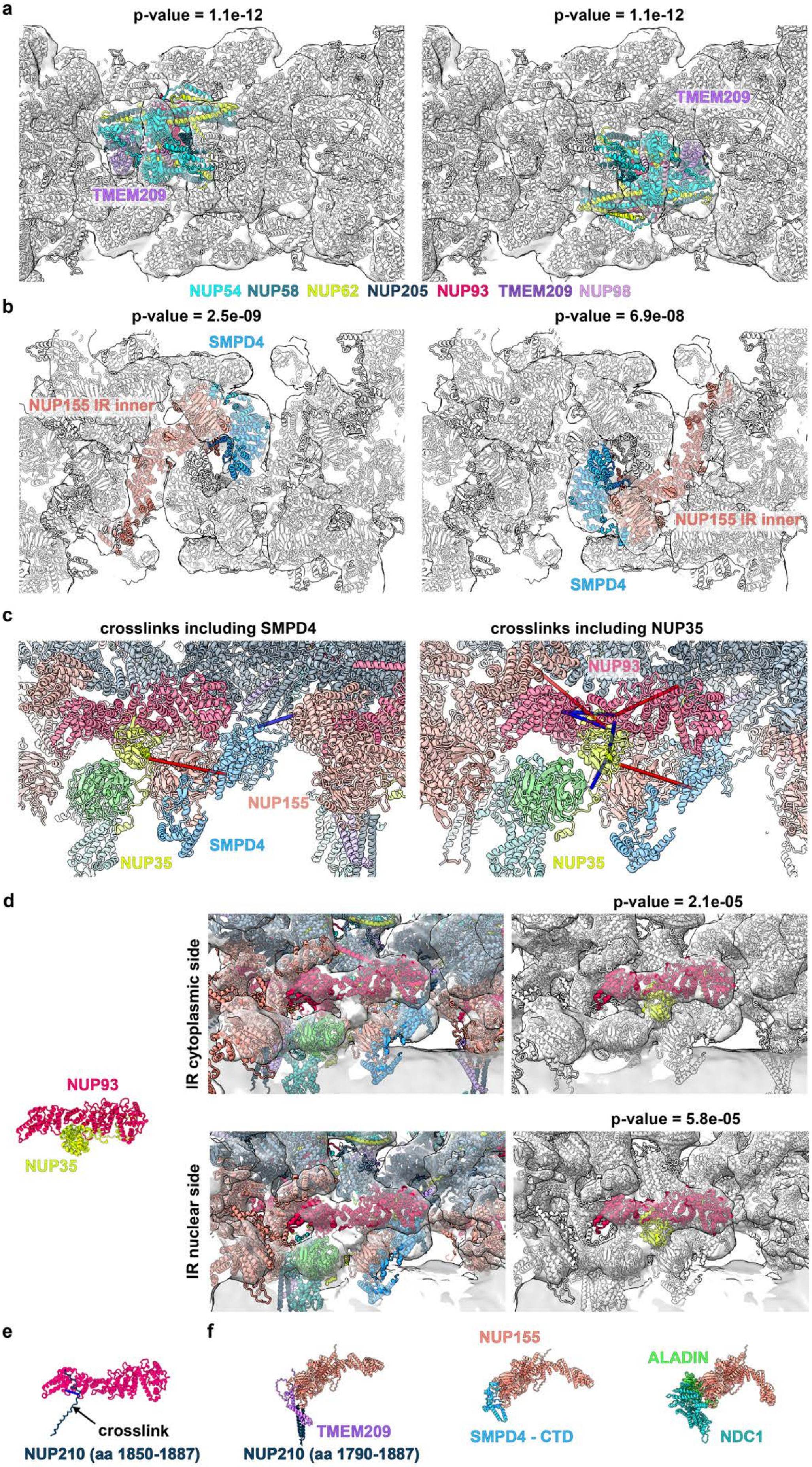
a
p-value = 1.1e-12
TMEM209
p-value = 1.1e-12
TMEM209
NUP54 NUP58 NUP62 NUP205 NUP93 TMEM209 NUP98
b
p-value = 2.5e-09
SMPD4
NUP155 IR inner
p-value = 6.9e-08
NUP155 IR inner
SMPD4
c
crosslinks including SMPD4
NUP155
SMPD4
NUP35
crosslinks including NUP35
NUP93
NUP35
d
p-value = 2.1e-05
IR cytoplasmic side
NUP93
NUP35
p-value = 5.8e-05
IR nuclear side
e
crosslink
NUP210 (aa 1850-1887)
f
TMEM209
NUP210 (aa 1790-1887)
NUP155
SMPD4 - CTD
ALADIN
NDC1

1325

**Fig. S3. Modeling of the human NPC IR. (a)** Systematic fitting of the AF3 model of the TMEM209 CTD-(NUP54-NUP58-NUP62-NUP93-NUP98)×2 subcomplex (shown color-coded) into the IR region of the cryo-ET map of the human NPC (EMD-14322[18]). Top-scoring positions in the IR are shown in context of the cryo-ET map and previously assigned proteins (white) of the IR as seen from the central channel; *p*-values are indicated. **(b)** Same as (a) but for AF3 model of NUP155-SMPD4 subcomplex. **(c-d)** Positioning of NUP93-NUP35 subcomplex. **(c)** Structural model of the IR highlighting crosslinks outgoing from SMPD4 (*left*) and the structured domain of NUP35 (*right*). Crosslinks shown as pseudo-bonds between the Cα atoms of lysines; satisfied crosslinks ( $\leq$ 40 Å) blue, violated red. **(d)** AF3 model of NUP93-NUP35×2 (*left*; Fig. S2); Structural model of the IR (*center*) without NUP35 fitted into the IR of the human NPC cryo-ET map (EMD-14322 [18]) as seen from the CR (*top*) and NR (*bottom*); Same as center panel but highlighting the two top-scoring positions of systematic fitting of the same subcomplex (colored) into the cytoplasmic and nuclear part of the IR; *p*-values are indicated. **(e)** AF3 model of NUP93-NUP210 supported by one satisfied crosslink. **(f)** AF3 models of NUP155 bound to TMEM209-NUP210 (*left*), SMPD4 CTD (*center*) and ALADIN-NDC1 (*right*), respectively. As a control, other potential interactions involving NUP155 were assessed using AF3. However, no additional positive interactions were detected (Table S3).

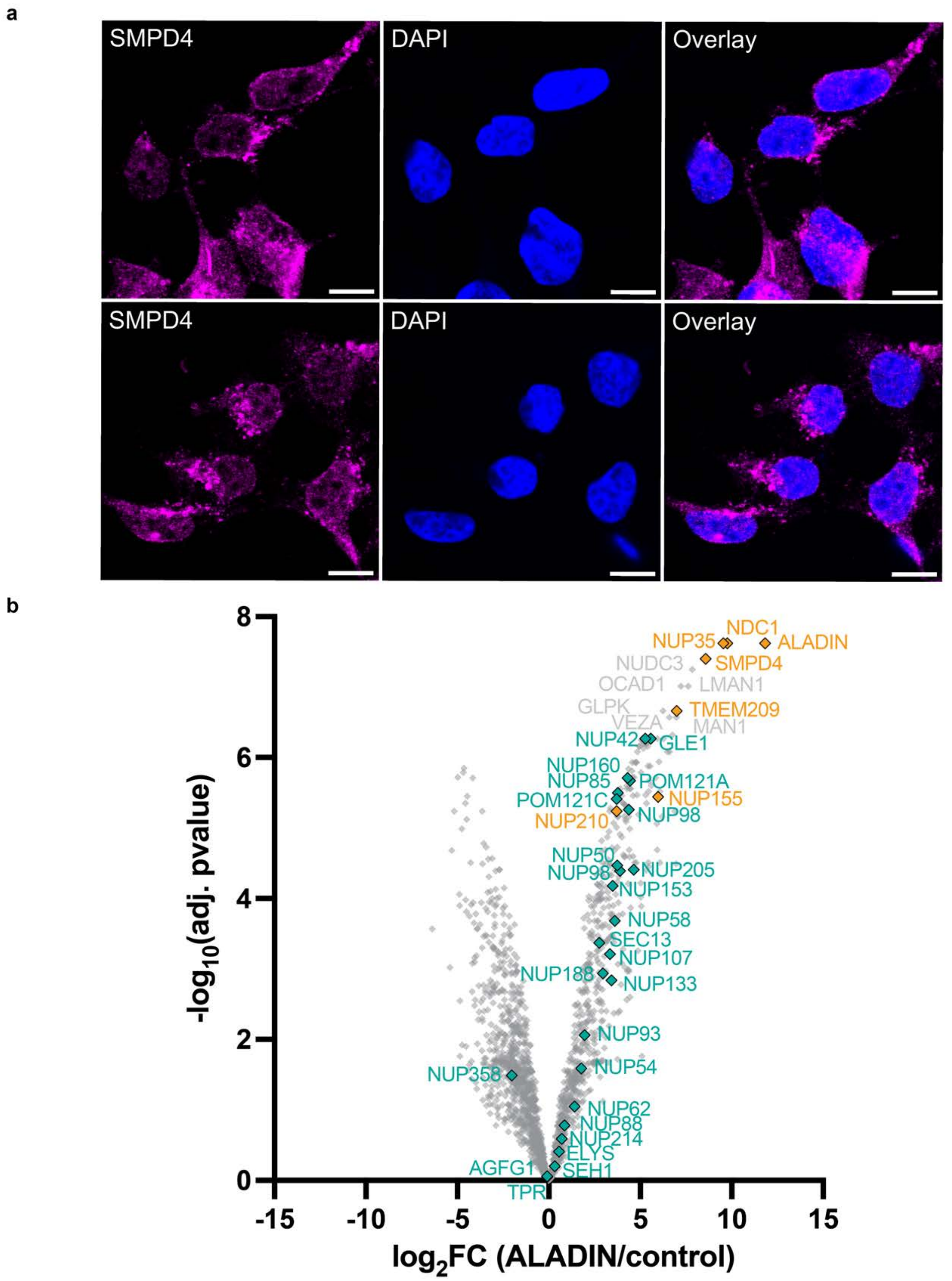

**Fig. S4. Validation of localization od SMPD4 and TMEM209**. **(a)** Two representative images of SMPD4 immunofluorescence acquired by confocal microscopy in HEK293 cells. Nuclei labeled with DAPI (blue). Single-channels and merged overlays are shown. Scale bars, 10 µm. **(b)** Volcano plot depicting interactors of ALADIN detected in a proximity labeling experiment of BirA-Aladin vs. negative control (BirA-tagged NLS-NES-Dendra). The dataset was previously published[18] and reanalyzed here. NUPs in green; direct interactors of ALADIN, TMEM209, SMPD4 and NUP210 are depicted in orange.

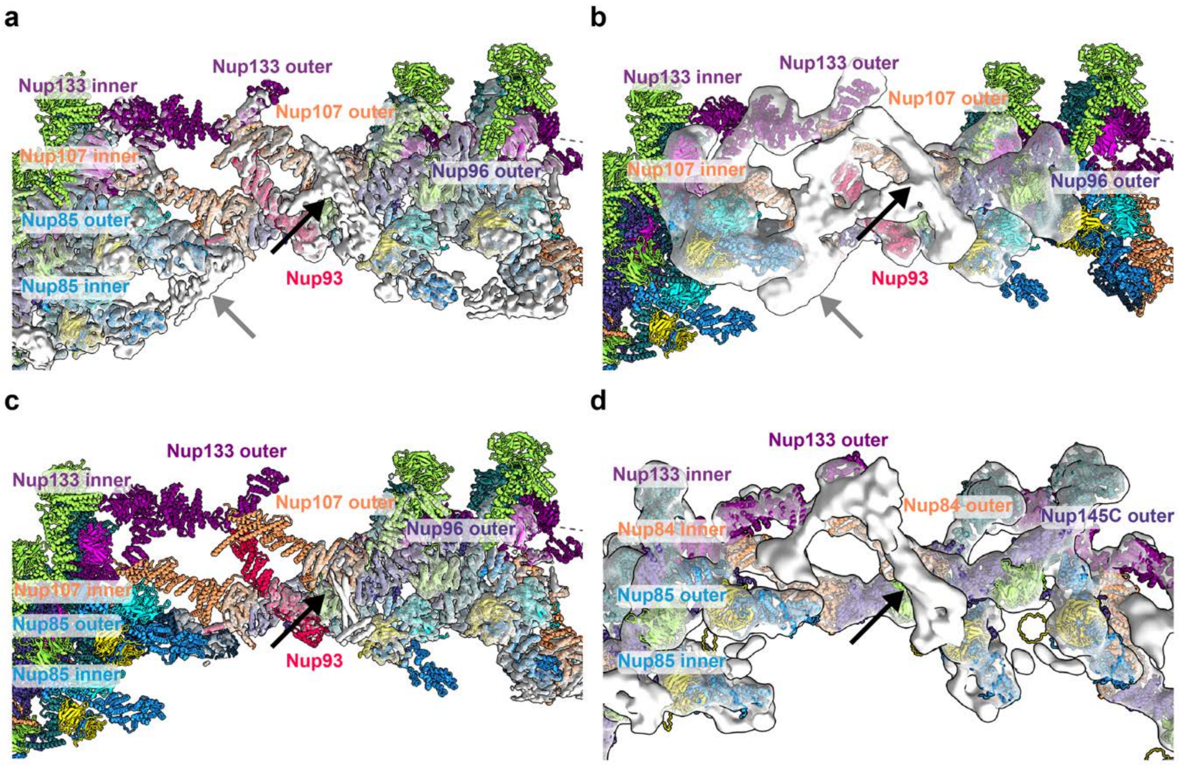

**Fig. S5. GANP and nuclear basket attachment site in the NR in *X. laevis* and yeast. (a)** Isosurface-rendered cryo-EM map of the *X. laevis* NR (EMD-31065) superposed with the respective structural model (PDB: 7wb4 [40]). **(b)** Same as (a) but for (EMD-31941). **(c)** Same as (a) but for (EMD-32394 [40]) **(d)** Cryo-EM map of the yeast NR (EMD-24231 [16]), superposed with structural model of the yeast NR built using AF3 based on the published model[16]. Previously unassigned densities corresponding to the nuclear basket attachment region and GANP domain are indicated with black and gray arrows, respectively. The match of the *X. laevies* Tpr insertion region and Ganp with the apparent unassigned secondary structure in the cryo-EM map of the *X. laevis* NR (EMD-31065) is demonstrated in Figs. S6b and S7b.

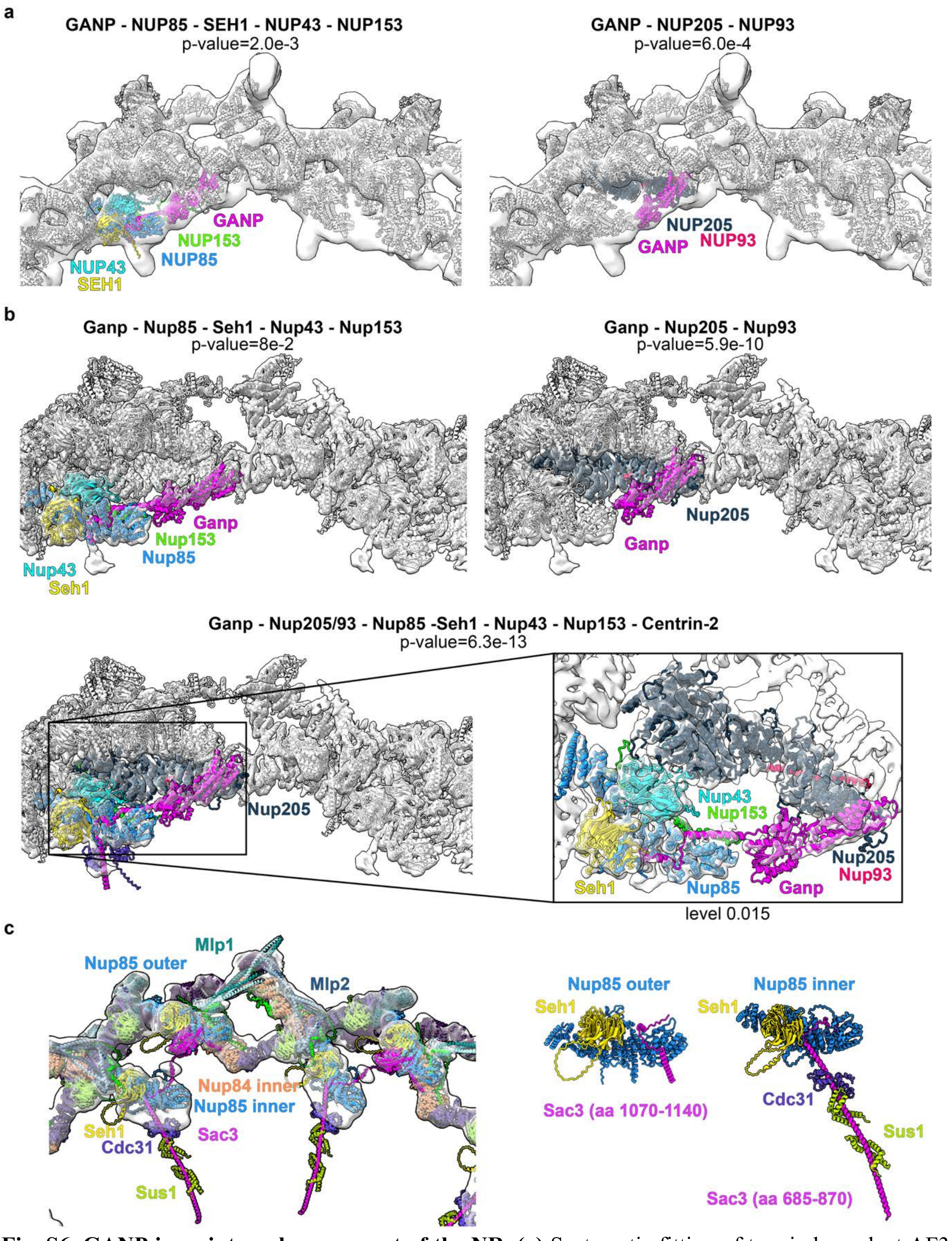

**Fig. S6. GANP is an integral component of the NR.** **(a)** Systematic fitting of two independent AF3 models of the two GANP-containing subcomplexes (shown color-coded) into the NR region of the cryo-ET map of the human macrophage NPC; *p*-values indicated. Top-scoring positions within the NR are shown in context of the cryo-ET map and previously assigned proteins (white) of the NR as seen from nucleoplasm. **(b)** Same as (a) but for three overlapping AF3 models of the *X. laevis* subcomplexes involving Ganp into the cryo-EM map of the NR of the *X. laevis* NPC (EMD-31065). Inset highlights agreement with secondary structure. **(c)** Model of Sac3 (yeast GANP homolog) in the NR of the yeast NPC (*left*) and individual models of the two Sac3-containing subcomplexes (*right*) are shown fitted into the cryo-EM map of the NR of the yeast NPC (EMD-24231 [16]).

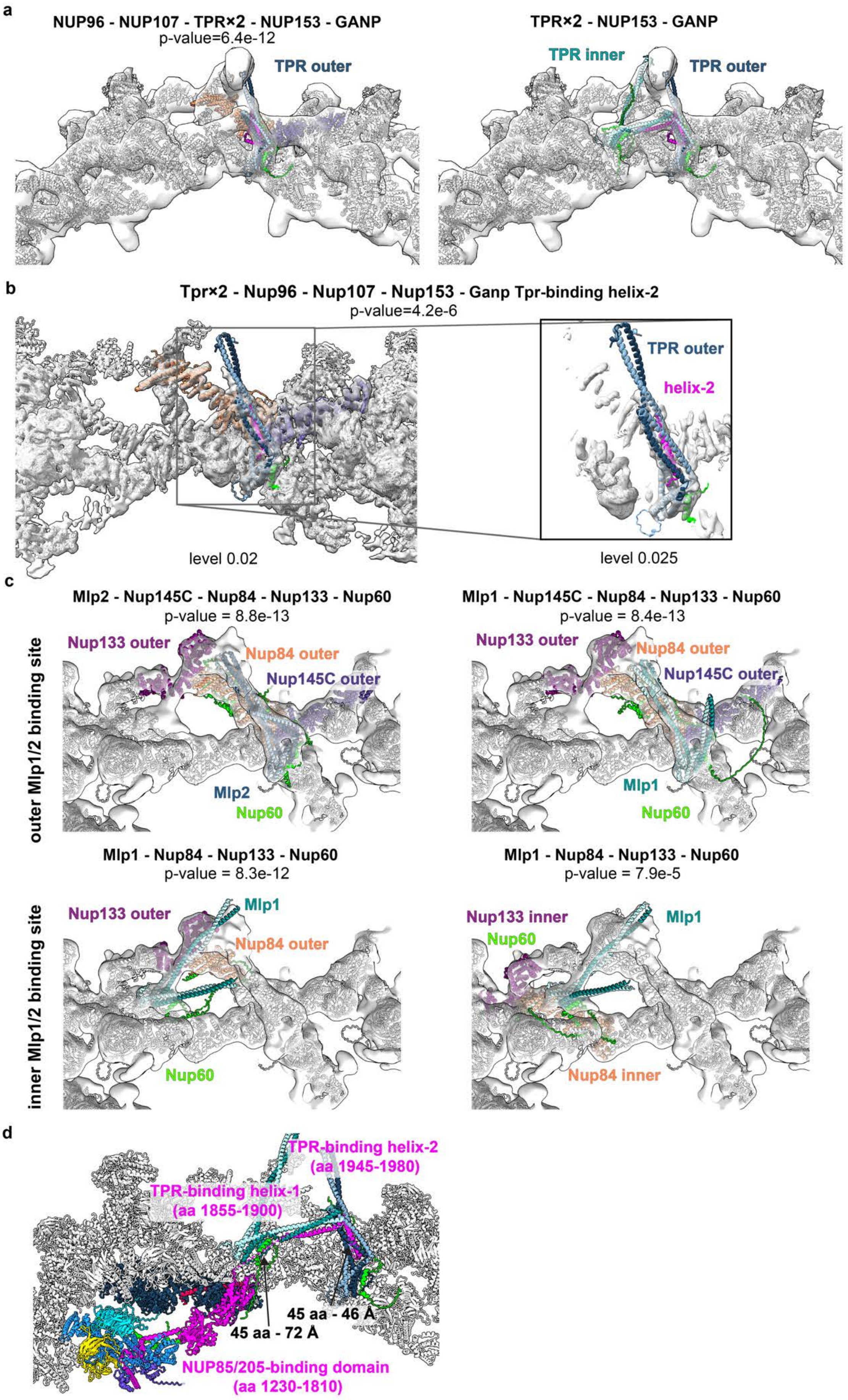

a
NUP96 - NUP107 - TPR×2 - NUP153 - GANP
p-value=6.4e-12
TPR outer
TPR×2 - NUP153 - GANP
TPR inner
TPR outer
b
Tpr×2 - Nup96 - Nup107 - Nup153 - Ganp Tpr-binding helix-2
p-value=4.2e-6
TPR outer
helix-2
level 0.02
level 0.025
c
Mlp2 - Nup145C - Nup84 - Nup133 - Nup60
p-value = 8.8e-13
Nup133 outer
Nup84 outer
Nup145C outer
Mlp2
Nup60
Mlp1 - Nup145C - Nup84 - Nup133 - Nup60
p-value = 8.4e-13
Nup133 outer
Nup84 outer
Nup145C outer
Mlp1
Nup60
outer Mlp1/2 binding site
Mlp1 - Nup84 - Nup133 - Nup60
p-value = 8.3e-12
Nup133 outer
Mlp1
Nup84 outer
Nup60
Mlp1 - Nup84 - Nup133 - Nup60
p-value = 7.9e-5
Nup133 inner
Nup60
Mlp1
Nup84 inner
inner Mlp1/2 binding site
d
TPR-binding helix-2
(aa 1945-1980)
TPR-binding helix-1
(aa 1855-1900)
45 aa - 46 Å
45 aa - 72 Å
NUP85/205-binding domain
(aa 1230-1810)

1372

**Fig. S7. Modeling the TPR attachment region. (a)** Systematic fitting of the AF3 model of the outer TPR insertion region in complex with adjacent Y-complex NUPs, NUP153 and GANP helix-2 (*left*; shown color coded) into the NR region of the cryo-ET map of the human macrophage NPC; single top-scoring position in the NR is shown in context of the cryo-ET map and previously assigned proteins (white) of the NR as seen from nucleoplasm; *p*-value indicated. Manual fit of the inner TPR insertion region in complex with NUP153 and GANP helix-1 (*right*). **(b)** Same as (a; left) but for the respective *X. laevis* AF3 models and the corresponding cryo-EM map (EMD-31065). Inset highlights agreement with secondary structure. **(c)** Same as (a; left) but for the respective AF3 models of the yeast Mlp1/2 insertion region into the corresponding cryo-EM map of the yeast NR (EMD-24231, [16]). AF3 models for the outer insertion region were obtained with both isoforms, Mlp1 and Mlp2 and yielded highly similar results (*top*). Two overlapping AF3 models in which Mlp1 is attached to Y-complex NUPs were obtained for the inner insertion region (*bottom*). The part of the Mlp1 homodimer that projects into the basket filament that is not covered in the cryo-EM map of the NR; *p*-values shown. **(d)** Composite model illustrating how the C-terminal region of GANP is engaged with the NR scaffold at NUP85 and NUP205 and the inner and outer TPR binding site. See also Fig. S2.

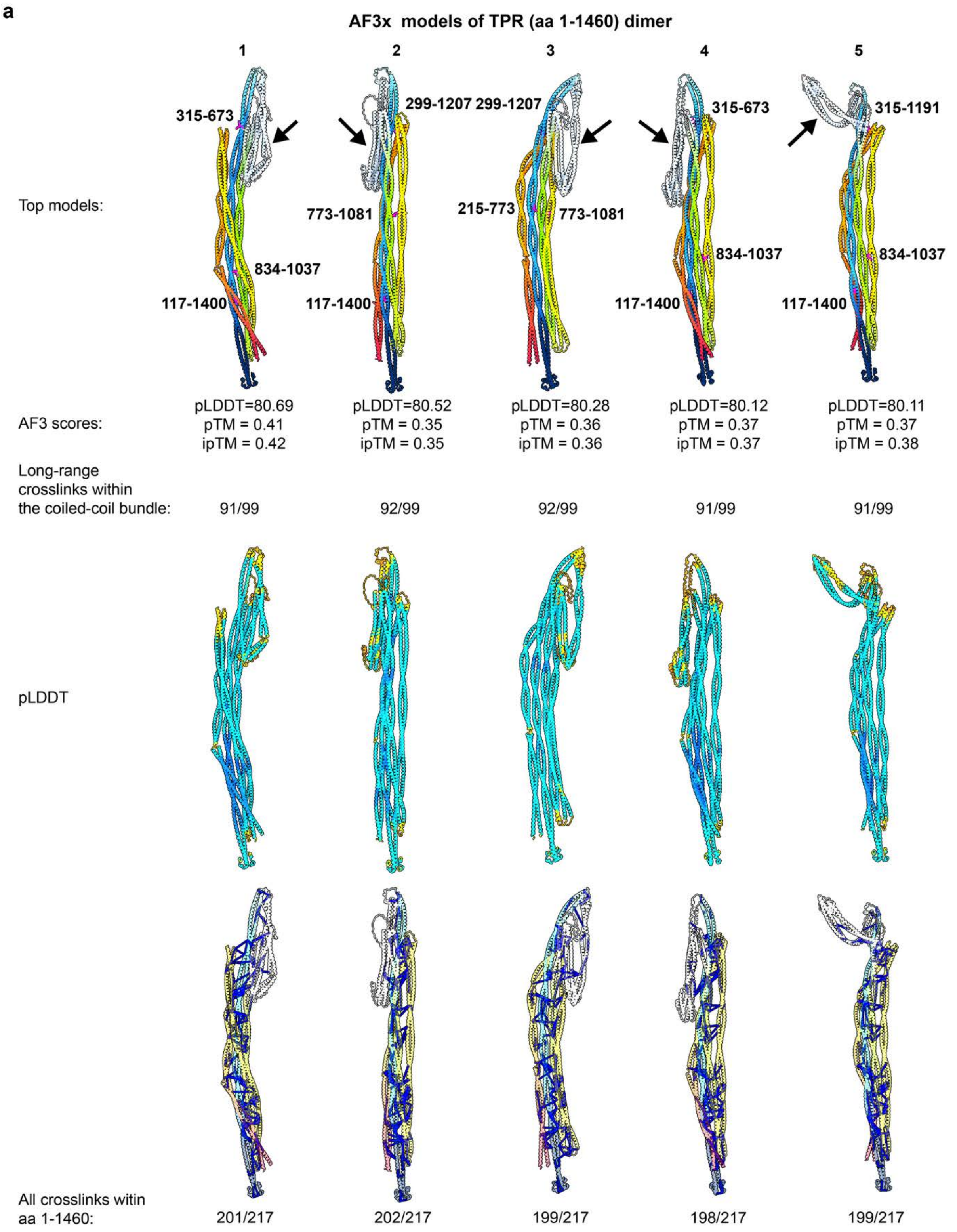
a
AF3x models of TPR (aa 1-1460) dimer
1
2
3
4
5
Top models:
315-673
299-1207
299-1207
315-673
315-1191
773-1081
215-773
773-1081
834-1037
834-1037
834-1037
117-1400
117-1400
117-1400
117-1400
AF3 scores:
pLDDT=80.69
pTM = 0.41
ipTM = 0.42
pLDDT=80.52
pTM = 0.35
ipTM = 0.35
pLDDT=80.28
pTM = 0.36
ipTM = 0.36
pLDDT=80.12
pTM = 0.37
ipTM = 0.37
pLDDT=80.11
pTM = 0.37
ipTM = 0.38
Long-range crosslinks within the coiled-coil bundle:
91/99
92/99
92/99
91/99
91/99
pLDDT
All crosslinks witin aa 1-1460:
201/217
202/217
199/217
198/217
199/217

1388

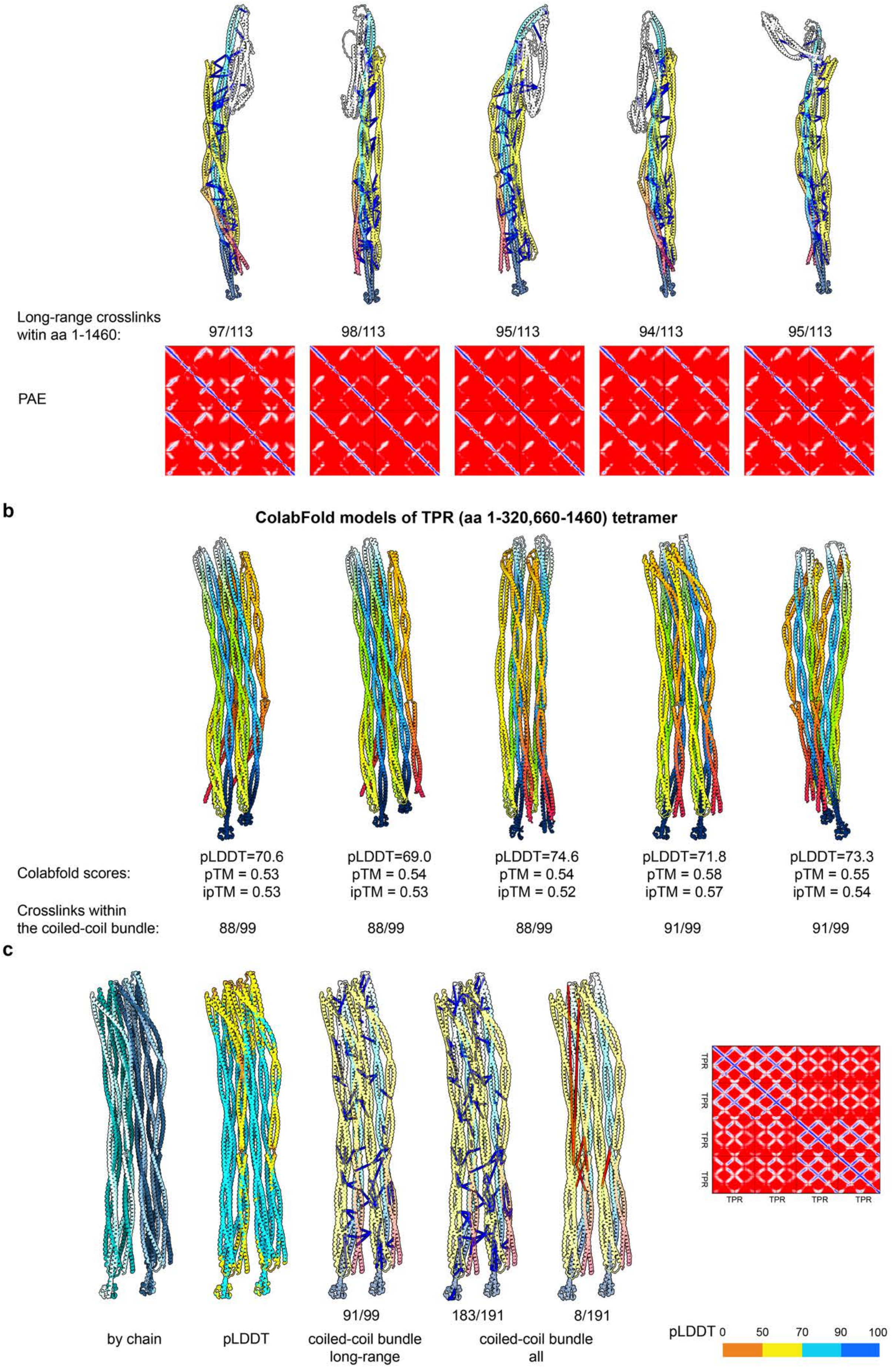
Long-range crosslinks
witin aa 1-1460:
97/113
98/113
95/113
94/113
95/113
PAE
b
ColabFold models of TPR (aa 1-320,660-1460) tetramer
Colabfold scores:
pLDDT=70.6
pTM = 0.53
ipTM = 0.53
pLDDT=69.0
pTM = 0.54
ipTM = 0.53
pLDDT=74.6
pTM = 0.54
ipTM = 0.52
pLDDT=71.8
pTM = 0.58
ipTM = 0.57
pLDDT=73.3
pTM = 0.55
ipTM = 0.54
Crosslinks within
the coiled-coil bundle:
88/99
88/99
88/99
91/99
91/99
c
TPR
TPR
TPR
TPR
TPR
TPR
TPR
TPR
91/99
183/191
8/191
by chain
pLDDT
coiled-coil bundle
long-range
coiled-coil bundle
all
pLDDT 0 50 70 90 100

**Fig. S8. Modeling of the TPR filaments. (a)** From top to bottom: Top five models of the TPR dimer (aa 1-1460) built using AF3x. Models are colored sequentially from N- to C-terminus with explicitly modeled crosslinks shown in magenta and the insertion region indicated by arrows; AF3 scores and the number of satisfied long-range sequence crosslinks (residues separated by ≥20 aa in sequence) within the coiled-coil bundle (aa 1-320 and 660-1460) are indicated; models colored by pLDDT score (legend below); models colored sequentially from N- to C-terminus with satisfied long-range sequence crosslinks mapped are shown with the number of crosslinks indicated below; models with all satisfied crosslinks mapped are shown with the number of crosslinks indicated below (all remaining crosslinks are shown in Fig. S9); PAE heat map shows the expected positional error between residue pairs (blue = high confidence, red = low confidence). **(b)** Example models of the TPR tetramer comprising residues 1–320 and 660–1460, corresponding to the coiled-coil bundle built using ColabFold with AF3x models shown in (a) as templates. **(c)** Top-scoring model of the TPR tetramer. From left to right: model colored by chain, model colored by pLDDT score (legend below); model colored sequentially from N- to C-terminus with satisfied long-range sequence crosslinks mapped; model with all satisfied crosslinks mapped, model with all violated crosslinks mapped; The PAE heat map shows the expected positional error between residue pairs (blue = high confidence, red = low confidence). Mapped crosslinks are shown as pseudo-bonds between Cα atoms of lysines, with ≤40 Å in blue and >40 Å in red. See also Fig. S10b.

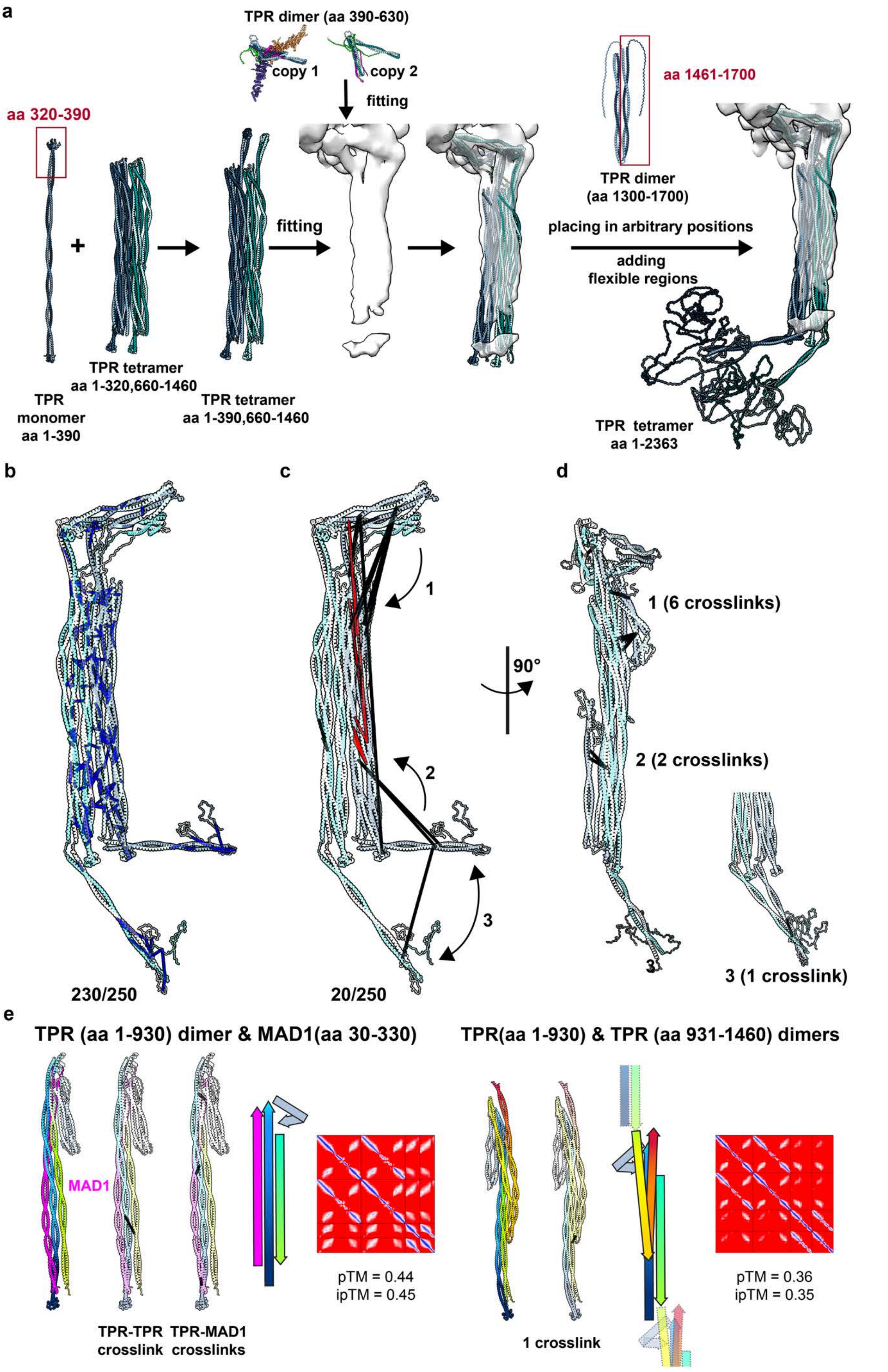
a
TPR dimer (aa 390-630)
copy 1
copy 2
fitting
aa 320-390
aa 1461-1700
TPR dimer
(aa 1300-1700)
fitting
placing in arbitrary positions
adding
flexible regions
TPR tetramer
aa 1-320,660-1460
TPR
monomer
aa 1-390
TPR tetramer
aa 1-390,660-1460
TPR tetramer
aa 1-2363
b
c
d
1
2
3
90°
1 (6 crosslinks)
2 (2 crosslinks)
3 (1 crosslink)
230/250
20/250
e
TPR (aa 1-930) dimer & MAD1(aa 30-330)
TPR(aa 1-930) & TPR (aa 931-1460) dimers
MAD1
pTM = 0.44
ipTM = 0.45
TPR-TPR
crosslink
TPR-MAD1
crosslinks
1 crosslink
pTM = 0.36
ipTM = 0.35

1408

**Fig. S9. Models of the basket filament. (a)** Sequential modeling workflow for the TPR tetramer: TPR fragments fitted into the cryo-ET map of the nuclear basket from human macrophages. The terminal coiled-coils were positioned arbitrarily since there was no interaction with the coiled-coil bundle detected by AF3 (Table S4) and the C-terminal disordered regions were added in arbitrary conformations. Individual components were connected using ChimeraX [105] and ISOLDE [108]**. (b)** All satisfied (≤ 40 Å) crosslinks mapped onto the TPR tetramer model (blue). **(c)** All violated crosslinks >40 Å when mapped onto the TPR tetramer model (red) with crosslinks that can be satisfied in alternative conformations colored in black. **(d)** Crosslinks that can be satisfied due to flexibility of the TPR fragments i.e. the insertion region or the terminal coiled-coil. **(e)** Two additional models obtained using AF3 where TPR interacts with MAD1 (left) or two different TPR dimers interact with alternative manner (*right*). TPR colored sequentially from the N- to the C-terminus, as in Fig. S8; crosslinks that are violated in the main conformation but satisfied in the alternative conformation (for the model with MAD1 only crosslinks involving MAD1 are shown mapped onto the models). Crosslinks are shown as pseudo-bonds between the Cα atoms of lysines; schematic illustrating the topology of the model; PAE heat map indicating the expected positional error between residue pairs (blue = high confidence, red = low confidence).

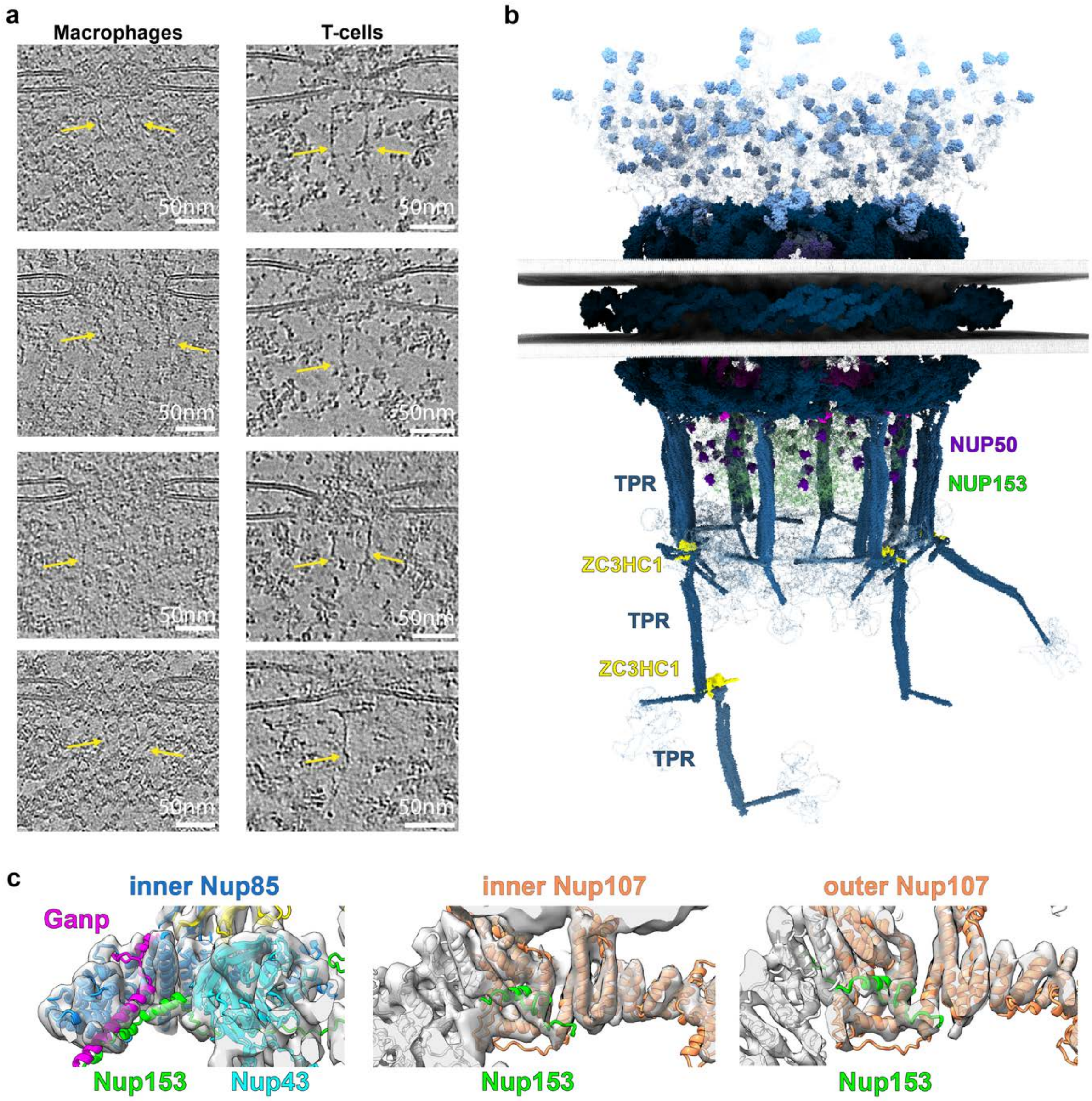

**Fig. S10. Nuclear basket filaments. (a)** Representative slices through tomograms visualizing the extended nuclear filaments (yellow arrows) in Cryo-CARE [112] denoised tomograms from human macrophages and T cells (data from [113]). **(b)** Model of how ZC3HC1 extends nuclear basket filaments by linking two TPR filaments end-to-end. **(c)** AF3 models of *X. laevis* Nup153 in complex with Nup85 subcomplex (*left*) and with Nup107 (*center* and *right*), superposed on the cryo-EM map of the *X. laevis* NR (EMD-31065).

**Table S 1. Cryo-ET data acquisition parameters and STA map information.**

| **Cell type** | Macrophages | | | | | HEK293 |
|---|---|---|---|---|---|---|
| **Microscope** | Titan Krios G4 | | | | | Titan Krios G2 |
| **Voltage (kV)** | 300 | | | | | 300 |
| **Camera** | Falcon 4 | | | | | Gatan BioQuantum-K3 |
| **Magnification** | 53000 | | | | | 33000 |
| **Pixel size (Å/px)** | 2.414 | | | | | 2.682 |
| **Targeted total electron dose ($e^-$/Å$^2$)** | 135 | | | | | 150 |
| **Targeted defocus range (µm)** | -2.0 – 4.0 | | | | | -2.5 – 4.0 |
| **Automation software** | SerialEM | | | | | SerialEM |
| **Tomograms used for. STA/TM** | 149 | | | | | 115 |
| **Initial # of NPC** | 321 | | | | | 247 |
| **Map type** | CR | IR | NR | LR | Basket | Basket |
| **Final # of particles** | 1915 | 1764 | 1596 | 1764 | 1595 | 1925 |
| **Resolution (Å) (FSC 0.143)** | 21.1 | 26.9 | 29.0 | 28.5 | 35.2 | 37.2 |

**Table S 2. Structural similarity between AF3 models.** The table lists the total length of the models, the aligned length used for RMSD and TM-score calculations using USalign[100], the ipTM and pTM values for the first and fifth AF3 models, and the differences in ipTM and pTM between these models.

| MODEL | Total length | Aligned length | RMSD | TM score | ipTM model_0 | ipTM model_4 | pTM model_0 | pTM model_4 | Δ ipTM | Δ pTM |
|---|---|---|---|---|---|---|---|---|---|---|
| NUP93 (aa 170_819) - NUP35×2 | 1302 | 1007 | 3,42 | 0,74 | 0,57 | 0,56 | 0,62 | 0,61 | -0,01 | -0,01 |
| NUP155 - SMPD4 | 2257 | 2191 | 2,98 | 0,94 | 0,68 | 0,68 | 0,66 | 0,66 | 0 | 0 |
| NUP54 (aa 100-507)×2 - NUP58 (aa 245-475)×2 - NUP62 (aa 325-522)×2 - NUP93 (aa 1-90)×2 - NUP98 (aa 450-550) - TMEM209 (aa 320-561) | 2296 | 2216 | 2,89 | 0,93 | 0,75 | 0,75 | 0,76 | 0,76 | 0 | 0 |
| NUP205 -NUP93 (aa 90-150) - TMEM209 (aa 215-300) | 2158 | 2143 | 1,33 | 0,99 | 0,74 | 0,73 | 0,76 | 0,76 | -0,01 | 0 |
| TMEM209 (aa 1-150) - NUP155 | 1541 | 1463 | 4,26 | 0,87 | 0,62 | 0,6 | 0,62 | 0,61 | -0,02 | -0,01 |
| NUP155 - TMEM209 (aa 1-150) - NUP210 (aa 1790-1887) ×2 | 1737 | 1634 | 3,78 | 0,89 | 0,48 | 0,47 | 0,61 | 0,6 | -0,01 | -0,01 |
| NUP93-NUP210 (aa 1850-1887) | 697 | 686 | 2,06 | 0,94 | 0,59 | 0,57 | 0,81 | 0,8 | -0,02 | -0,01 |
| NUP155-POM121 (aa 250-250) | 1492 | 1424 | 2,99 | 0,92 | 0,36 | 0,36 | 0,63 | 0,63 | 0 | 0 |
| NUP188-NUP93 (aa 90-160)-POM121 | 1971 | 1921 | 2,66 | 0,95 | 0,65 | 0,65 | 0,77 | 0,78 | 0 | 0,01 |
| NUP85 - SEH1 - NUP43 - GANP (aa 1100-1810) - NUP153 (aa 200-400)- Centrin-2- Eny2×2 | 2682 | 2441 | 4,38 | 0,86 | 0,67 | 0,66 | 0,71 | 0,71 | -0,01 | 0 |
| NUP205 - NUP93 (aa 90-150) - GANP (aa 1330-1810) | 2554 | 2546 | 2,9 | 0,97 | 0,71 | 0,7 | 0,67 | 0,66 | -0,01 | -0,01 |
| GANP (aa 1050-1250) - ENY2×4 | 605 | 590 | 3,58 | 0,85 | 0,56 | 0,55 | 0,56 | 0,56 | -0,01 | 0 |
| NUP96 (aa 320-937) - NUP107 (aa 140-925) - TPR (aa 390-630)×2 - NUP153 (aa 200-350) -GANP (aa 1900-1980) | 2118 | 2068 | 2,8 | 0,95 | 0,65 | 0,65 | 0,69 | 0,69 | 0 | 0 |
| TPR (aa 390- 630) - NUP153 (aa 250-400) - GANP (aa 1810-1910) | 734 | 619 | 3,47 | 0,77 | 0,52 | 0,49 | 0,51 | 0,49 | -0,03 | -0,02 |
| NUP85 - SEH1 - NUP43 - NUP153 (aa 150-350) | 1597 | 1537 | 2,2 | 0,94 | 0,73 | 0,73 | 0,76 | 0,76 | 0 | 0 |
| NUP107- NUP153 (aa 150-350) | 1007 | 891 | 2,52 | 0,86 | 0,64 | 0,64 | 0,66 | 0,67 | 0 | 0,01 |
| NUP93 (aa 170-819) - NUP153 (aa 250-450) | 851 | 757 | 3,1 | 0,84 | 0,68 | 0,67 | 0,69 | 0,69 | -0,01 | 0 |
| NUP50 (aa 120-210) - NUP153 (aa 500-600) | 2097 | 1990 | 4,5 | 0,89 | 0,6 | 0,6 | 0,65 | 0,65 | 0 | 0 |
| TPR (aa 1-200)-TPR (aa 390-630)-ZC3HC1 | 1384 | 1217 | 5,17 | 0,77 | 0,46 | 0,46 | 0,52 | 0,52 | 0 | 0 |
| GANP (aa 1-1000) - PCID2 - DSS1 - DDX39B | 1897 | 1492 | 3,15 | 0,76 | 0,85 | 0,84 | 0,67 | 0,67 | -0,01 | 0 |
| NUP214 (aa 1-430) - DDX19B - GLE1 (aa 380-698) - NUP42 (aa 380-423) | 1272 | 1248 | 1,37 | 0,97 | 0,8 | 0,78 | 0,78 | 0,76 | -0,02 | -0,02 |

**Table S 3. Modeling of NUP155 interactions using AF3.** The table lists proteins tested for interaction with NUP155, including the three interacting and various non-interacting models. For each protein, ipTM and pTM values are reported, along with the corresponding PAE heat map, which indicates the expected positional error between residue pairs (blue = high confidence, red = low confidence).

| Protein | iPTM | PTM | PAE |
|---|---|---|---|
| Positive | | | |
| NDC1-ALADIN | 0.5 | 0.56 | 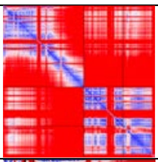 |
| TMEM209 (aa 1-200)-NUP210 (aa 1850-1887) | 0.48 | 0.61 | 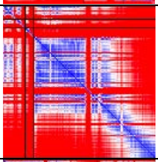 |
| SMPD4 | 0.68 | 0.66 | 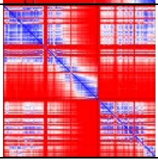 |
| Negative | | | |
| Centrin-2 | 0.3 | 0.59 | 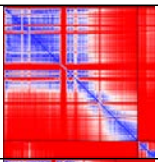 |
| DDX19B | 0.18 | 0.53 | 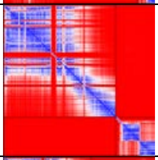 |
| ELYS aa 1–1000 | 0.23 | 0.47 | 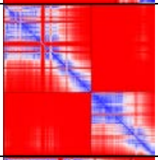 |
| EMERIN | 0.22 | 0.58 | 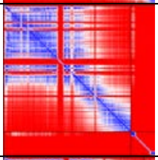 |
| ENY2 | 0.23 | 0.61 | 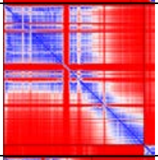 |
| GANP aa 1000–1980 | 0.24 | 0.47 | 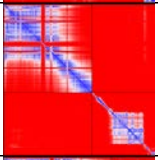 |
| GANP aa 1–550 | 0.21 | 0.55 | 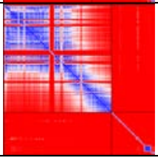 |
| GANP aa 550–1000 | 0.18 | 0.53 | 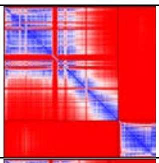 |
| LBR | 0.21 | 0.5 | 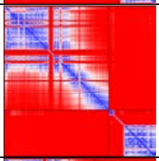 |
| NUP107 | 0.2 | 0.45 | 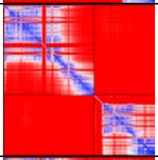 |
| NUP133 | 0.24 | 0.47 | 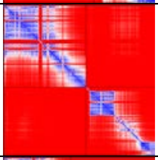 |
| NUP188 | 0.24 | 0.49 | 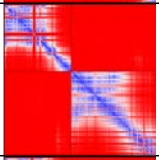 |
| NUP205 | 0.28 | 0.49 | 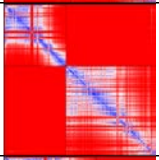 |
| NUP214 aa 1–900 | 0.2 | 0.47 | 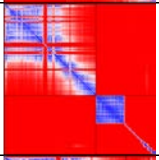 |
| NUP35 | 0.38 | 0.57 | 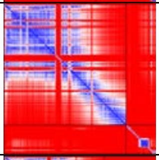 |
| NUP358 aa 1–700 | 0.18 | 0.48 | 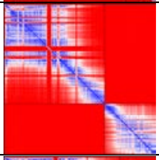 |
| NUP42 | 0.19 | 0.55 | 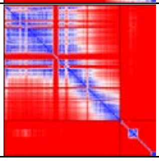 |

| | | | |
|---|---|---|---|
| NUP43 | 0.24 | 0.52 | 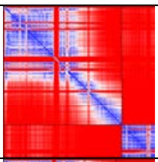 |
| NUP50 | 0.17 | 0.53 | 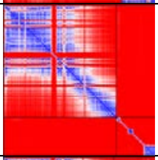 |
| NUP54 | 0.19 | 0.52 | 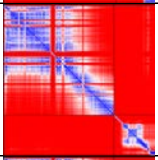 |
| NUP58 | 0.16 | 0.51 | 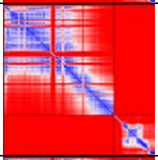 |
| NUP62 | 0.22 | 0.53 | 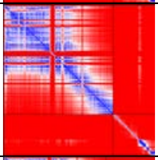 |
| NUP85 | 0.19 | 0.48 | 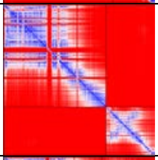 |
| NUP88 | 0.24 | 0.51 | 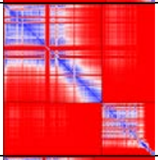 |
| NUP96 | 0.18 | 0.47 | 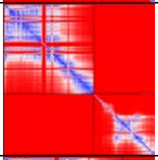 |
| RAE1 | 0.2 | 0.53 | 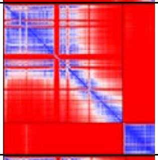 |
| SEC13 | 0.25 | 0.54 | 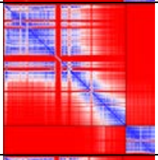 |
| SEH1 | 0.16 | 0.52 | 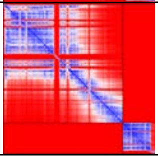 |

| | | | |
|---|---|---|---|
| SUN1 | 0.19 | 0.49 | 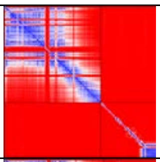 |
| SUN2 | 0.19 | 0.5 | 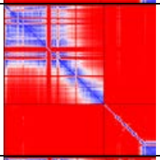 |
| TMEM201 | 0.31 | 0.53 | 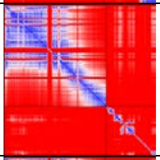 |
| TMEM214 | 0.22 | 0.5 | 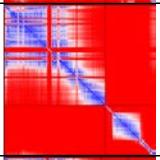 |
| TPR aa 1–390 | 0.22 | 0.48 | 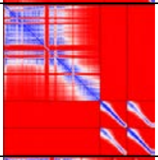 |
| TPR aa 390–660 | 0.23 | 0.52 | 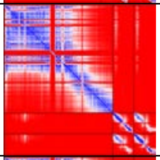 |
| TPR aa 660–930 | 0.22 | 0.52 | 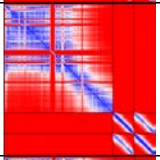 |
| TPR aa 930–1180 | 0.22 | 0.52 | 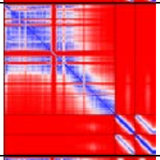 |
| TPR aa 1180–1460 | 0.32 | 0.58 | 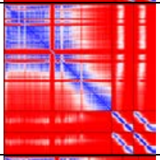 |
| TPR aa 1460–1650 | 0.21 | 0.53 | 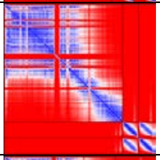 |
| ZC3HC1 | 0.2 | 0.53 | 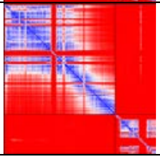 |

**Table S 4. AF3 models of interactions between the C-terminal coiled-coil of TPR (aa 1460-1650) and the remaining region of TPR.** ipTM and pTM values are reported for each model, along with the corresponding PAE heat map, which indicates the expected positional error between residue pairs (blue = high confidence, red = low confidence).

| Protein 1 | Protein 2 | iPTM | PTM | PAE |
|---|---|---|---|---|
| TPR aa 1–390 | TPR aa 1460–1700 | 0.25 | 0.3 | 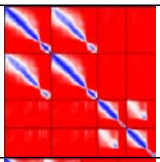 |
| TPR aa 700–1500 | TPR aa 1460–1700 | 0.2 | 0.27 | 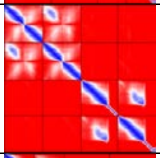 |
| TPR aa 1300–1459 | TPR aa 1460–1700 | 0.24 | 0.3 | 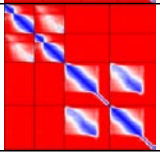 |

**Table S 5. Composition and molecular weight of the NPC.** List of NUPs and associated proteins with their copy numbers per NPC, calculated molecular weight per molecule, and total molecular weight contribution. Proteins highlighted in red indicate NUPs that were either newly identified in this study or for which we used a different copy number than in the previous calculation [16]. The final row shows the total estimated molecular weight of the NPC.

| Protein | Copies | Molecular Weight (Da, each) | Total Molecular Weight (Da) |
|---|---|---|---|
| NUP160 | 32 | 162119.71 | 5187830.56 |
| NUP96 | 32 | 105911.68 | 3389173.76 |
| NUP85 | 32 | 75018,53 | 2400593.12 |
| SEH1 | 32 | 39648.25 | 1268744.16 |
| SEC13 | 32 | 35540.30 | 1137289.76 |
| NUP107 | 32 | 106373.15 | 3403940.80 |
| NUP133 | 32 | 128977.61 | 4127283.36 |
| NUP358 | 40 | 358196.40 | 14327855.80 |
| NUP43 | 32 | 42150.50 | 1348816.16 |
| ELYS | 16 | 252496.05 | 4039936.72 |
| NUP37 | 32 | 36707.38 | 1174636.32 |
| NUP188 | 16 | 196041.00 | 3136656.00 |
| NUP205 | 40 | 227919.17 | 9116766.60 |
| NUP155 | 48 | 155197.65 | 7449487.20 |
| NUP93 | 56 | 93487.38 | 5235293.28 |
| NUP35 | 32 | 34773.45 | 1112750.56 |
| NUP62 | 48 | 53254.40 | 2556211.44 |
| NUP54 | 32 | 55435.21 | 1773926.72 |
| NUP58 | 32 | 60896.45 | 1948686.40 |
| NUP88 | 16 | 83541.08 | 1336657.28 |
| NUP214 | 16 | 213617.56 | 3417881.04 |
| NUP98 | 48 | 91684.07 | 4400835.6 |
| NDC1 | 16 | 76304.03 | 1220864.56 |
| NUP210 | 64 | 205109.48 | 13127006.72 |
| ALADIN | 16 | 59573.58 | 953177.36 |
| POM121 | 32 | 127718.71 | 4086998.72 |
| TPR | 32 | 267291.08 | 8553314.56 |
| NUP153 | 24 | 153936.97 | 3694487.28 |
| NUP50 | 48 | 50143.79 | 2406901.92 |
| NUP42 | 16 | 44871.22 | 717939.52 |
| DDX19B | 16 | 53926.43 | 862822.88 |
| GLE1 | 16 | 79835.59 | 1277369.52 |
| SMPD4 | 16 | 97808.95 | 1564943.28 |
| TMEM209 | 16 | 62921.13 | 1006738.08 |
| GANP | 8 | 218402.60 | 1747220.76 |
| CENTRIN-2 | 8 | 19738.28 | 157906.28 |
| ENY2 | 32 | 11528.52 | 368912.64 |
| | | **Total:** | **125037857.04** |
| ZC3HC1 | 8 | 55260.85 | 442086.84 |
| DSS1 | 8 | 8277.59 | 66220.68 |
| PCID2 | 8 | 46029.55 | 368236.40 |
| DDX39B | 8 | 48990.94 | 391927.56 |
| | | **Total:** | **126306328.52** |

**Table S6. Confidence assessment of individual assignments.** Individual analyses, experimental data and literature that support each individual assignment are listed by protein domain.

| NUP | AF3 models | Crosslinks | EM-Map | Experimental validation | Overall Confidence |
|---|---|---|---|---|---|
| | | **Inner ring** | | | |
| **TMEM209 NTD**<br>(aa 1–120)<br>interacting with<br>**NUP155-NUP210** | **Human**<br>High confidence<br>TMEM209–NUP155–NUP210×2<br>TMEM209–NUP155<br>(Fig. S2a)<br>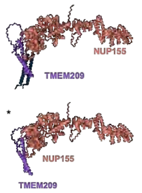 | | **Human**<br>Superposition with the established outer copy of NUP155 in the IR positions the transmembrane helices of TMEM209 and NUP210 into the membrane, NUP210 transmembrane helices in the correct direction and proximate to the NUP210 LR domains. | Proximity labeling MS (Fig. S4b).<br><br>Literature: NE/NPC co-localization of SMPD4, NUP210 co-IP [55]. | high |
| **TMEM209**<br>(aa 215–300)<br>interacting with<br>**NUP205-NUP93** | **Human**<br>High confidence<br>TMEM209–NUP205–NUP93<br>(Fig. S2a)<br>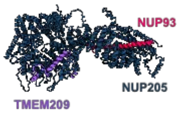 | **Human**<br>Eight satisfied crosslinks between TMEM209 and NUP205/NUP155:<br>TMEM209 252–NUP205 975<br>TMEM209 252–NUP205 1230<br>TMEM209 230–NUP205 974<br>TMEM209 232–NUP205 1271<br>TMEM209 304–NUP205 571<br>TMEM209 287–NUP155 740<br>TMEM209 288–740 NUP155<br>TMEM209 288–NUP155 766<br>(Fig. 2d)<br>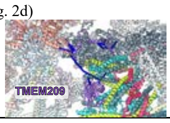 | **Human**<br>Superposition with the established position of NUP205 in the IR positions TMEM209 aa 215-300 such that it can be connected to the TMEM209 CTD (aa 325-561). | | excellent |
| **TMEM209 CTD**<br>(aa 325–561)<br>interacting with two<br>**NUP98–93–62–58–54**<br>subcomplexes | **Human**<br>High confidence<br>TMEM209 CTD–2× (NUP98–93–62–58–54)<br>(Fig. S2a)<br>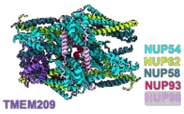 | **Human**<br>Three satisfied crosslinks between TMEM209 and NUP58/NUP205:<br>TMEM209 304–NUP58 320,<br>TMEM209 519–NUP58 330,<br>TMEM209 415–NUP205 892<br>(Fig. 2d)<br>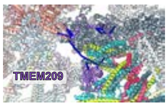 | **Human**<br>Systematic fitting into the IR region of the cryo-ET map of the human NPC (EMD-14322) (Fig. S3a) identifies two highly significant C2-symmetric top-scoring positions, consistent with the established positions of the two NUP62-subcomplexes.<br>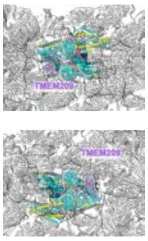 | | excellent |
| **SMPD4**<br>interacting with<br>**NUP155** | **Human**<br>High confidence<br>NUP155–SMPD4<br>(Fig. S2a)<br>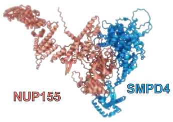 | **Human**<br>Satisfied crosslink:<br>SMPD4 277–NUP155 1069<br><br>Slightly violated crosslink between SMPD4 and NUP35; possibly explained by conformational changes or an additional copy of NUP35:<br>SMPD4 604–NUP35 218<br>(Fig. S3c)<br>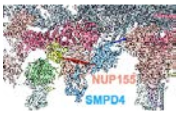 | **Human**<br>Systematic fitting into the IR region of the cryo-ET map of the human NPC (EMD-14322) (Fig. S3b) identifies two highly significant C2-symmetric top-scoring positions, consistent with the established positions of the two inner copies of NUP155 within the IR.<br>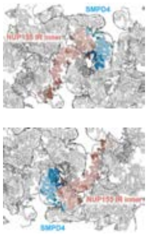<br>In the model, the amphipathic and transmembrane helices of the SMPD4 CTD are positioned within the membrane. | Immunofluorescence staining of SMPD4 in HEK293 cells (Fig. S4a).<br><br>Proximity labeling MS (Fig. S4b).<br><br>Literature: proximity labeling [46], co-IP [51]. | high |

| | | | | | |
|---|---|---|---|---|---|
| | | | 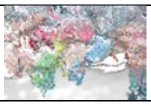 | | |
| **NUP35 folded domain**<br>(aa 150–250)<br>interacting with<br>**NUP93** | **Human**<br>High confidence<br>NUP93–NUP35×2<br>(Fig. S2a)<br>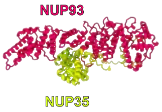 | **Human**<br>Four satisfied crosslinks supporting the direct interaction of NUP93 and NUP35 as in the model:<br>NUP35 227–NUP93 600<br>NUP35 218–NUP93 370<br>NUP35 227–NUP93 370<br>NUP35 227–NUP35 294<br><br>Three slightly violated crosslinks between NUP35 and proximate proteins:<br>NUP35 218–SMPD4 604<br>NUP35 218–NUP205 496<br>NUP35 227–NUP205 1988<br><br>This may be explained by alternative conformations or additional copies of NUP35<br>(Fig. S3c)<br>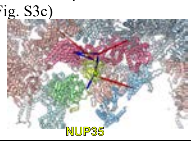 | **Human**<br>Systematic fitting into the IR region of the cryo-ET map of the human NPC (EMD-14322) identifies two highly significant, C2-symmetric top-scoring positions, consistent with the established positions of the outer NUP93 copies in the IR<br>(Fig. S3d).<br><br>IR cytoplasmic side:<br>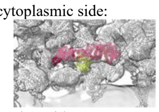<br>IR nuclear side:<br>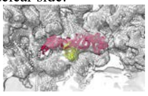 | Literature: Crosslinking MS and integrative modeling of the yeast NPC [16,17]. | high |
| **NUP210**<br>(aa 1870–1887)<br>interacting with<br>**NUP93** | **Human**<br>High confidence<br>NUP93–NUP210<br>(Fig. S2a)<br>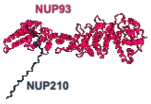 | **Human**<br>Satisfied crosslink:<br>NUP210 1872–NUP93 718<br>(Fig. S3e)<br>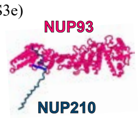 | **Human**<br>The linker between the NUP210 transmembrane helix and its NUP93-interacting region is long enough to allow an interaction with the two inner copies of NUP93 of the IR, but not with the outer copies. | | good |
| **POM121**<br>(aa 305–320)<br>interacting with<br>**NUP155** | **Human**<br>Low confidence<br>NUP155–POM121<br>(Fig. S2a)<br>*<br>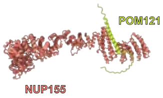 | **Human**<br>Satisfied crosslink:<br>POM121 339–NUP155 1069<br>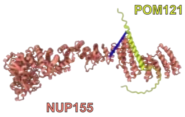 | | Literature: The crosslinks and AF3 model would support a role of POM121 in linking the NUP155 connector towards the Y-complex, in agreement with a previous proposal [84]. | moderate |
| **POM121**<br>(aa 500–555)<br>interacting with<br>**NUP188** | **Human**<br>High confidence<br>NUP188–NUP93–POM121<br>(Fig. S2a)<br>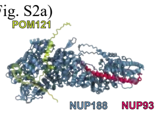 | **Human**<br>Two satisfied crosslinks between POM121 and NUP188:<br>POM121 489–NUP188 463<br>POM121 489–NUP188 167<br>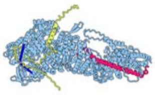 | | Literature: The crosslinks and AF3 model would support a role of POM121 in linking the outer IR core module towards the Y-complex, in agreement with previous a proposal [84]. | good |
| | | **Nuclear Ring** | | | |
| **GANP**<br>(aa 1230–1810)<br>interacting with the<br>**inner NUP85** and **NUP205** subcomplexes in the NR | **Human**<br>High confidence<br>NUP85–SEH1–NUP43–GANP–NUP153<br>(Fig. S2b)<br>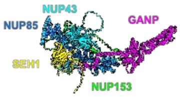<br>High confidence<br>NUP205–NUP93–GANP<br>(Fig. S2b)<br>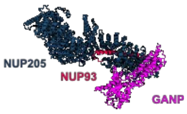 | **Human**<br>Two satisfied crosslinks between GANP and NUP153 bound to the inner NUP107/TPR:<br>GANP 1801–NUP153 353<br>GANP 1801–NUP153 356<br>(Fig. 4f)<br>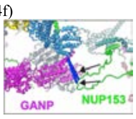<br>Thirteen satisfied crosslinks between NUP153 and NUP85 or SEH1:<br>NUP153 317–NUP85 92<br>NUP153 326–NUP85 92<br>NUP153 326–NUP85 218<br><br>NUP153 356–NUP85 218<br>NUP153 353–NUP85 218<br>NUP153 353–NUP85 103<br>NUP153 353–SEH1 251<br>NUP153 353–NUP85 92<br>NUP153 356–NUP85 92<br><br>NUP153 371–NUP85 218<br>NUP153 371–NUP85 95<br>NUP153 371–NUP85 92<br>NUP153 371–NUP85 103 | **Human**<br>Systematic fitting of the AF3 models of the NUP85–SEH1–NUP43–GANP–NUP153 and NUP205–NUP93–GANP subcomplexes into the NR region of the cryo-ET map of the human macrophage NPC identifies a single top-scoring position for each model, independently placing GANP in the same previously unassigned density and aligning with the known positions of the well-characterized scaffold NUPs, NUP85, SEH1, NUP93 and NUP205, in the NR<br>(Fig. S6a).<br>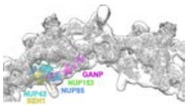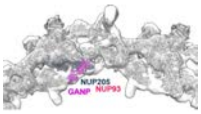 | Literature: NPC localization in human cells [67] and yeast [114,115]; localization to the nuclear basket [10]; TPR is required for GANP localization to the NPC [11]. | excellent |

| | | | | | |
|---|---|---|---|---|---|
| | *X. laevis*<br>High confidence<br>Nup85–Seh1–Nup43–Nup153–Ganp<br>(Fig. S2i)<br>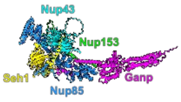<br>High confidence<br>Ganp–Nup205–Nup93<br>(Fig. S2k)<br>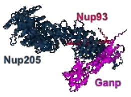<br><br>**Yeast**<br>High confidence<br>Nup85– Seh1– Sac3–Cdc31– Sus1×2<br>(Fig. S2k)<br>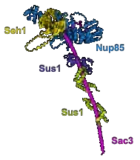 | Two satisfied crosslinks between NUP153 and NUP160 or NUP205:<br>NUP153 353–NUP205 892<br>NUP153 371–NUP160 1331<br><br>Three slightly violated NUP153-SEH1 crosslinks (likely due to flexibility of NUP153):<br>NUP153 317–SEH1 251<br>NUP153 317–SEH1 255<br>NUP153 326–SEH1 251<br>(Fig.5f)<br>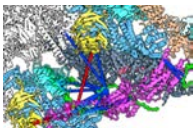 | *X. laevis*<br>Identical results are obtained by systematic fitting of the respective AF3 models of the *X. laevis* subcomplexes into the cryo-EM map of the NR of the *X. laevis* NPC (EMD-31065)<br>(Fig. S6b); secondary structure of Ganp observed in cryo-EM map is consistent with the Ganp fold.<br>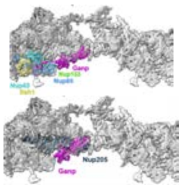<br>Nup153 helix is resolved at the interface with the inner Nup85 subcomplex in the density of the cryo-EM map of the NR of the *X. laevis* NPC (EMD-31065).<br>(Fig. S10c)<br>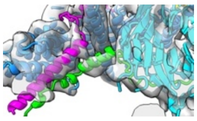<br>**Yeast**<br>The absence of density corresponding to the human NUP85/205 domain in the yeast NR cryo-EM map is consistent with the absence of yNup192 (hNUP205) in the yeast NR and the lack of a yNup192 (hNUP205)-binding domain in yeast Sac3.<br>(Fig. S6c)<br>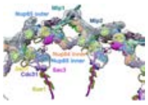 | | |
| **ENY2**<br>four copies interacting with **GANP** | **Human**<br>High confidence<br>GANP–ENY2×4<br>(Fig. S2b)<br>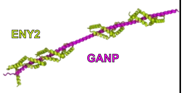<br>*X. laevis*<br>High confidence<br>Ganp–Eny2×4<br>(Fig. S2i) | **Human**<br>Four satisfied GANP–ENY2 crosslinks supporting three copies of ENY2:<br>GANP 1194–ENY2 19<br>GANP 1194–ENY2 43<br>GANP 1140–ENY2 43<br>GANP 1140–ENY2 51<br>(Fig. 4f)<br>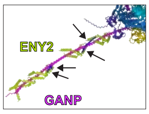 | Not covered by density in any of the cryo-EM maps analyzed here, therefore likely flexible. | **Human**<br>X-ray structure of the human GANP–ENY2×2 (PDB-4DHX[12]).<br><br>**Yeast**<br>X-ray structure of the Sac3–Cdc31–Sus1×2 (PDB–3FWC[114]). | high |

<table>
<tr>
<td></td>
<td>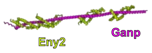

Yeast – 2 copies<br>High confidence<br>Nup85– Seh1– Sac3–<br>Cdc31– Sus1×2<br>(Fig. S2k)
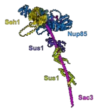
</td>
<td></td>
<td></td>
<td></td>
<td></td>
</tr>
<tr>
<td>Centrin-2<br>interacting with GANP</td>
<td>Human<br>High confidence<br>NUP85–SEH1–NUP43–<br>GANP–Centrin-2–ENY2×2<br>(Fig. S2b)
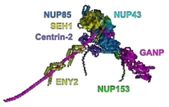

X. laevis<br>High confidence<br>Nup85–Seh1–Nup43–<br>Ganp-Centrin-2–Eny2×2<br>(Fig. S2i)
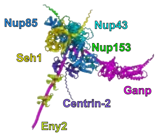

Yeast<br>Nup85–Seh1– Sac3–<br>Cdc31–Sus1×2 (Fig. S2k)
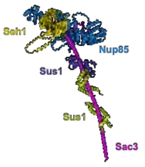
</td>
<td></td>
<td></td>
<td>Yeast<br>X-ray structure of the Sac3–Cdc31–Sus1×2 (PDB–3FWC[114]).</td>
<td>high</td>
</tr>
<tr>
<td>Outer TPR insertion region<br>(aa 390–630)<br>interacting with<br>GANP TPR binding helix-2 (aa 1945–1980), NUP153 (aa 305–320) and NUP107/NUP96</td>
<td>Human<br>High confidence<br>NUP96–NUP107–TPR×2–<br>NUP153–GANP<br>(Fig. S2c)
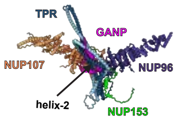

X. laevis<br>High confidence model<br>Nup96–Nup107– Tpr×2–<br>Nup153–Ganp<br>(Fig. S2l)
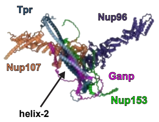
</td>
<td>Human<br>Satisfied TPR–NUP96 crosslink:<br>TPR 474–NUP96 525<br>Satisfied TPR–GANP crosslink:<br>GANP 1951–TPR 474<br>(Fig. 4f)
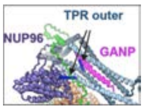

Six satisfied crosslinks between NUP153 and TPR:<br>NUP153 353–TPR 474<br>NUP153 353–TPR 477<br>NUP153 353–TPR 494<br>NUP153 356–TPR 474<br>NUP153 356–TPR 477<br>NUP153 356–TPR 494<br><br>Two satisfied crosslinks between NUP153 and NUP96:<br>NUP153 371–NUP96 242<br>NUP153 384–NUP96 327<br><br>(Fig. 5f)
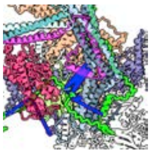</td>
<td>Human<br>Systematic fitting into the NR region of the cryo-ET map of the human macrophage NPC identified a single top-scoring position where TPR occupies the previously unassigned density, while NUP107 and NUP96 overlap with their known positions in the NR outer Y-complex.<br>(Fig. S7a)
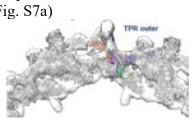

X. laevis<br>Identical results are obtained by systematic fitting of the corresponding AF3 model of the X. laevis subcomplex into the cryo-EM map of the NR of the X. laevis NPC (EMD-31065)<br>(Fig. S7b)
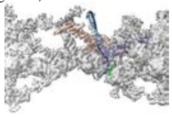</td>
<td>Literature: TPR region aa 437-513 is required (but not sufficient) for TPR localization to the NPC basket, shown in mutagenesis studies [30].<br><br>Literature: Y2H, GST-PD showing that the TPR NBD (NPC binding domain) binds Nup153 [29].<br><br>Literature: Direct interaction of NUP153 mediates TPR binding to the NR[29].</td>
<td>excellent</td>
</tr>
</table>

| | | | | | |
|---|---|---|---|---|---|
| | **Yeast**<br>High confidence model Mlp2×2–Nup84–Nup145C–Nup133–Nup60<br>(Fig. S2l)<br>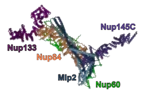 | | Ganp Tpr binding helix-2 and Nup153 helix (aa 305-320) are resolved in the density of the cryo-EM map of the NR of the *X. laevis* NPC (EMD-31065)<br>(Fig. S7b)<br>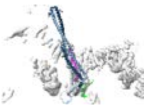<br>**Yeast**<br>Systematic fitting of the corresponding AF3 yeast model into the cryo-EM map of the NR of the yeast NPC (EMD-24231) places Mlp2 dimer at the position corresponding to the outer TPR copy in the human NR.<br>(Fig. S7c)<br>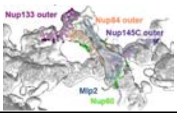 | | |
| **Inner TPR insertion region**<br>(aa 390–630)<br>interacting with<br>**GANP TPR binding helix-1** (aa 1855–1900) and **NUP153** (aa 305–320) | **Human**<br>Moderate confidence TPR×2–NUP153–GANP<br>(Fig. S2d)<br>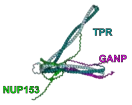<br>***X. laevis***<br>Moderate confidence Tpr×2–Ganp-Nup153<br>(Fig. S2j)<br>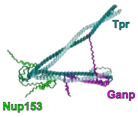<br>**Yeast**<br>Two independent, high confidence models Mlp1×2–Nup84–Nup133–Nup60<br>(Fig. S2l)<br>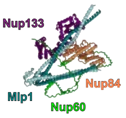<br>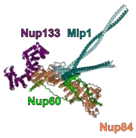 | **Human**<br>Satisfied TPR–NUP93 crosslink:<br>TPR 477–NUP 93 718<br>(Fig. 4f)<br>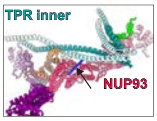<br>Two satisfied crosslinks between GANP and NUP153 bound to the inner NUP107/TPR:<br>GANP 1801–NUP 153 353<br>GANP 1801–NUP 153 356<br>(Fig. 4f)<br>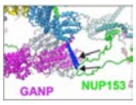<br>*Six satisfied crosslinks between NUP153 and TPR:<br>NUP153 353–TPR 474<br>NUP153 353–TPR 477<br>NUP153 353–TPR 494<br>NUP153 356–TPR 474<br>NUP153 356–TPR 477<br>NUP153 356–TPR 494<br><br>*These crosslinks are also satisfied by the NUP153 interacting with the outer copies of NUP107/TPR. | **Human**<br>A second TPR attachment site is supported by the cryo-ET map, where the TPR filament branches proximate the NR scaffold. (Fig. 1a; Fig. 5c)<br><br>It is not identified by systematic fitting, possibly due to its size, but explains the experimentally observed density in the respective region.<br>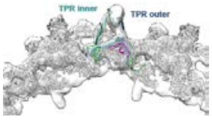<br>***X. laevis***<br>Density not well resolved in this region.<br><br>**Yeast**<br>Systematic fitting of the two AF3 models into the cryo-EM map of the NR of the yeast NPC (EMD-24231) consistently assigns the position of Mlp1 dimer.<br>(Fig. S7c)<br>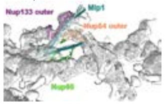<br>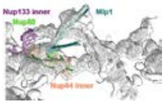 | Literature: A second attachment site is supported by stoichiometric measurements of NPC components in *S. pombe* and human tissue culture cells [61,116].<br><br>Literature: Deletion of NUP153 and TPR [10] or TPR [11] compromises association of TREX-2 (GANP) with the nuclear basket.<br><br>Literature: PD of Sac3 identifies Mlp1/1 and Nup1 as interactors [117].<br><br>Literature: Direct interaction of NUP153 mediates TPR binding to the NR[29]. | good |
| **TPR**<br>coiled-coil bundle<br>(aa 1–390 and 660–1460) | **Human**<br>Moderate confidence of AF3 models probably due to underrepresentation of elongated coiled-coils in the AF3 training dataset.<br><br>Although alternatives are possible for the exact geometry of the coiled-coil domains of TPR, all models adhere to the same basic architectural principle of the | **Human**<br>183 out of 191 crosslinks involving TPR are satisfied.<br>(Fig. 5b; Fig. S9b) | **Human**<br>The overall shape and size of the structural model of the TPR tetramer is consistent with the electron density observed in subtomogram averages of the basket filaments in human macrophages and HEK293 cells (Fig. 1a, 5c, S9a). | Literature: This domain arrangement was previously determined by[28].<br><br>Literature: Parallel homo-dimerization of TPR has been determined[30]. | good |

| | | | | | |
|---|---|---|---|---|---|
| | TPR dimer: four stacked TPR coiled-coils spanning residues 1–390 and 660–1460. | 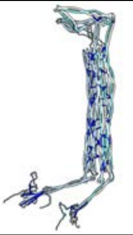 | | | |
| **TPR**<br>C-terminal coiled-coil<br>(aa 1460–1560)<br>disordered region<br>(aa 1560–2363) | **Human**<br>Moderate confidence model of the C-terminal coiled-coil.<br><br>No interaction with the TPR coiled-coil bundle detected by AF3 (Table S4) | XL-MS data do not identify extensive interactions of this part of TPR and other TPR coiled-coils (Table S4). | | | low |
| **ZC3HC1** | **Human**<br>High confidence<br>TPR N-terminal region–TPR insertion region–ZC3HC1<br>(Fig. S2f)<br>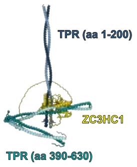 | **Human**<br>Two satisfied crosslinks:<br>-between ZC3HC1 and the TPR N-terminal region:<br>ZC3HC1 89–TPR 47<br>-between ZC3HC1 and the TPR insertion region:<br>ZC3HC1 31–TPR 438<br>(Fig. 5d)<br>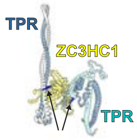 | | Literature: Interaction of TPR with ZC3HC1 in *X. laevis* and HeLa cells [36].<br><br>Literature: ZC3HC1 binds the NPC via two domains: aa 72–290 and 398–467[38].<br><br>Literature: Yeast Pml39 interacts with Mlp1 aa 1–297 and 287–584, and Mlp2 aa 1–210 (Y2H) [118]. | high |
| **NUP153**<br>(aa 250–275)<br>interacting with<br>**NUP107** | **Human**<br>High confidence<br>NUP107–NUP153<br>(Fig. S2e)<br>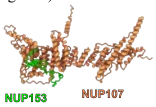 | | **Human**<br>Superposition with the established copies of NUP107 in the NR positions NUP153 such that it can be connected to the neighboring fragment of NUP153 (aa 305–320) interacting with the corresponding copy of TPR.<br><br>***X. laevis***<br>Nup153 is resolved in the density of the cryo-EM map of the NR of the *X. laevis* NPC (EMD-31065) (Fig. S10c)<br><br>Nup107 outer:<br>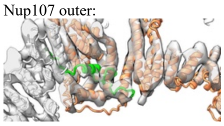<br>Nup107 inner:<br>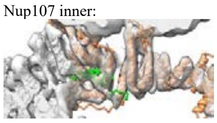 | Literature: Consistent with domain mapping experiments for NUP153 binding to the Y-complex [23]. | high |
| **NUP153**<br>(aa 280–320)<br>interacting with the<br>**inner NUP85** subcomplex<br>(exclusive to the interaction with TPR and NUP107) | **Human**<br>High confidence<br>NUP85–SEH1–NUP43–GANP–NUP153<br>(Fig. S2b)<br>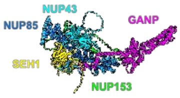<br>***X. laevis***<br>High confidence<br>Nup85–Seh1–Nup43–Nup153–Ganp<br>(Fig. S2i)<br>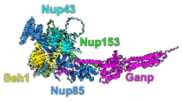 | **Human**<br>*Thirteen satisfied crosslinks between NUP153 and NUP85 or SEH1:<br>NUP153 317–NUP85 92<br>NUP153 326–NUP85 92<br>NUP153 326–NUP85 218<br><br>NUP153 356–NUP85 218<br>NUP153 353–NUP85 218<br>NUP153 353–NUP85 103<br>NUP153 353–SEH1 251<br>NUP153 353–NUP85 92<br>NUP153 356–NUP85 92<br><br>NUP153 371–NUP85 218<br>NUP153 371–NUP85 95<br>NUP153 371–NUP85 92<br>NUP153 371–NUP85 103<br><br>Two satisfied crosslinks between NUP153 and NUP160 or NUP205:<br>NUP153 353–NUP205 892<br>NUP153 371–NUP160 1331 | ***X. laevis***<br>Nup153 is resolved at the interface with the inner Nup85 subcomplex in the density of the cryo-EM map of the NR of the *X. laevis* NPC (EMD-31065)<br>(Fig. S10c)<br>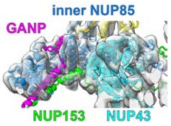 | Literature: Consistent with domain mapping experiments for NUP153 binding to the Y-complex [23]. | high |

| | | | | | |
|---|---|---|---|---|---|
| | | Three slightly violated NUP153–SEH1 crosslinks (likely due to flexibility of NUP153):<br>NUP153 317–SEH1 251<br>NUP153 317–SEH1 255<br>NUP153 326–SEH1 251<br>(Fig. 5f)<br>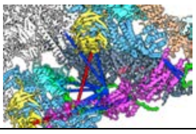 | | | |
| **NUP153**<br>(aa 360–370)<br>interacting with<br>**NUP93** | **Human**<br>High confidence<br>NUP93–NUP153<br>(Fig. S2e)<br>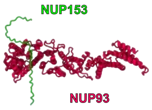 | | **Human**<br>Superposition with the established copy of NUP93 in the NR positions NUP153 such that it can be connected to the neighboring fragment of NUP153 (aa 250–275) interacting with the outer copy of TPR. | | moderate |
| **NUP50** dimer<br>(aa 150–200)<br>interacting with<br>**NUP153** (aa 520–560) | **Human**<br>High confidence<br>NUP50×2–NUP153<br>(Fig. S2e)<br>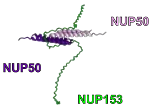 | **Human**<br>Three satisfied crosslinks between NUP50 and NUP153:<br>NUP50 173–NUP153 526<br>NUP50 127–NUP153 526<br>NUP50 62–NUP153 954<br>(Fig. 5e)<br>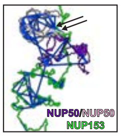 | | Literature: The NUP153 aa 401–609 region is necessary and sufficient for its interaction with NUP50 [62]. | high<br>(but position arbitrary) |
